# Innovation, conservation and repurposing of gene function in plant root cell type development

**DOI:** 10.1101/2020.04.09.017285

**Authors:** Kaisa Kajala, Lidor Shaar-Moshe, G. Alex Mason, Mona Gouran, Joel Rodriguez-Medina, Dorota Kawa, Germain Pauluzzi, Mauricio Reynoso, Alex Canto-Pastor, Vincent Lau, Mariana A. S. Artur, Donnelly A. West, Concepcion Manzano, Sharon B. Gray, Andrew I. Yao, Marko Bajic, Elide Formentin, Niba Nirmal, Alan Rodriguez, Asher Pasha, Alexander T. Borowsky, Roger B. Deal, Daniel Kliebenstein, Torgeir R. Hvidsten, Nicholas J. Provart, Neelima Sinha, Daniel E. Runcie, Julia Bailey-Serres, Siobhan M. Brady

**Affiliations:** Department of Plant Biology and Genome Center, University of California, Davis, Davis, CA 95616 USA; Plant Ecophysiology, Institute of Environmental Biology, Utrecht University, 3584 CH Utrecht, the Netherlands; Plant Biology Graduate Group, University of California, Davis, Davis CA 95616 USA; Integrative Genetics and Genomics Graduate Group, University of California, Davis, Davis CA 95616 USA; Center for Plant Cell Biology, Botany and Plant Sciences Department, University of California, Riverside, Riverside, CA 92521; IBBM, FCE-UNLP CONICET, La Plata, 1900 Argentina; Department of Cell and Systems Biology/Centre for the Analysis of Genome Evolution and Function, University of Toronto, 25 Willcocks St, Toronto, Ontario M5S 3B2 Canada; Department of Plant Biology, University of California, Davis, Davis, CA 95616 USA; Department of Biomedical Engineering and Genome Center, University of California, Davis, Davis, CA 95616 USA; Emory University, Department of Biology, Atlanta, GA 30322; Department of Biology, University of Padova, Padova, Italy; Faculty of Chemistry, Biotechnology and Food Science, Norwegian University of Life Sciences, 1432 Ås, Norway; Department of Plant Sciences, University of California, Davis, Davis, CA 95616 USA

## Abstract

Plant species have evolved myriads of solutions to adapt to dynamic environments, including complex cell type development and regulation. To understand this diversity, we profiled tomato root cell type translatomes and chromatin accessibility. Using xylem differentiation in tomato, relative to Arabidopsis, examples of functional innovation, repurposing and conservation of transcription factors are described. Repurposing and innovation of genes are further observed within an exodermis regulatory network and illustrate its function. Translatome analyses of rice, tomato and Arabidopsis tissues suggest that root meristems are more conserved, and that the functions of constitutively expressed genes are more conserved than those of cell type/tissue-enriched genes. These observations suggest that higher-order properties of cell type and pan-cell type regulation are conserved between plants and animals.

**One Sentence Summary:** Pan-species cell type translatome and chromatin accessibility data reveal novelty, conservation and repurposing of gene function.

## Main Text

*Solanum lycopersicum* (tomato) in the family Solanaceae originated on the west coast of South America. Different species in the family, including tomato, pepper and eggplant, arose and adapted to a variety of habitats from coastal desert to humid rainforests, in part through modifying cell type-specific developmental programs. Irrespective of species, all plant roots contain a stem cell niche at the root tip and cell types along the radial axis are arranged in concentric cylinders. Further, these cell types are constrained within files along the root longitudinal axis. Many of these cell types, including the epidermis, endodermis, xylem and phloem, are hypothesized to be functionally homologous across species. Despite these similarities, there is also diversity in root cell types across plant species as well as in cell signaling and metabolic programs. As an example, the exodermis is an outer cortex layer which produces an apoplastic barrier, and is present in a reported 89% of angiosperms, but absent in the intensely studied model species Arabidopsis (*1*). In rice, multiple cortical cell types exist, and unlike Arabidopsis, some central cortical cells undergo programmed cell death to form gas-filled aerenchyma that promote oxygen diffusion to apical regions of waterlogged roots.

Cell type-resolution and single cell transcriptomes or ribosome-associated mRNA profiles (translatomes) have provided insight into the regulatory mechanisms underlying root development as well as its interaction with the environment in single species (*2, 3*). However, our understanding of how conserved cell type programs are across species relies on morphology, or on annotation via developmental markers identified in a reference species. The degree to which cell type developmental programs are molecularly or functionally conserved across species is, as of yet, unknown. In an effort to address this gap, we map cell type-resolution translatomes of 12 different cell type or tissue populations as marked by distinct promoters in *S. lycopersicum*, cv. M82 (*4*) (**Fig. 1A, S1**). Translatomes are easily accessed by ribosome immunopurification and provide a proxy for translation, eliminating protoplasting-induced changes. Promoters were fused to a FLAG-tagged ribosomal protein (RPL18/eL18) to enable Translating Ribosome Affinity Purification of mRNA coupled with sequencing (TRAP-seq) (**Fig. 1B**) (*4–6*). Principal component analysis confirmed the reproducibility of the marker line-derived translatomes (**Fig. S2A**) and the resulting data can be visualized on a gene-by-gene basis in the ePlant browser (**Fig. 1C**). We identified <u>c</u>ell type-<u>e</u>nriched <u>g</u>enes (CTEGs) that display significant translatome enrichment in one cell type relative to others using limmaVoom (*7*) and ROKU, a Shannon-entropy-based method (*8*) (**Fig. 1D, S2B, Data S1**).

**Fig. 1.**
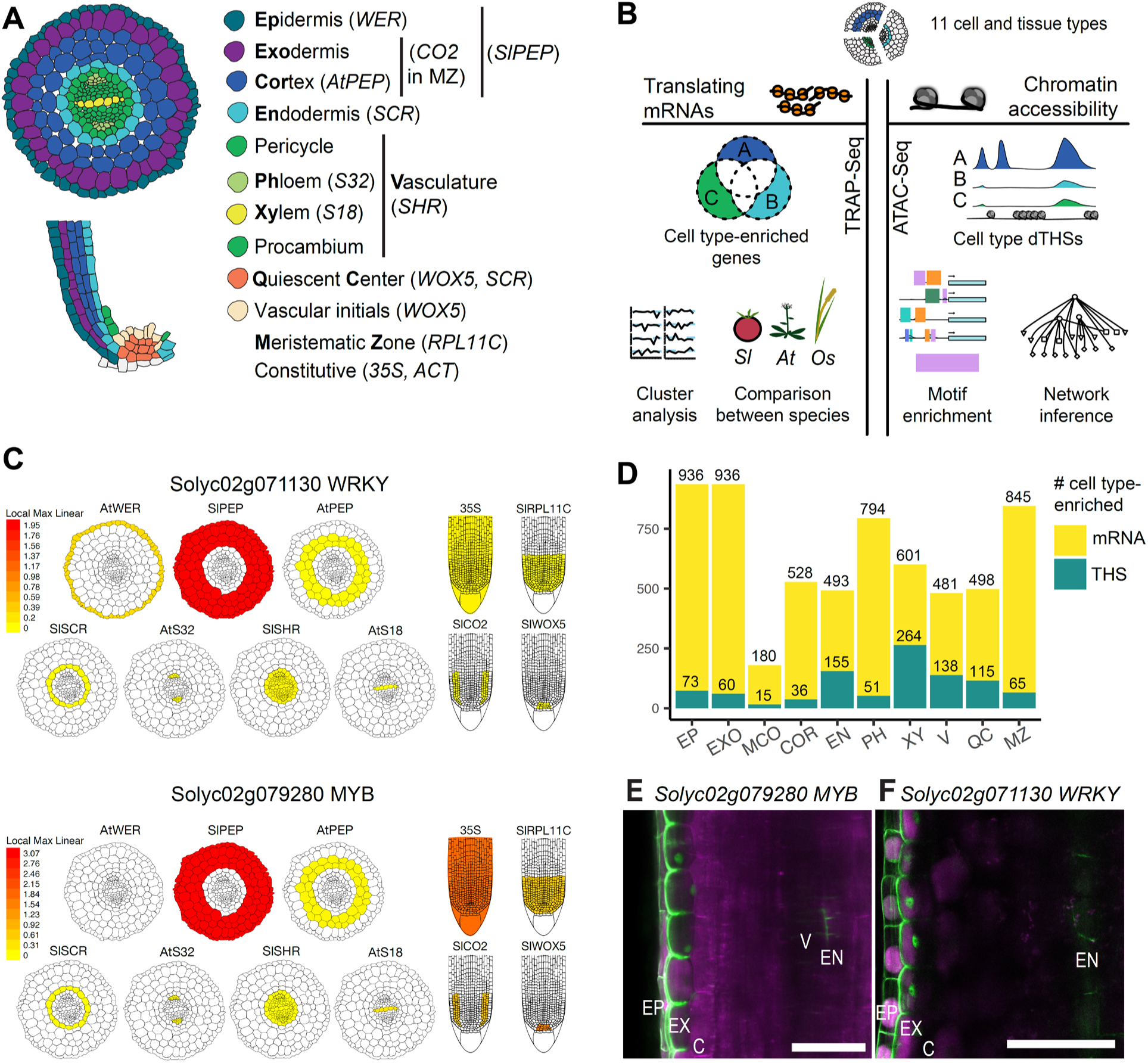
Tomato root atlas of chromatin accessibility and translatome patterns at cell type resolution. (**A**) Tomato root cell types profiled and the respective promoters that mark them. (**B**) Overview of experimental approach. (**C**) Relative TRAP-RNA abundance of *Solyc02g079280* and *Solyc02g71130* at cell type resolution in the Tomato ePlant “Experiment eFP” view. (**D**) Number of cell type enriched THSs and TRAP-RNA transcripts. (**E**) *Solyc02g079280* and (**F**) *Solyc02g71130* were identified to have exodermis-specific TRAP-RNA expression, confirmed by their *promoter:nlsGFP* patterns.

To validate our approach to identify CTEGs, we explored expression conferred by promoters of two genes inferred to be preferentially expressed in the exodermis (**Materials and Methods**), a *MYB* (*Solyc02g079280*) and a *WRKY* (*Solyc02g071130*) transcription factor, fused to GFP (**Fig. 1E-F**). Strong expression was seen in the exodermal layer underlying the epidermis. Gene function enrichment analysis using Gene Ontology (**Data S2**) and Mapman ontologies (**Data S2**) of the exodermis-enriched genes revealed putative exodermal functions. As of yet undescribed at the molecular level due to its absence in Arabidopsis, the exodermal layer was enriched for genes involved in phenylpropanoid and lipid biosynthesis, as well as for nitrogen metabolic activity and hormone metabolism (ethylene, brassinosteroid and jasmonate biosynthesis). These support histochemical observations suggesting the exodermis contains an apoplastic barrier (*9*), and that it may be a nexus for hormone and nutrient metabolism.

To explore conservation and differences in cell type-regulatory networks between tomato and Arabidopsis, we focused on xylem cell regulation as a case study given its extensive molecular characterization in Arabidopsis (*10*). We hypothesized that critical regulators of xylem differentiation would be conserved between Arabidopsis and tomato. Transcriptional regulation determines the final steps of xylem cell differentiation, including the coordinated transcription of secondary cell wall biosynthetic enzymes. The NAC domain transcription factors (TFs) *VND6* and *VND7*, act at the top of this hierarchy and are sufficient to specify Arabidopsis protoxylem and metaxylem differentiation, respectively (*11*). We reasoned that a functional ortholog of *VND6* or *VND7* in tomato would (i) have sequence similarity; (ii) show transcript abundance in the xylem; (iii) when overexpressed be sufficient to drive metaxylem (VND6) or protoxylem (VND7) differentiation (*11*). Using phylogenetic analyses (**Fig. S3A**), we identified two potential tomato orthologs of *AtVND7* (*Solyc11g018660* and *Solyc06g065410*), two putative orthologs of *AtVND6 (Solyc03g083880* and *Solyc06g0343*0) and another tomato gene with some sequence similarity to *AtVND6* and *AtVND7* (*Solyc08g0079120*) (**Fig. S3A**). We used the cell type-resolution translatome data to reduce this list of potential “functional” orthologs to two TFs, *Solyc08g0079120* and *Solyc06g03430*, given their expression in the xylem translatome, as expected of a *VND6/7* ortholog (**Fig. S3A**). Overexpression of only *Solyc06g034340,* and not *Solyc08g079120*, using hairy root transformation (*4*) was sufficient to drive ectopic helical secondary cell wall deposition in all cells within the root, consistent with *AtVND7* function (**Fig. 2A**). Thus, the combination of phylogenomic, translatome, and overexpression phenotypic data suggests that *Solyc06g03430 (SlVND7)* is a functional ortholog of *AtVND7.* Further, these approaches failed to identify a functional ortholog of *AtVND*6, and thereby demonstrate only partial conservation of VND factors between Arabidopsis and tomato.

**Fig. 2.**
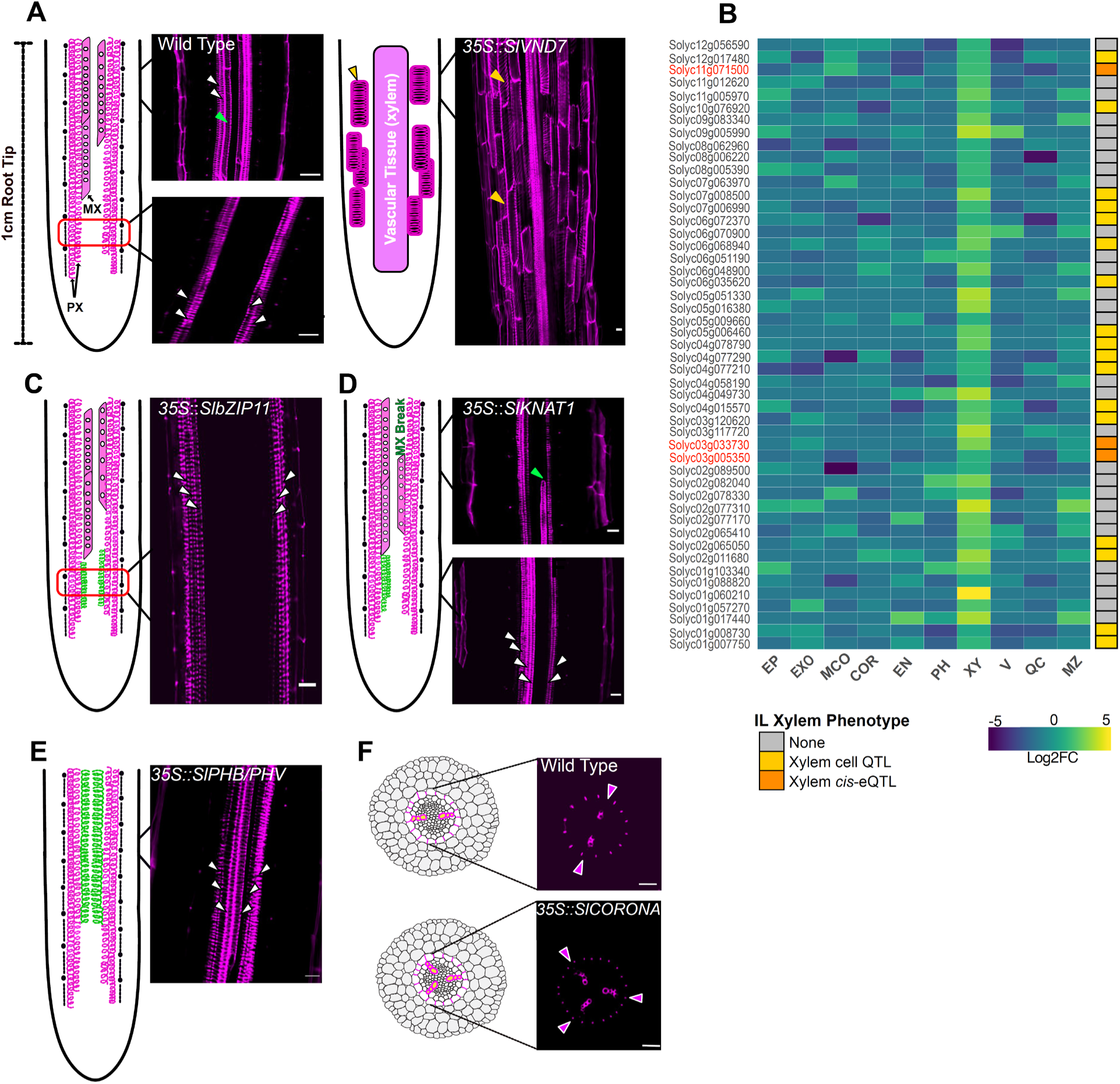
Identification of xylem vessel transcriptional regulators in tomato. (**A, C, D, E, F**) Schematic and confocal images of basic fuchsin-stained roots of wild type and overexpression lines. Fuchsin stains lignin and phenylpropanoid molecules. (**A**) Wild type (left panel) and *35S:SlVND7* (right panel). (**B**) Heatmap of xylem-enriched TFs in tomato roots. In red are genes that were validated in overexpression lines. (**C**) *35S::SlbZIP11*, (**D**) *35S::SlKNAT1* (**E**) *35S::SlPHB/PHV* (**F**) 3*5S::SlCORONA.* White arrowheads indicate protoxylem, green arrowheads indicate metaxylem, yellow arrowheads indicate secondary cell wall deposition in non-xylem cell types and purple arrowheads indicate the xylem axis, PX =protoxylem and MX=metaxylem. Red boxes indicate the region imaged in the image to the right, and contain the endodermis and stele within the distal meristem/elongation zone. Scale bars: 20µm

Conversely, there may be lineage-specific, Solanaceae/Solanum-expanded, or tomato-specific gene duplications that represent innovation or repurposing of regulation in tomato xylem compared to Arabidopsis. In a complementary approach, we used tomato xylem-enriched TFs and associated quantitative genetic data to find novel or conserved regulators of xylem development. Xylem-enriched TFs (**Fig. 2B**) were filtered for genes with *cis*-expression quantitative trait loci located within intervals significantly associated with natural variation in xylem cell number (*12*), (*13*) (**Fig. 2B, right panel**). Five of these 20 enriched TFs could represent lineage- or species-specific gene innovation, including a member of a Solanaceae-specific clade (*Solyc06g072370*), a Solanum-expanded clade (*Solyc11g071500*), a gene duplication in Solanaceae (*Solyc03g005350*) and a tomato gene duplication (*Solyc02g011680, Solyc07g008500*) (**Fig. S4**). To validate this complementary approach and test the functional conservation of these xylem-enriched TFs, we selected six TFs: the Solanum-expanded *MYB* (*Solyc11g071500)* whose distantly related Arabidopsis relative, *At5g23650* has not been functionally characterized and is not expressed in root xylem cells (*2*), a *bHLH* duplicated in the Solanaceae *(Solyc03g005350)*, *SlbZIP11* (*Solyc03g033730)*, whose likely Arabidopsis ortholog (*At4g34590; AtbZIP11*) is not expressed in Arabidopsis root xylem (*2, 14*), the putative *AtKNAT1* ortholog (*Solyc04g077210*), as *AtKNAT1* is a repressor of lignification within Arabidopsis inflorescence stem vascular bundles and is not expressed in primary root xylem (*15*), and two HD-ZIPIII TFs, *SlPHB/PHV (Solyc02g069830)* and *SlCORONA (Solyc03g120910)*, whose Arabidopsis orthologs regulate root protoxylem vessel differentiation via positional signals derived from a miR165/166 gradient (*2, 11, 16*). Contrary to their function in Arabidopsis, over-expression of *SlbZIP11* or *SlKNAT1* was sufficient to specify additional protoxylem cell files (**Fig. 2C-D**), although these files were often non-contiguous for the *SlbZIP11* lines (**Fig. 2C**) (statistical analyses in **Fig. S5, Data S3**). The *bHLH* and *MYB* overexpression lines had no vascular phenotype. Relative to Arabidopsis, In the case of *SlKNAT1*, this demonstrates “repurposed” regulation, while in the case of *SlbZIP11* it represents innovation in function. miRNA-resistant versions of *SlCORONA* and *SlPHB/PHV* were sufficient to regulate protoxylem vessel identity and patterning within the vascular cylinder similar to their Arabidopsis function and are thus conserved regulators (**Fig. 2D, E**).

Cell type/tissue translatomes are likely dynamic over developmental time and in response to the environment. In Arabidopsis, cell type-enriched genes that maintain expression despite stress are also critical regulators of cell fate (*3, 17*). However, the majority of plant cell type profiles are generated from plants grown in sterile culture, leaving a knowledge gap as to how these signatures translate to plants grown in soil. We thus profiled translatomes of the meristematic cortex, endodermis and meristematic zone in two-month-old plants grown in the field under standard cultivation practices as well as of the meristematic cortex and the entire cortex of one-month-old plants grown in pots (**Data S4**). First, we identified “core” CTEGs whose cell type expression was evident across different environmental conditions or plant age, and which, therefore, can be considered reliable cell type markers. By comparing the translatomes of plate- and field-grown plants we identified 47, 2 and 50 core CTEGs for the endodermis, meristematic cortex and meristematic zone, respectively (**Data S5**, **Fig. S6**). Endodermis-enriched core CTEGs included *SCARECROW (SlSCR, Solyc10g074680),* two zinc finger TFs - *Solyc01g090840,* an ortholog of <u>Z</u>INC *F*INGER <u>A</u>RABIDOPSIS <u>T</u>HALIANA <u>G</u>ENEs (*AtZAT4* and *AtZAT9*) and the Solanum zinc finger (C-x8-C-x5-C-x3-H) family protein *Solyc06g054600* (**Data S5**). *SCR* is also considered a core endodermis-enriched gene in Arabidopsis (*3*). Therefore, despite considerable differences in developmental stage and environment, examples of core root CTEGs do exist.

Genes displaying CTE expression dependent on environmental condition or root type, or across ontogenetically related tissues, can reveal context-specific functional specialization. As an example, exodermis cells, inner cortex cells and endodermis cells originate from the same initial cell, yet it is unknown if these cells have any co-expressed genes, and thus common functions (*12*). Such patterns were determined using weighted gene correlation network analysis (WGCNA) (*18*), coupled with ontology enrichment analysis (**Fig. S7**, **Data S4**). Modules with enriched gene expression in (i) the cortex and exodermis of primary roots, and the cortex of lateral roots (**Fig. 3A**), (ii) the cortex of lateral roots and adventitious roots (**Fig. 3B**) and (iii) exodermis-enrichment only (**Fig. 3C**) indicate commonalities and specificities of the exodermis and inner cortex layers, as well as potential conservation of function over root type, environment and developmental stage. Genes in the exodermis module include peroxidases and other enzymes associated with nitrogen metabolism, similar to those identified in the CTEG lists (**Data S4**). The general cortex module includes several TFs, including two GRAS family TFs *Solyc02g092570* and *Solyc03g123400*, orthologs of Arabidopsis *PUTATIVE SCARECROW-LIKE PROTEIN 16* and *SCARECROW-LIKE PROTEIN 29*, respectively, and genes associated with lipid metabolism (fatty acid synthesis and elongation), ADP binding and ATPase activity (**Data S4**). Genes enriched in the cortex of lateral and adventitious roots are associated with calcium signaling and hydrolase activity (**Data S4**). A large module with genes uniquely translated in the endodermis of older plants (2 months) under typical field cultivation conditions, but not in seedlings grown on agar plates, (**Figure 3D**) includes genes associated with nucleic acid binding and protein dimerization activity **Data S4**). There was no module showing relatively high expression in the exodermis, inner cortex and endodermis, indicating that, despite a common progenitor cell for these tissues, their cell type functions are divergent. These modules underscore the importance of profiling cell type resolution gene expression across diverse growth conditions.

**Fig. 3.**
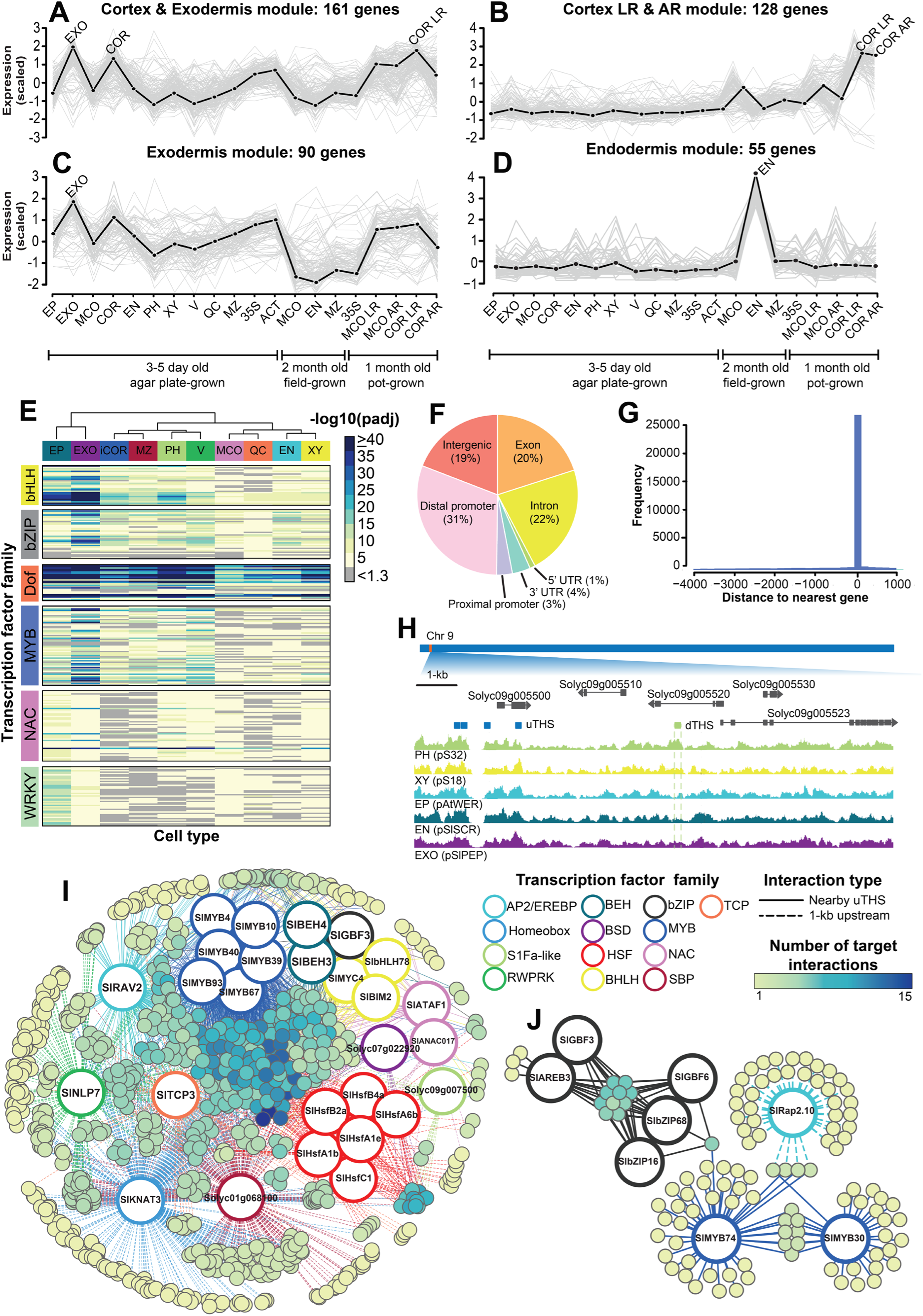
Core and variable cell type-enriched genes and inferred unique cell type regulatory networks. (**A**) Cortex and endodermis module. (**B)** Cortex lateral root and adventitious root module. (**C**) Exodermis module. (**D**) Endodermis field module. (**A-D**) WGCNA co-expression modules with scaled expression values (y-axis) across translatome profiles derived from different promoters and conditions. Three-five day old plants grown on agar plates in a growth chamber: EP=epidermis; EXO=exodermis; MCO=meristematic cortex; COR=general cortex; EN=endodermis; PH=phloem; XY=xylem; V=vasculature; QC=quiescent center; MZ=meristematic zone; 35S=constitutive promoter a; ACT=constitutive promoter 2. Two month old plants grown in the field: MCO, MZ, END and 35S as for the 5-7 day old plants. One month old plants grown in the growth chamber; MCO and COR as for the 5-7 day old plants; LR=lateral roots; AR=adventitious roots. Black dotted line = eigengene expression profile. The maximum peak enrichment within the module is indicated by black font on top of the eigengene expression line. Grey line = expression values of all genes within the module. Most of the genes in these modules were positively correlated to the eigengene. (**E**) Transcription factor motif enrichment for selected transcription factor families (y-axis) within 1kb promoter regions of CTEGs (x-axis). Adjusted p-values (-log_10_) are indicated in the heat scale. (**F**) Genomic location of uTHSs. Proximal promoter = 500 bp upstream of transcription start site. Distal promoter = Between 500 bp and 4kb upstream of the transcription start site. (**G**) The frequency (y-axis) of uTHSs relative to the distance to the nearest gene (**Materials and Methods**). (**H**) Representative phloem cell type-enriched accessible region (CTEAR). Y-axis = merged cut counts normalized with deeptools (**Materials and Methods**); x-axis = chromosome location; Gene models = top track; uTHSs are indicated with a blue rectangle while a CTEAR is indicated with a green rectangle. PH= phloem; XY= xylem; EP=epidermis; EN=endodermis; EXO=exodermis. (**I**) Unique inferred exodermis regulatory network. (**J**) Unique inferred inner cortex regulatory network. (**I-J**) Solid edges = motif-uTHS interaction; dashed edges = motif-1kb upstream regulatory region interaction; large circles = TF expressolog for cognate TF motif; colored edges indicate transcription factor family. Small circles = exodermis-enriched target genes which contain the motif in either the uTHS or 1kb upstream regulatory region; color scale indicates the number of target interactions.

TFs are critical regulators of cell type development by spatially restricting gene expression, and differences and similarities in such regulators are hypothesized to exist between Arabidopsis and tomato. Hence, we surveyed the promoters of CTEGs for enriched *cis*-regulatory motifs. WRKY and bHLH TFs are known to regulate Arabidopsis epidermal cell fate (*19, 20*), and we correspondingly found WRKY and bHLH TF binding sites enriched in the promoters (1-kb upstream of transcription start site) of genes expressed in the tomato epidermis (**Fig. 3E, S8**). While 9 NAC domain TF-binding sites are over-represented in xylem-enriched promoters, as a whole, the NAC-domain TF binding sites do not show specificity to the xylem. This supports the limited conservation of VND regulators between Arabidopsis and tomato (**Fig. 2A, S3**). MYB regulation in xylem cells is also a critical component of Arabidopsis xylem development (*21*), and MYB domain binding sites are significantly over-represented in xylem-enriched genes, demonstrating likely similarity in MYB regulation of xylem development between Arabidopsis and tomato. Highly significant MYB and bHLH binding site enrichment in the exodermis-enriched genes suggest that these factors are important in exodermis specification or function. Another difference between Arabidopsis and tomato is that of DOF domain TFs, which are influential in vascular patterning in Arabidopsis (*22*), but which do not show unique cell type-enrichment in tomato. Collectively, these cell type-enriched motifs suggest both similarity and divergence of transcription factor-mediated regulation of cell type development.

Chromatin availability assays, including Assay for Transposase Accessible Chromatin (ATAC) coupled with sequencing, provide further insight into mechanisms producing cell type-specific transcriptomes. In Arabidopsis root hair and non-hair cells, the majority of Transposase Hypersensitive Sites (THSs) were shared between epidermal cell types, although differential THSs were also present (*23*). We hypothesized that these properties of largely common THSs and smaller numbers of cell type-restricted THSs would not be species-specific, and that this observation would extend across multiple cell type comparisons in tomato. Thus, 11 cell type/tissue promoters were used to drive a fusion gene encoding a biotin ligase receptor peptide, GFP, and a nuclear envelope protein, along with biotin ligase driven from a constitutive promoter for Isolation of Nuclei TAgged in specific Cell Types to enable ATAC-seq (**Fig. 1B**) (*4, 24*). 108,335 reproducible transposase hypersensitive sites (THSs) were identified across cell types (**Fig. S2B**), with half found in intergenic regions distal to the transcription start site (TSS) as previously described (*23*) (**Methods, Supplemental Dataset 1, Fig. 3F**). 1,774 THSs differ in their accessibility between at least two cell types, and 972 THSs were enriched in a given cell type and are called <u>C</u>ell <u>T</u>ype-<u>E</u>nriched <u>A</u>ccessible <u>R</u>egions (CTEARs) (**Fig. 1D, Data 6**). Union THSs (<u>u</u>THSs) were annotated to their nearest gene, with the vast majority found within genic regions (**Figure 3G**). As shown in animals, CTEARs overlap with cell type expression (*25*). In tomato, CTEARs are associated with genes more likely to be expressed in the target cell type than expected by chance (**Fig. S9**). A CTEAR (**Fig. 3H**) for a phloem-enriched accessible region, and other select CTEARs are demonstrated in **Fig. S10**. Motif enrichment of uTHSs showed similar patterns to those described for promoters of CTEGs (**Fig. S11**). Taken together, these properties of largely common THSs, smaller numbers of cell type-restricted THSs and overlap with cell type gene expression extends between Arabidopsis and tomato.

Motif enrichment within open regulatory regions and promoters of CTEGs provide an excellent opportunity to infer cell type regulatory networks. Such analyses have revealed both conservation and flexibility of networks in response to submergence in several species (*6*). We identified TF motifs from target gene promoters and nearby accessible regions (*6*), and the most likely tomato ortholog of the motif’s cognate Arabidopsis TF. Putative “functional” orthologs of the TFs (*i.e.* expressologs **Materials and Methods**), were determined by coupling sequence homology (Phytozome gene families) with expression correlation within homologous cell type/tissues in tomato and Arabidopsis (*i.e.* root TRAP-expressologs, **Materials and Methods**). Finally, TF expressologs were included in the network if they were expressed in the translatome of a given cell type (**Materials and Methods**). Combining these data, we inferred unique regulatory networks for each tomato root cell type (**Materials and Methods**, **Fig. 3I,J, S12, Data S7**). These networks were particularly informative for the as yet molecularly uncharacterized exodermis and inner cortex layers.

The expressologs of *AtMYB4* and *AtMYB39*, which are associated with lignin and suberin metabolism in the Arabidopsis root endodermis, are instead associated with lignin and suberin metabolism in the exodermis network. This suggests potential co-option of this regulon to the exodermis (*26*). Several Heat Shock Factors (HSFs) were also prevalent in the exodermis network and are predicted to coordinately bind promoters of genes associated with auxin transport, ethylene biosynthesis, phosphate transcriptional regulation and lignin biosynthesis or polymerization (**Fig. S13**). There was also a link with nitrogen transcriptional regulators in the exodermis. Tomato expressologs of *AtNLP7, AtRAV2* and *AtTCP3* were predicted to bind to promoters of genes associated with lignin polymerization, nitrate or amino acid transporters, and a nitrate reductase (**Data S7**). Of these three Arabidopsis genes, only *AtRAV2* is expressed in the cortex. Thus, for *AtNLP7* and *AtTCP3*, this represents a change in their expression domain. Support of this observation of nitrogen transcriptional regulators embedded within the exodermis network is provided by more Arabidopsis nitrogen response regulatory network genes having expressologs within the exodermis network (*27*) than expected by chance (odds ratio = 3.0, *p* < 0.01) **(Fig. S14)**. Cell type- and tissue-specific nitrogen responses have been mapped in the Arabidopsis root, with the pericycle and lateral root cap being the most transcriptionally responsive (*28*). This unique exodermis-enrichment demonstrates likely differences in nutrient signaling between Arabidopsis and tomato.

The inner cortex regulatory network **(Fig. 3J)** has a prevalence of TFs whose most likely Arabidopsis ortholog is involved in dehydration or ABA-response (AREB3, RAP2.10). These include a bZIP/GBF/AREB module whose common module targets include several TFs, transporters and a homolog of the PYR/PYL ABA receptor; and a MYB/RAP module, whose common module targets include three metabolic genes and an AREBP transcription factor (**Data S7**). Limited functions are known for inner cortex layers, and these data suggest they are a hub for ABA signaling and metabolism. Of particular note, the GBF3 TF binding site, in Arabidopsis, differs depending on DNA methylation (*29*). In this instance, the methylated site was enriched in the exodermis-unique network, while the unmethylated site was enriched in the inner cortex-unique network. Exodermis and inner cortex cells are derived from a common initial cell, and future studies should explore if methylation plays a role in the specification of the exodermis and inner cortex cell type identity.

The pan-developmental and -environmental stage comparisons within tomato (**Fig. 3A-D**) provide insight into both the variability and robustness in gene expression within individual cell types or tissues. We next set out to explore this phenomenon in potentially homologous tissues in diverse plant species. Comparative transcriptome studies of homologous tissues in vertebrates demonstrate that gene expression data tend to cluster by homologous tissue rather than by species (*30*) supporting the hypothesis that conserved gene regulatory networks drive homologous cell type/tissue identity across species and suggesting functional equivalency of these tissues. The similarity in root cell type patterns and the primary function of roots in water acquisition, mineral transport and anchoring of the plant suggest a similar phenomenon in plants. A conserved and environmentally flexible response to submergence by root tips was identified in divergent species - from rice to tomato, which diverged over 250 million years ago (*6*). Yet, the functional equivalency of plant tissues and cell types is largely unexplored.

Here, we sought to determine the degree to which tissue expression conservation observed in vertebrates extend to plants (**Fig. S15**). We generated and collected translatome profiles of the meristematic cortex, endodermis, vasculature and meristematic zone of tomato, Arabidopsis (*5*) and rice (**Materials and Methods, Data S8**) as marked by similar promoters (**Fig. 4A**, **S16**). To explore translatome similarities we focused on 1:1:1 orthologs (**Materials and Methods, Data S8**), since duplicated genes rapidly lose their expression similarity (*31*). PCA analysis shows that the translatome profiles of the meristematic zone from all three species group together and are distinct from the other tissues (**Fig. 4B**) and this was supported for two additional independently derived orthology maps (**Materials and Methods**, **Fig. S17B and C**). In addition, similarities between Arabidopsis and tomato endodermis and vasculature are supported by some of these orthology maps (**Fig. 4B, S17B and C**).

**Fig. 4.**
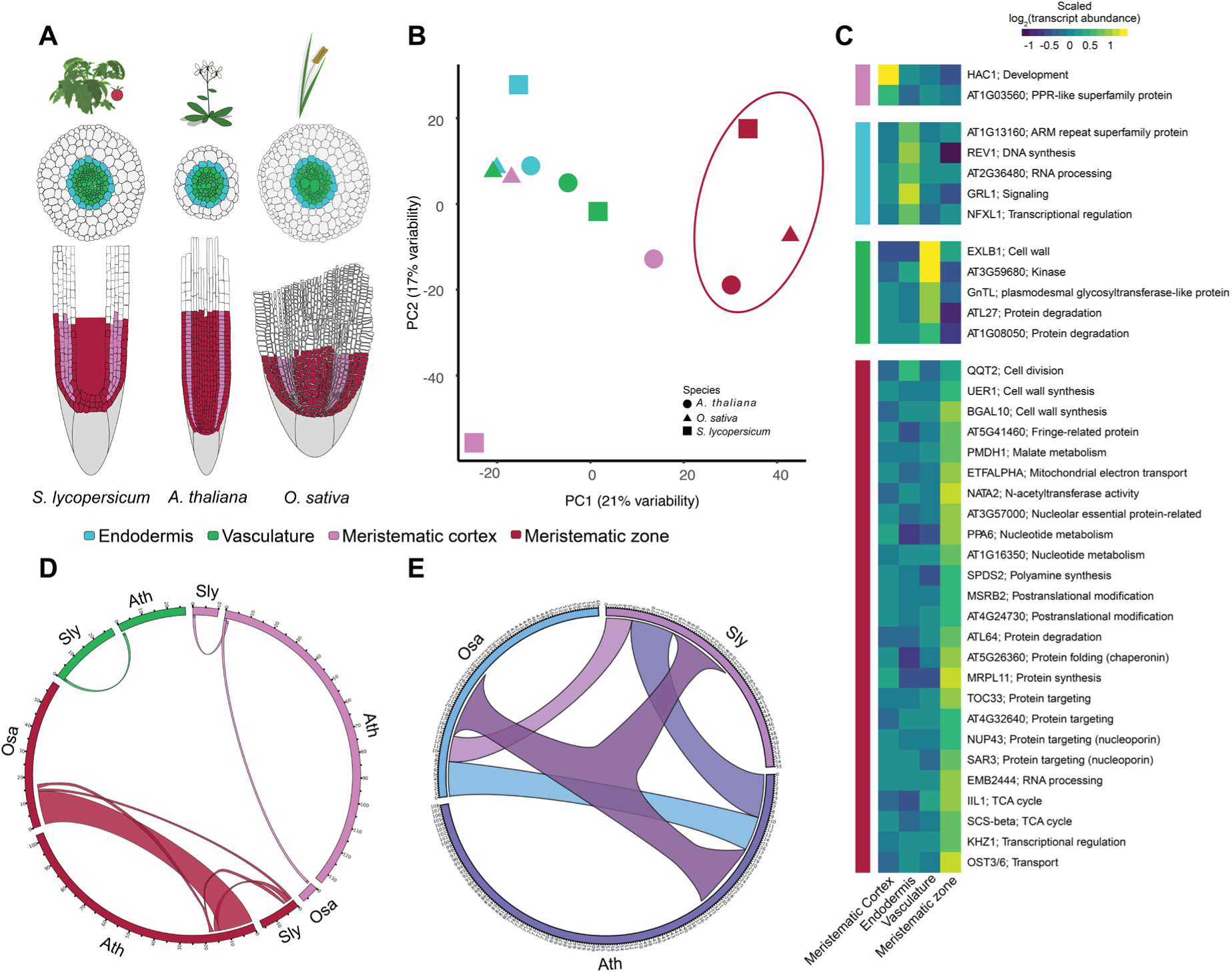
Homologous cell types and tissues show limited conservation of gene expression. (**A**) Species and cell types selected for comparative translatome analysis. Colors in legend are used throughout Figure 4. (**B**) Grouping of cell type/tissue expression profiles between Arabidopsis (circle), rice (triangle) and tomato (square). Colors indicate cell types/tissues as in (**A**). Plot of principal component (PC) analysis of cell type/tissue expression of 2,642 1:1:1 orthologs. Orthology was determined based on Phytozome gene families (**Materials and Methods**). (**C**) Thirty seven conserved cell type- and tissue-enriched expressologs. The mean expression of each consensus expressolog in tomato, Arabidopsis and rice is presented for each cell type/tissue. Transcript abundance is scaled across the four cell types/tissues. The orthology of the consensus expressologs was deciphered based on sequence homology coupled with positive expression correlation, independent of the reference species (**Materials and Methods**). EN=endodermis; MCO=meristematic cortex; MZ=meristematic zone; V=vasculature (**D**) Overlap of MapMan ontology terms between homologous cell types/tissues. The width of the ribbon is proportional to the number of common ontology terms. Ath=*Arabidopsis thaliana*; Osa=*Oryza sativa*: Sly=*Solanum lycopersicum*. Colors are as in (**A**). The numbers in the inner circle represent the number of terms within each group. (**E**) Overlaps of MapMan ontology terms for constitutively expressed genes (CEGs). Color palette is chosen to maximally differentiate pairwise comparisons between species, and three-way overlap is shown in dark purple.

To identify genes with conserved cell type/tissue enriched expression among the three species, we generated a list of root TRAP-expressologs (**Materials and Methods, Fig. S18**). Using a refined list of 1,555 “consensus expressologs” and an ANOVA approach with a Tukey post-hoc test (**Materials and Methods**), we detected 37 expressologs with cell type/tissue-enriched expression conserved across species (**Materials and Methods**, **Fig. 4C, Data S8,** *F*-value>6.6, p-value<0.1). In concordance with the clustering analysis, most of these genes show specific expression in the meristematic zone. Among these genes is *QQT2*, which is essential for correct cell divisions during embryogenesis (*32*) and required for the assembly of RNA polymerases II, IV, and V (*33*). Additional conserved meristematic zone-enriched genes encode two nucleoporins, SAR3 and NUP43, subunits of the nuclear pore complex, which regulate nucleocytoplasmic transport of protein and RNA and play important roles in hormone signaling and developmental processes (*34*). Genes associated with tricarboxylic acid metabolism and cell wall biogenesis are also enriched in the meristematic zone. The *GLR1.1* glutamate receptor is enriched in the endodermis. Given the limitations of orthology relationships and resulting gene annotation, these conserved, cell type/tissue-enriched genes provide an avenue for gene discovery with respect to cell type-specific function.

The two ortholog maps (i.e. 1:1:1 orthologs and consensus expressologs) limit the number of genes that could be assessed by requiring conservation among all three species. We therefore also identified CTEGs within each species (**Data S8**) and assessed their functional similarity (**Materials and Methods**). The similarity observed in the overall translatomes of the meristematic zone (**Fig. 4B**) and the number of conserved meristematic zone-enriched genes (**Fig. 4C**) was further reflected in the relatively high overlap of meristematic zone-enriched ontology terms (**Fig. 4D, S19, Methods**) across species. The lower similarity of the endodermis, vasculature and meristematic cortex (**Fig. 4B**) was also reflected in the limited gene function overlap (**Fig. S19, S20, Data S8, Methods**) between their respective homologous cell types/tissues. Across these four cell types/tissues therefore, the meristematic zone is homologous, while CTEGs and functions of the other cell types/tissues are largely species-specific. Similar observations have been made in animals, where embryonic tissues or early developmental stages of homologous cell types show higher similarity across species than mature cell types/tissues (*35*). The meristematic zone of all three species is enriched with leucine rich repeat receptor kinases (LRR-RKs, **Data S8**), shown to regulate diverse signal transduction pathways including root development. The meristematic zone of tomato and rice shows enriched expression of *RGF1 INSENSITIVE 5* (*RGI5*, **Data S8**), a receptor of Root Meristem Growth Factor 1 in Arabidopsis and which together with additional LRR receptor-like kinases is essential for meristem development (*36*). Despite its meristematic characteristics, the meristematic cortex demonstrates few to no overlaps of enriched terms and expressologs (**Fig. 4D**, **S19, S20, Data S8**). One explanation for this finding is the variable number of cortical cell files and their identity in each species. The Arabidopsis cortex consists of a single cell layer, while in tomato (cv. M82) it consists of three (including the exodermis), and in rice, over ten layers have been reported (*37*) (**Fig. 4A**). Alternatively, the limited similarity can arise from the low number of meristematic cortex-enriched genes in tomato and rice.

A comparative transcriptome study in mammals demonstrated that genes with low expression variation across tissues are enriched for housekeeping genes (*38*), which tend to evolve more slowly than tissue-specific genes (*39*). To test if this observation is also true for plants, that is, that genes with low expression variation (constitutively expressed genes) have housekeeping function, we identified a set of genes with minimal expression variation within each species, referred to as constitutively expressed genes (CEGs) (**Methods, Data S8**). In concordance with the literature, overlapping ontology terms and expressologs between the CEGs are involved in housekeeping functions (e.g., cell division, chromatin remodeling, RNA binding and protein metabolism) (**Fig. S21, Data S8**). In addition, a larger number of ontology terms overlap between the CEGs (**Fig. 4E**) compared with the CTEGs, even when considering only the meristematic zone-enriched genes (odds ratio = 1.9, *p* < 0.03), suggesting that the expression patterns of CTEGs are more affected by speciation than CEGs.

Plant roots are morphologically simple with similar radial and longitudinal cell type organization, and many morphologically similar cell types. Our data demonstrates that this simplicity arises from complex regulation both within a single species and between multiple species. Our integration of multiple cell type-resolution data types shed light not only on how cell type molecular signatures in a single species change over time and between *in vitro* culture relative to field cultivated conditions, but also into the function of cell types with limited molecular characterization, like the exodermis and inner cortex cells. Gene-by-gene functional validation of putative xylem cell regulators revealed examples of conservation (HD-ZIPIII TFs), partial conservation (VND TFs), repurposing (bZIP11 function) and co-option (KNAT1 function)) between Arabidopsis and tomato. Identification of Solanaceae-specific and Solanum-expanded TFs, as well as local TF duplications within tomato, where specific members show enriched xylem expression, provides a rich resource of evolutionary variation to explore from a functional perspective. While limited due to lineage-specific gene family expansion, sub- and neo-functionalization, our multi-species analyses confirm that translation of research between Arabidopsis and other dicots or monocots is not straightforward. Extensive similarity was observed by multiple analyses of the root meristem across divergent species, and we propose that the meristematic developmental stage represents a phylotypic developmental period for plants, that is, the period at which organisms within a common phylum show the maximum degree of morphological similarity (*40*). Given the differences in translatome between the endodermis and vascular tissues across species, such phylotypic homology cannot be ascribed for these cell types. Similar observations between plants and animals were made for early developmental stages as well as for constitutively expressed genes, and suggest that although animals and plants have different common ancestors, at the molecular level, higher-order organizational properties that regulate development are indeed the same. These data and resources serve as powerful tools for evaluating cell type- and single cell processes relevant to breeding stress resilient crops where such applications are limited.

## Acknowledgments

We would like to thank Helen Masson, Reina Marie Sanz, Ayumi Gothberg, Steven Vitales, Jiawen Zheng, Kristina Zumstein and Barbara Waring for experimental assistance, plant maintenance and harvest. We would also like to thank Kerry Bubb for advice on ATAC-seq data analysis. We most sincerely appreciate the assistance of Philip Benfey, Niko Geldner and John Harada for critiquing the manuscript

## Author contributions

Conceptualization - K.K, L.S-M, G.A.M, G.P, M.R, D.W, S.B.G, N.S, D.E.R, J.B-S, S.M.B; Data curation - K.K, L.S-M, G.A.M, J.R-M, D.K, G.P, M.R, A.C-P; Formal analysis - K.K, L.S-M, G.A.M, M.G, J.R-M, D.K, A.C-P, V.L, A.T.B, D.J.K, T.H, N.S, D.E.R; Funding acquisition - K.K, L.S-M, G.A.M, R.B.D, N.J.P, N.S, J.B-S, S.M.B; Investigation - K.K, L.S-M, G.A.M, M.G, J.R-M, D.K, G.P, M.R, V.L, M.A.S.A, D.W, C.M, S.B.G, A.I.Y, M.B E.F, N.N, A.R, A.T.B, S.M.B; Methodology - K.K, L.S-M, G.A.M, J.R-M, G.P, M.R, M.B, D.E.R, S.B.B; Project administration - R.B.D, N.S, J.B-S, S.M.B; Software - L.S-M, G.A.M, J.R-M, A.C-P, V.L, T.H, N.J.P, D.E.R; Supervision - K.K, L.S-M, G.A.M, J.R-M, C.M, R.B.D, T.H, N.J.P, N.S, D.E.R, J.B-S, S.M.B; Validation - K.K, L.S-M, G.A.M, M.G, J.R-M, D.K, G.P, M.R, A.C-P, M.A.S.A, D.E.R, S.M.B; Visualization - K.K, L.S-M, G.A.M, M.G, J.R-M, D.K, G.P, M.R, V.L, M.A.S.A, D.W, C.M, N.J.P, S.M.B; Writing - original draft - K.K, L.S-M, G.A.M, M.G, J.R-M, D.K, M.A.S.A, S.M.B; Writing - review and editing - K.K, L.S-M, G.A.M, M.G, J.R-M, D.K, M.R, A.C-P, M.A.S.A, D.W, C.M, A.T.B, R.B.D, D.K, N.S, D.E.R, J.B-S, S.M.B

## Data and materials availability

Transgenic materials available upon request. Raw sequencing and processed data for expression and chromatin accessibility analyses will be available upon a pending GEO Accession number.

## Supplementary Materials

### Materials and Methods

#### Plant material and growth conditions

Transgenic INTACT and TRAP marker lines of *Solanum lycopersicum* cultivar M82 (LA3475) were generated by *Agrobacterium tumefaciens* transformation at the UC Davis Plant Transformation Facility. The pK7WG-TRAP and pK7WG-INTACT-Sl binary vectors (*4*), https://gateway.psb.ugent.be/search) were used with a range of promoters to drive the expression of either the nuclear tagging fusion (*WPP-GFP-BLRP*) for INTACT or the polysome tag (*His6-FLAG-RPL18-GFP*) for TRAP. The promoters used were the previously published *SlACT2p, 35Sp, SlRPL11Cp, AtWERp, AtPEPp, SlCO2p, SlSCR, SlSHR, SlWOX5, At18Sp* and *At32Sp* (*4*), and *SlPEPp* (*Solyc04g076190*) amplified using CACCTTCTCCAACAACGTAGAAGCTCCTCGCT and GGTGTGCTTTTTCCTTATCAACAAC. The promoters were recombined into pENTR-D/TOPO (Invitrogen) and introduced into pK7WG-TRAP and pK7WG-INTACT-Sl vectors using LR Clonase II Enzyme mix (Invitrogen). In order to visually confirm cell type specificity, the expression patterns of all the promoters driving the GFP-containing INTACT and TRAP tags in tomato (**Figure S1, Figure S16, Data S1**) were imaged using an LSM 700 laser scanning microscope (Carl Zeiss) with the following settings: 488-nm excitation laser, the preset eGFP emission spectrum, 70% laser power, 1.87-Airy unit pinhole and gain optimised to the signal strength (450-1200). Additionally, the 561-nm laser and the preset RFP emission spectrum were used to capture autofluorescence. Rice TRAP lines were imaged using a Leica SP5 laser scanning microsope (Leica) with 488-nm excitation laser at 50% power, 56.7 μm pinhole, the preset eGFP emission, and Smart Gain 650-1100. Additionally, brightfield images were captured to show localization of GFP within the root.

The nuclear and translating ribosome affinity purification experiments were conducted with T1 seed stocks (and T2 as needed) from one independent line per construct (line IDs listed in **Data S1**). Plate-based experiments were conducted with four independent replicates of each line, and for each replicate, 1 cm of primary root tips were pooled from up to 200 seedlings. The seeds were surface sterilized with 3% hypochlorite (Clorox) for 20 minutes and rinsed three times with sterile water. Seven seeds were planted per 12 cm x 12 cm square plate containing 1x MS without vitamins (Caisson), 1% (w/v) sucrose, 0.5 g/L MES, pH = 5.8 and 1% (w/v) agar (Difco). Plates were placed vertically into racks using a completely random design in a growth chamber with a 16:8 light:dark cycle at 25°C and 50%–75% humidity with a light intensity of 55-75 μE. As tomato germination is uneven, the germination day of each seedling was scored and 1 cm of root tip was harvested from 3-5 days after germination. The tissue was harvested at relative noon and placed immediately into liquid nitrogen.

Experiments with 1-month-old plants were conducted as follows. Transgenic seeds (**Data S4**) were surface sterilized and germinated on 1xMS media as described above, with the addition of 200µg/ml kanamycin to screen for the presence of the transgenic construct. After 7 days, seedlings were transplanted into pots with Turface Athletic Profile Field & Fairway clay substrate (Turface Athletics) that was pre-wetted with a nutrient water solution containing 4% nitrogen, 18% phosphoric acid, and 38% soluble potash. Plants were grown in a completely randomized design for 31 days in a Conviron Growth Chamber at 22°C, 70% RH, 16/8 hour light/dark cycle and light intensity of 150-200 µmol/m^2^/s. The root systems were harvested as close to relative noon as feasible (±2h) by immersing the pot into cool water, massaging the rootball free, rinsing three times sequentially with water, and then dissecting the root tissues and flash-freezing with liquid nitrogen. The harvested tissues were the lateral roots at the depth of 6-12 cm, and the adventitious (hypocotyl-derived) roots.

Tomato plants were grown in the field as follows: transgenic seeds (**Data S4**) were surface sterilized and germinated on 1xMS media as described above, and the root tips were dissected for screening for the correct GFP pattern. The remaining seedlings were transplanted on soil and grown in a growth chamber with a 16:8 light:dark cycle at 25°C and 50%–75% humidity with a light intensity of 55-75 μE for one week. The plants were then transferred into a screen house for two weeks prior to transplanting into the field in Davis, California, USA on August 25, 2016 in a randomized block design with six replicate blocks, each block consisting of five plants of each genotype. Plants were grown in the field for 32 days with furrow irrigation once weekly and biweekly removal of flower buds to follow the local genetic modification guidelines. The root systems were harvested by digging the plant out, immersing the root ball with soil into water, massaging the rootball free, and three sequential water rinses prior to flash-freezing the entire root ball with liquid nitrogen.

Transgenic marker lines of rice (*Oryza sativa* cv. Nipponbare) were generated by *Agrobacterium tumefaciens* transformation as described by (*41*) or at the UC Davis Plant Transformation Facility. The Rice TRAP binary vector was constructed as described by (*4*) using the gateway binary vector pH7WG, for hygromycin resistance, as a backbone instead of pK7WG (https://gateway.psb.ugent.be/search) and incorporating rice *OsRPL18-2* as described in (*42*) Promoters were incorporated by LR recombination as performed for *S.lycopersicum* constructs to drive the expression *His6-FLAG-RPL18-GFP* for TRAP. The promoters used were the previously published *35Sp*(*4*)*, AtSCRp* (*5*)*, RSS1p* (*43*), as well as *OsCMZp (Os01g0957100)* and *OsSHR1p (Os07g0586900)* that were amplified using primers described in **Data S10**.

Rice (*Oryza sativa* cv. Nipponbare) seeds from transgenic lines (**Data S8**) were dehulled and surface sterilized in 50% (v/v) bleach solution for 30 min and then rinsed with sterile distilled water. Seedlings were grown on plates (10 cm x 10 cm) containing half-strength Murashige and Skoog standard medium (MS) agar (1% w/v) and 1% w/v sucrose, for 7 days in a growth chamber (16 h day / 8 h night; at 28°C/25°C day/night; 110 μEm-2s-1).The whole root system was placed immediately into liquid nitrogen upon harvesting.

#### Nuclei purification by INTACT for ATAC-seq

These steps were conducted as described in (*6*). In brief, cell type-specific nuclei were isolated from the frozen root tip material using INTACT (*23, 24, 44*), and the nuclei were counted and used for ATAC-seq library preparation (*23*). Libraries were size selected for under 750nt and up to 24 barcoded libraries were pooled together. ATAC-seq libraries were sequenced on the NextSeq 500 at the UC Davis DNA Technologies Core to obtain 40-bp paired-end reads.

#### Ribosome-associated mRNA purification by TRAP and RNA-seq library construction

These steps were conducted as described in (*6*). In brief, cell type-specific ribosome-associated mRNAs were isolated from the frozen root tip material using TRAP (*4, 6, 42, 44, 45*) and mRNA was isolated from the ribosome complexes for non-strand specific random primer-primed RNA-seq library construction (*46*). Barcoded libraries were pooled together and sequenced on the Illumina HiSeq 4000 at the UC Davis DNA Technologies Core to obtain 50-bp reads.

#### RNA-seq data processing and analysis

Barcoded libraries were pooled together and sequenced on the Illumina HiSeq 4000 at the UC Davis DNA Technologies Core to obtain 50-bp reads. Sequences were pooled, and then trimmed and filtered using Trim Galore! (v0.4.5) (*47*) with parameter -a GATCGGAAGAGCACA. Trimmed reads were pseudo-aligned to ITAG3.2 transcriptome (cDNA) (*48*) using Kallisto (v0.43.1) (*49*), with the parameters -b 100 --single -l 200 -s 30, to obtain count estimates and transcript per million (TPM) values.

#### Tomato RNA-seq quality control and differential expression

Raw RNA-seq read counts were filtered to remove genes with zero counts across all samples. Reads were converted to count per million (CPM) using the cpm() function in edgeR. Genes with CPM > 0.5 in at least 4 biological replicates were kept, thus removing genes that were consistently lowly expressed across all samples. In order to perform data quality control, we conducted exploratory data analysis. The data were log_2_ transformed with a prior count of 3 to reduce the contribution of low-abundance genes. Batch effects due to sequencing date were corrected with the removeBatchEffect function (*50*). Similarities and dissimilarities between samples were assessed with principal component analysis (PCA) using the function ‘prcomp’. PCA plots were generated with the ggplot2 package (*51*) (**Fig. S2**). Differentially expressed genes (DEGs) were detected with the limma R package (*50*). CPM values were normalized with the voom function (*7*) using quantile normalization with a design matrix that included identifiers for the cell types and the sequencing replicates (batch). The functions lmfit, contrasts.fit, and ebayes were used to fit a linear model and calculate differential gene expression between the different contrasts. Genes with a log_2_ fold change (FC) value ≥ 2 and adjusted P-value (adj.P.Val) ≤ 0.05 were considered as differentially expressed. The fdr method was used to control the false discovery rate (FDR).

ROKU, an approach based on Shannon entropy statistics, has previously been used to identify genes enriched in a tissue specific manner (*8, 52*). This approach calculates an entropy score of 1, 0 and -1, for depleted, no change, or enriched, respectively, for each gene across cell or tissue specific samples. A gene could be considered as enriched or depleted in no more than half of the cell types. ROKU uses a subset of constitutively expressed genes to determine empirical baseline distributions of entropy scores and to calculate a threshold to call significantly enriched genes. Since the batch effect cannot be modeled for the ROKU method, and since batch correction changes the expression data (i.e. DEGs, based on batch corrected TPM values, have low correlations with batch modeled DEGs [r≤0.5, *p*<0.01, data not shown]), we used upper quartile normalized TPM values to calculate gene entropy. The parameters to determine enriched genes using the Shannon entropy approach were *delta*=0.08, *lowexp*=0.05, *bgfold*=2, *bgmedian*=0.5, and *pvalue*=0.001. The R script and functions are hosted in {https://github.com/plant-plasticity/tomato-root-atlas-2020<u>}.</u>

#### Detection of tomato cell type- and tissue-enriched genes and ontology terms

To identify genes with enriched expression in each cell type we combined the two independent approaches described above. First, DEGs, as determined by limma’s contrasts, were processed with the Brady method (described in (*2*) to identify genes with enriched expression (log_2_FC≥2, FDR≤0.05) in each cell type compared with all other non-overlapping cell types (see **Data S1** for these queries). Next, a union gene set, based on both the Brady and ROKU methods, was obtained for each cell type. A non-redundant list of enriched genes was curated by including only genes with a TPM value ≥ 2 that have the highest expression in the target cell type compared with all other cell types, excluding 35S and Actin (**Data S1**). To differentiate between the general cortex (gCOR), which includes the exodermis, and the inner cortex (iCOR), which includes only the two inner cell files of the cortex, the union set of enriched cortex genes was not filtered against the exodermis, resulting in a partially redundant list with the exodermis of gCOR-enriched genes.

Enrichment analyses of <u>G</u>ene <u>O</u>ntology (GO) terms and MapMan bin terms were carried out with the CTEGs. GO enrichment analysis was done with the GOseq R package (*53*), using the effective transcript length (Kallisto output) for correction of the length bias present in the data. Gene Ontology annotation (ITAG3.2) was downloaded from Sol Genomics Network (<u>solgenomics.net</u>). A term was considered significantly enriched if it has a p-value<0.05 and a fold enrichment > 1. Fold enrichment was calculated as (genes annotated with a term in the query dataset / total genes in the dataset) / (genes annotated with a term in the background set / total expressed genes) (**Data S2**). The hierarchical and non-redundant MapMan bin terms (*54*) were used as a reference database for functional enrichment analysis using the FunRich tool (v3.1.3 www.funrich.org; (*55*). Mapping files (ITAG2.3) were retrieved from the MapMan Store (<u>mapman.gabipd.org</u>). Enrichment analysis was carried out independently for four hierarchy levels; the two top- and lower-level terms of the MapMan hierarchy. Terms with a fold enrichment > 1 were selected for p-value adjustments using p.adjust function in R. Only terms with an FDR < 0.15 were considered significantly enriched (**Data S2**).

#### Identification of tomato cell type-enriched genes in field and pot-grown plants

Four TRAP lines profiled in agar plate-grown plants were also profiled in a field experiment (driving expression in the endodermis (*SCR*), meristematic zone (*RPL11C*), meristematic cortex (*SlCO2*) and whole root (*35S*). The cell type-enriched genes were derived from comparisons involving only these cells and were performed as described for the whole atlas dataset. Gene lists were filtered for FC>1 in the case of the field experiment and a FC greater than 2 for the tomato atlas experiment and can be found in **Data S5**. Genes identified as cell type-enriched in both the field and atlas experiments were considered as “core” cell type genes (**Data S5**). GO and MapMan enrichment analysis was carried out for the cell type-enriched genes derived from four cell type comparisons (separate for atlas and field experiment as well for the list of core genes) in the same manner as for the full dataset (**Data S5**). Enriched categories and annotations shared between the atlas and field experiment (meaning enriched among CTEGs in both the field and atlas, respectively) can be found in **Data S5**.

#### Co-expression network analysis

Co-expression network modules were created with the WGCNA R package version 1.68 (*18*). Individual libraries from each growth condition (agar plates, field, pots) were quantile normalized together and 75% of the most variable genes were used for analysis. A soft threshold of 5 was used to create a scale-free network. An unsigned network was created using the *blockwiseModules*-function with the bicor correlation measure and the following parameters: maxPOutliers = 0.05, mergeCutHeight = 0.35 and maxBlockSize = 25000. Gene Ontology and MapMan enrichment analysis for genes from each individual module was carried out in the same manner as for the cell type-enriched genes. A list of genes assigned to each module, as well as GO and MapMan annotations enriched in each module, can be found in **Data S4**.

#### Mapping of chromatin accessibility, identification of Transposase Hypersensitive Sites, and evaluation of accessibility changes between cell types

A flow chart describing all steps of THS identification and analysis is available (**Fig. S22**). For each sample, 40-bp PE sequencing reads were trimmed using CutAdapt 2.0 and parameters for Nextera libraries (*56*). Trimmed reads were mapped using BWA-mem (*57*) software with default parameters to SL3.0 (https://www.ncbi.nlm.nih.gov/assembly/GCF_000188115.4/).

Aligned sam files were converted to bam format using Samtools 1.6 (*58*), sorted and filtered to retain only reads that had a mapping quality score of 2 or higher, and filtered to retain only reads that mapped to true nuclear chromosomes. The tomato genome is repeat-rich (*59*), and thus to account for mis-annotation of repeats as well as unknown copy number variation, we used methods described for human DNAseI hypersensitive site sequencing to remove high-depth sequencing regions (*60*). Genomic DNA-based ATAC-seq libraries from root tips were sequenced on the NextSeq 500 at the University of Georgia Genomics and Bioinformatics Core to obtain 36-bp paired-end reads (*6*). After mapping with Bowtie2 (*61*) to SL3.0, the number of reads mapping to each position in the genome was determined. Next, the number of reads within 150-bp sliding windows (step size 20-bp) was counted and plotted in a histogram (**Fig. S23A**). The top 0.1% most-accessible windows were then identified, merged and removed from cell type ATAC-seq sample bam files. **Fig. S23B** demonstrates the distribution of sizes for these high sequencing depth regions. Masked bam files were then sub-sampled to a final count of 25 million reads.

In order to determine the best window size for peak calling, we took the most deeply sequenced sample (WOX_O08) and called peaks using three different window size parameters relative to increasing sizes of randomly sampled reads (**Fig. S24**). From these, we determined that a 10 kb window size (the HOMER default) led to an asymptote at ∼25 million reads. Peak calling was thus performed using the “Findpeaks” function of the HOMER 4.9 package (*62*) with the parameters “-size 150”, “-minDist 150” “-region” and “-regionRes 1”. These regions are hereby referred to as Transposase Hypersensitive Sites (THSs).

Independent of peak calling, “per base” bed files were also created. Specifically, the number of aligned reads within a bam file, or cut counts, were tallied at each position within the tomato genome. Any position with zero cut counts was discarded. Results were reported in standard bed file format. For visualization of data within a genome browser, bigWig files were also created from the sub-sampled with Deeptools 3.1.0 (*63*), with the parameters “--binSize 20”, “-- normalizeUsing RPGC”, “ --effectiveGenomeSize 807224664”, and “--extendReads”.

To find replicable THSs across a minimum of three, or a maximum of four biological replicates within a cell type, THSs from the replicates were merged into master replicate THS file using Bedtools 2.27 “merge” (*64*). Pairwise comparison of cut counts between cell types is found in **Fig. S25**. Next, for each replicate, the number of cut counts within each region in the master replicate THS file were counted using BEDOPS 2.4.33 ‘bedmap,’ with the replicate perbase bed file as the map file and the master replicate THS bed file as the reference (*65*). The coefficient of variation was then calculated for each THS across the replicates and the top 15% most variable THSs were removed from further analysis. (**Fig. S26**)

After repTHSs from each cell type were identified, repTHSs were merged into a master union THS bed file (uTHS bed file) using bedtools “merge”. These uTHS regions were then used for downstream analysis of motif enrichment and differential THS identification.

#### Identification of cell type-enriched accessible regions (CTEARs)

The union THSs from the cut counts were analyzed using a similar approach to the detection of cell type- and tissue-enriched genes. CTEARs were identified with limma (v3.34.9) using the *voom()* function with a model including cell type and preparation group to account for batch effects (*∼0+Cell_type + Prep_group*). To identify dTHS (<u>d</u>ifferential <u>T</u>ransposase <u>H</u>ypersensitive <u>S</u>ites) that are enriched in a cell or tissue type, contrasts between the markers whose domain does not overlap were not considered (**Data S1)**. If two tissues overlapped domains, as determined by GFP marker lines, these contrasts were not considered in the enrichment calling after limma. For example, xylem and vascular markers have overlapping domains, thus the Xylem vs Vascular contrast was not used when calling enriched THS, but Xylem vs all other cell types were used. **Data S1** contains contrast exceptions for each cell type. dTHSs were considered cell type-enriched if a logFC > 0 and FDR < 0.2 was present in at least one of the comparisons. ROKU (*8, 52*) was also used to determine THSs with enriched abundance. The script used to analyze these data is hosted in {https://github.com/plant-plasticity/tomato-root-atlas-2020<u>}. The union of both methods was considered for the identification of CTEARs. To resolve dTHS enriched in multiple cell types, we sorted the log_2_-transformed pseudo-cut counts (using *log2(THS +1)*) and interrogated each THS that showed enrichment in more than one cell type. If the logFC between the two THS with the highest abundance was > 0.5, the THS was considered to be enriched in the cell type with the highest cut-counts.</u>

The enrichment of expressed genes near CTEARs in the target cell type was tested using the *fisher.test()* function in R. The contingency matrix for each cell type was built by calculating the fraction of genes near CTEARs with TPM >2 compared to a null background (genes near THS in the rest of the cell types). Multiple-testing correction was performed using False Discovery Rate (*66*).

Please see **Data S6** for a summary of the ATAC-seq data, uTHS regions, and dTHS regions.

#### Annotation of Transposase Hypersensitive Sites

For each cell type’s replicate THSs, the mid-point of the THS region was determined. We then used bedmap with the parameters “--echo”, “--sum”, “--delim “\t”” to determine which genomic feature they overlap ITAG3.2 (https://solgenomics.net). **Fig. S27** shows the distribution of annotated regions per cell type.

In order to assign genes to a THS (uTHS), we used bedtools “distance -D” to calculate the distance between a THS and the nearest nearby gene. uTHSs were filtered for distances within 4-kb upstream of the TTS, overlapping a gene, or 1-kb downstream of the TTS. Promoter sizes were based on previous findings in ChIP-seq data for the transcription factor SlMYC2 (*67*).

#### Motif enrichment and TF networks

##### Motif database construction

Motif files were downloaded from CisBP for Weirauch, DAP-seq, Franco-Zorilla, and Sullivan motif datasets (*29, 68–70*). If a motif from the protein binding array studies overlapped with the DAP-seq database, it was discarded.

##### 1-kb promoter network construction

1-kb upstream sequences of the TSS for each group of cell type-enriched genes were identified. Next, these sequences were used to perform motif enrichment with our custom motif database using Meme Suite AME (*71*), with the parameters “--scoring avg”, “--method fisher”, “--hit-lo-fraction 0.25”, --evalue-report-threshold 2000”, “--control”, “--shuffle--”, and “--kmer 2”. Next, the motif enrichment files for all cell types were converted to a matrix file where each row represents a transcription factor motif and each column represents the adjusted p-value for that motif in a given cell type. Motifs were then filtered for ones that were significantly enriched in at least one cell type (padj>=0.05). Next, the motifs were filtered for genes in Arabidopsis that have positively correlated root TRAP expressologs in tomato. Motifs that had a corresponding tomato expressolog were then filtered for expression in the given cell type (TPM >= 2). The matrix file was then split by motif family and adjusted p-values were visualized in R 3.6 (https://www.R-project.org/) <u>using Pheatmap</u> (https://cran.r-project.org/web/packages/pheatmap/index.html).

##### uTHS promoter network construction

For each cell type-specific group of genes, uTHSs were identified that were 4-kb upstream from TTS, overlapping genic regions, or 1-kb downstream from the TTS. This was done using the bedtools “closest” tool, with the parameter “-D” and the uTHS files and the bed files for the genic locations for the cell type-specific genes. Fasta sequences for these regions were obtained using bedtools “getfasta”. Next, motif enrichment was performed using Meme suite AME using the same parameters as the 1-kb upstream regions. Motif filtering and heat map creation were performed as they were for the 1-kb upstream regions.

##### Cell type-unique network construction

To identify unique cell type functions and their underlying regulation, we also constructed unique cell type networks (**Fig. S28)**. Transcription factor motifs that were significant and unique to each cell type were identified separately for uTHSs and 1-kb promoters. Next, we filtered the unique transcription factor motifs for positively correlated expressologs in tomato and whether they were expressed in the cell type of interest (TPM >= 2). After identification of unique expressologs, the union of unique transcription factors was taken between the 1-kb promoters and uTHS networks. These union networks comprising transcription factor motifs, as well as their targets, were then visualized with Cytoscape 3.7.1. (*72*). Please see **Data S7** for unique cell type network files.

#### Nitrogen network overlap

To test for enrichment of the exodermis-inferred network and an Arabidopsis nitrogen metabolic network (*27*), we filtered the expressolog list for positively correlated expressologs (cor > 0). The Arabidopsis nitrogen network contains a total of 429 genes. Of these, 362 have at least one positively correlated expressolog in *S. lycopersicum*. A total of 301 genes have an expressolog for both the TF and its target promoter in a TF/promoter interaction. We calculated if the overlap between the *S. lycopersicum* exodermis-inferred network genes and the expressologs of the orthologous nitrogen network genes in tomato was greater than expected by chance using the *fisher.test()* function in R with *alternative=“greater” (***Data S9)**.

#### A comparative analysis of root cell type-atlases across species

##### Arabidopsis microarray data

CEL files containing data resulting from translatome profiles of Arabidopsis root tips expressing endodermis (*SCR*), vasculature (*SHR*) and whole root (*35S*) markers, as well as from translatome profiles of the meristematic zone (*RPL11C*) and meristematic cortex (*AtCO2*) marker lines, were downloaded from GEO (<u>GSE14493)</u> (*5*). The raw files were reanalyzed with the limma package (*50*) and normalized log_2_ intensity values can be found in **Data S8.**

##### Rice RNA-seq data processing and analysis

Rice data was processed as described above for tomato RNA-seq data processing and analysis with the following modifications: trimmed reads were pseudo-aligned to IRGSP-1.0 transcriptome (cDNA, https://rapdb.dna.affrc.go.jp/index.html) using Kallisto (v0.43.1) (*49*) to obtain count estimates and transcript per million (TPM) values. Splice variants were summed to assess transcript values.

##### Sample integration and clustering of expression profiles

Comparisons of transcript abundance were conducted for four homologous cell types (meristematic cortex, endodermis, vasculature and meristematic zone, **Fig. S16**) within and between species. Since genes that undergo duplication events rapidly diverge in their expression profiles (*31, 73*) three different orthology maps were generated, two maps based on sequence similarity and one based on sequence similarity coupled with expression correlation (i.e., “expressologs”) (**Data S8**). The first orthology map includes 2,642 1:1:1 orthologs based on sequence homology, using Phytozome v12 gene families. Phytozome predicted gene families were generated using genome sequence data from 57 plant species. In Phytozome, the relationships between genes and species are determined by InParanoid, which uses an all-vs-all BLAST alignment of pair-wise proteomes to identify orthology groups (*74*). Phytozome uses an *S. lycopersicum* ITAG2.4 annotation, while data for all other analyses in Figures 1 through 3 are from the ITAG3.2 genome. Hence, genes annotated in ITAG3.2 that are absent from ITAG2.4 were assigned to a gene family based on a blastp search against *A. thaliana* cDNAs (- max_target_seqs 1), with an E value cutoff of < 0.01.). To identify 1:1:1 orthologs, only predicted gene families with one gene from each species were included (**Data S8** and **Fig. 4B**). The second orthology map includes 3,505 1:1 orthologs, based on sequence homology to Arabidopsis. This map takes advantage of the plant-specific MapMan tool, which was originally developed for Arabidopsis, and currently supports more than 80 plant species (https://mapman.gabipd.org/home) (*75*). The freely available MapMan annotation files of tomato and rice were parsed to include only 1:1 orthologs that are present in both files (**Data S8** and **Fig. S17B**). Finally, the third orthology map consists of 1,771 Arabidopsis and rice expressologs of tomato with an expression correlation coefficient > 0.6 (**Data S8, Fig. S17**). “Expressologs” are determined using an approach to resolve orthologs by predicting putative functional orthology (i.e., expressologs). This map was constructed using sequence homology, based on OrthoMCL (*76*), and complemented by published expression profile similarity to refine ortholog predictions as described in (*77*).

As previously described, analyses of gene expression variation between species must take into account confounding factors (*30*). Thus, we next considered how to address differences in experimental design between tomato, rice and Arabidopsis, and the fact that *i*) translatome samples were obtained from two expression platforms (i.e. RNA-seq for rice and tomato and microarray for Arabidopsis) and thus possess distinct dynamic ranges (**Fig. S29A**), and *ii*) data obtained from each species was collected and processed in a different laboratory, which drives the clustering of samples (**Fig. S29B**). We accounted for these issues by applying the functions normalizeBetweenArrays() and removeBatchEffect(), from the limma package, which were used to quantile normalize expression values across samples of homologous cell types and 35S (**Fig. S29C**) and to correct for the laboratory effect (**Fig. 4B**), respectively (*50*). Since species and laboratory are completely confounded, by correcting for the batch effect we also removed the contribution of the species to gene expression variation, hence we can only assess the contribution of the tissues. Clustering of expression profiles was carried out for the four homologous cell types as described above for tomato RNA-seq quality control.

##### Root Cell Type TRAP-expressologs

Cell type- or tissue-resolution TRAP data can be utilized to define “expressologs” based on expression variation similarity across homologous root cell types. Ortholog annotations for tomato, Arabidopsis and rice were determined as described in (*77*) with the following modifications: (i) putative gene families that include at least two of the three species were retrieved from the ITAG3.2-updated Phytozome v12 gene family file (described above for the first orthology map); (ii) within each gene family, the Pearson correlation coefficient was calculated for each ortholog pair using the TRAP expression values of homologous cell types and tissues. Tomato and *Arabidopsis* included eight homologous cell types and tissues (EP, COR, MCO, EN, V, PH, MZ and 35S), tomato and rice included six homologous cell types and tissues (MCO, EN, V, MZ, QC and 35S) and Arabidopsis and rice included five homologous cell types and tissues (MCO, EN, V, MZ and 35S). (iii) The correlation matrices were reciprocally parsed to include only the best matching expressolog pairs using each species as a reference (e.g. maximum correlation between Arabidopsis to tomato and tomato to Arabidopsis, based on Arabidopsis as a reference species). (iv) To identify high confidence expressologs and define ortholog annotations for the cell type-enriched genes, only expressolog pairs with a positive correlation and a reciprocal match between Arabidopsis and tomato and Arabidopsis and rice were considered (**Data S8**). These filtering criteria resulted in the identification of 6,059 expressologs between Arabidopsis and rice, and 7,295 expressologs between Arabidopsis and tomato. To detect conserved expressologs, we selected only positively correlated expressologs that maintain the same relationship among the three species, independently of the reference species. To this end, high confidence expressologs among the three species were identified, using each species as a reference. This analysis resulted in identification of 6,293, 6,470 and 6,516 expressologs based on tomato, Arabidopsis and rice as a reference species, respectively. Next, the three datasets were intersected and expressologs with negative expression correlations were excluded, resulting in the identification of 1,555 expressologs that have both identical expressolog relationships independent of the reference species and positive expression correlations (referred to as consensus expressologs) (**Data S8**). Clustering of expression profiles of homologous cell types, based on the consensus expressologs, was done following quantile normalization and batch effect correction of log_2_ expression values, as described for the sample integration and clustering of expression profiles across-species (**Fig. S18**).

##### ANOVA to identify conserved cell type and tissue-specific expressologs

The clustering of consensus expressologs based on tissue identity suggests that some of these genes have conserved tissue-specific patterns of expression (**Fig. S18**). To further explore these expression patterns and to identify consensus expressologs with conserved cell type and tissue-enriched expression we used an ANOVA. Expression values of MCO, EN, V, MZ and 35S were processed for each species separately. For tomato and rice, upper quantile-normalized TPM values were filtered to remove genes with low expression (TPM≤2), followed by adding a prior count of 3 to reduce the contribution of low-abundance genes, followed by log_2_ transformation of tomato data to further correct for differences in sequencing date as described for the RNA-seq quality control and differential expression. For Arabidopsis we used normalized log_2_ intensity values. For each cell type and tissue in each species, we calculated the mean gene expression, if up to three biological replicates existed, or median gene expression, if four biological replicates existed. Next, the three datasets were combined based on the 1,555 consensus expressologs. The 15 sample mean/median values were quantile normalized and corrected for the batch effect arising from the different laboratories, using the functions normalizeBetweenArrays() and removeBatchEffect() from the limma package, respectively, as described for the sample integration and clustering of expression profiles. To detect genes with conserved cell type and tissue specific expression the R Stats functions lm(), aov() and the function HSD.test(), from the agricolae package (*78*), were used to fit a linear model, to test the effect of the tissue on gene expression and to identify the cell types or tissues with a significant effect, respectively. The consensus expressologs with the top 15% F-values were filtered to include genes with a conserved enriched or depleted expression in one cell type, based on a Tukey test (p-value≤0.1) (e.g., conserved high expression in the MZ compared with the other three cell types). Finally, these genes were filtered against constitutively expressed genes (CEGs) within each species, as described below, resulting in the detection of 139 conserved cell type-specific expressologs (**Data S8**). Thirty-seven of these genes showed conserved cell type/tissue enriched expression among the three species (**Fig. 4C**).

##### Detection of cell type and tissue-enriched genes and ontology terms across species

To allow a balanced comparison of CTEGs across species we used the same pipeline as described for the detection of cell type- or tissue-enriched genes within tomato (queries and parameters are specified in **Data S8**). Orthologs were resolved using the high confidence expressologs between Arabidopsis and the two other species, as described above for the Root Cell Type TRAP-expressologs (**Data S8**). Enrichment of GO and MapMan ontology terms of cell type-enriched genes were determined for each species as described above (**Data S8**). GO annotations were downloaded for the TAIR10 genome assembly (<u>arabidopsis.org</u>) and retrieved from Ensembl with the biomaRt package (*79*), for Arabidopsis and rice, respectively. Overlapping ontology terms among homologous cell types were visualized using Circos (*80*) based plots, which also included 3-way overlaps, and their enrichment was evaluated by Fisher’s exact test using fisher.test() function in R (**Fig. 4D**, **Data S8**).

##### Constitutively expressed genes (CEGs)

To identify CEGs we used a fold change difference < 1.5 between the maximum and minimum TPM or intensity values of each gene across the five homologous cell types/tissues (i.e., MCO, EN, V, MZ and 35S), together with a cutoff of a per gene median expression > median expression of each species. These filtering criteria resulted in detection of 308, 1,154 and 1,523 CEGs in tomato, Arabidopsis and rice, respectively (**Data S8**). Orthologs were resolved using the high confidence expressologs between Arabidopsis and the two other species, as described above for the Root Cell Type TRAP-expressologs (**Data S8**). Enrichment of gene and MapMan ontology terms of CEGs were determined for each species as described above for the detection of cell type and tissue-enriched genes and ontologies across species (**Data S8**). Assessment of the enrichment of the overlaps between the ontology terms of the CEGs compared with the CTEGs was carried out with a Fisher’s exact test using fisher.test() function in R.

#### Phylogenetic tree construction

Phylogenetic trees are generated by a custom script which identifies homologous amino acid sequences of genes related to the target gene using BLASTp v2.9.0+ (NCBI BLAST) (arguments: -max_target_seqs 15 -evalue 10E-6 -qcov_hsp_perc 0.5 -outfmt 6) against proteomes of 15 species (*Arabidopsis thaliana, Brachyodium distachyon, Chlamydomonas reinhardtii, Glycine max, Malus domestica, Medicago truncatula, Nicotiana tabacum, Quercus suber, Oryza sativa, Solanum lycopersicum, Selaginella meoellendorffii, Solanum tuberosom, Sorghum bicolor, Vitis vinifera, Zea mays*). To remove identical or near-identical sequences, clusters are generated using CD-HIT v.4.8.1 (*81*)(argument: -c 0.99). MAFFT v7.271 was used for multiple sequence alignment (arguments --auto --reorder --quiet)(*82*). Alignments were trimmed using trimAl v1.2rev59 (*83*) (arguments: -gt .5). Phylogenetic trees were built using RAxML v8.2.12 (*84*); (arguments: -f a -m PROTGAMMAAUTO -N 100; The seeds for the arguments -x and -p were generated calling a $RANDOM variable for each in a Unix environment). Trees were further refined by selecting proteins from a single relevant subtree and repeating the alignment and tree construction process from MAFFT.(*81*)(*83*)

#### Gene Orthology Determination

To identify the best orthologs between Arabidopsis and tomato we used a gene family tree-based approach (as described for xylem-enriched TFs, the VND TF family and the “core” endodermis-enriched ZAT gene. We constructed gene family phylogenies using a maximum likelihood (ML) algorithm as described above. The topology of the resulting tree was analyzed within the clade and all successive clades dependent on the point where bootstrap values were no longer well supported. In the cases where orthology was defined, it was done so based on the position of the target tomato gene relative to its closet Arabidopsis ortholog.

We identified the closet possible orthologs as follows: (*At4g34590; AtbZIP11*) to be a likely ortholog for SlBZIP11(*Solyc03g033730) (***Fig. S4F)***, (At4g08150; AtKNAT1*) as a likely ortholog for *SlKNAT1*(*Solyc04g077210) (***Fig. S4G)**, and *AT1G30490 or AT2G34710* (paralogs on the same clade) as possible orthologs for *Solyc02g069830; SlPHB/PHV* and *Solyc02g024070* (un-named) **(Fig. S4H)**, and *AT1G02030-ZAT 4 and AT2G4512-ZAT9* (paralogs on the same clade) as orthologs to *Solyc01g090840* (**Fig.S4I)** In the latter two cases (*SlPHB/PHV* and *SlbZIP11*) it was impossible to define a single Arabidopsis ortholog for the tomato gene. For *SlPHB/PHV*, we additionally carried out pair-wise nucleotide BLAST comparisons for candidate genes – *AtPHB (At2g34710)* to *Solyc02g069830* = 76.81% similarity; E value of 0.0*; AtPHB (At2g34710*) to *Solyc02g024070* = 79.26% similarity; E value of 0.0; *AtPHV (At1g30490)* to *Solyc02g069830* = 74.11% similarity; E value of 0.0; *AtPHV (At1g30490)* to *Solyc02g024070* = 77.29% similarity; E value of 0. Again, it was impossible to discriminate between these two possibilities, therefore we refer to *Solyc02g069830* as *SlPHB/PHV*.

#### Ranking candidate xylem regulatory TFs – Intersection of QTL and eQTL data

Genetic intervals significantly associated with variation in xylem cell number were identified using data reported in (*12*). Introgression lines containing these significant genetic intervals were then screened for significant *cis-*eQTL (*13*) of (1) TF loci enriched in tomato xylem cells or vascular tissue, or of (2) HDZIPIII family putative orthologs.

#### Transcriptional reporter construction and imaging

Promoters of the exodermis-enriched *WRKY* (*Solyc02g071130*) and *MYB* (*Solyc02g079280*) TFs were cloned from *Solanum lycopersicum* cultivar M82 genomic DNA. Cloning primers (**Data S9**) were designed to amplify 2,130 bp and 3,408 bp upstream of the translational start site of WRKY and MYB, respectively, using the tomato reference genome annotation ITAG3.2 (https://solgenomics.net). The promoters were amplified from genomic DNA using Phusion DNA polymerase (New England Biolabs). Amplified fragments were cloned into pENTR5’Topo (Invitrogen) and sequences were confirmed by Sanger sequencing. LR Clonase II Enzyme mix (Invitrogen) was used to clone the promoters upstream of a *nlsGFP-GUS* reporter gene fusion in the binary vector pMR074 (*4*), which also contains a ubiquitously expressing plasma membrane marker TagRFP-LTI6b. The binary vectors were used for hairy root transformation as described below. Transgenic hairy root fluorescence was visualized using Confocal Laser Scanning Microscopy with a Zeiss Observer Z1 LSM700 (Zeiss) microscope (water immersion, ×20 objective) with excitation at 488 nm and emission at 493–550 nm for GFP and excitation at 555 nm and emission at 560–800 nm for mRFP. Images were taken at approximately 1 cm from the root tip.

#### Overexpression construct design and cloning

The coding sequence (CDS) for target genes (**Data S9**) was obtained from the Sol Genomics database (https://solgenomics.net - ITAG version 3.2). CDS were amplified from tomato (*Solanum lycopersicum* cv. M82) cDNA. In brief, total RNA was isolated from 50mg of tomato root tissue using the Zymo-Direct-Zol RNA Miniprep Plus Kit (Zymo Research-catalog#R2071) according to manufacturer’s instructions and treated with RNase-Free DNase (1unit/10µl). 1µg of DNAse-treated RNA was reverse-transcribed into cDNA using oligo(dT) primers and SuperScript III Reverse Transcriptase (SuperScript III First-Strand Synthesis System; Invitrogen) per kit instructions. Cloning primers (**Data S9**) were designed to PCR amplify the CDS without the stop codon. PCR products were purified from the <u>agarose</u> gel (QIAquick Gel Extraction kit. Catalog#28704) for subsequent recombination and cloning.

Purified cDNAs were introduced into the pENTR/D-Topo vector (Invitrogen). The resulting pENTR plasmids were then LR recombined (LR Clonase II Enzyme mix; Invitrogen) into the pGWB417 binary destination vector (Addgene plasmid # 74811; http://n2t.net/addgene:74811; RRID:Addgene_74811) containing a 35S promoter driving the expression of the CDS. All constructs were confirmed by Sanger sequencing.

#### Site directed mutagenesis for miRNA resistant HD-ZipIII TF constructs

A point mutation causing a silent substitution in predicted miRNA binding site of *Solyc03g120910, Solyc02g069830* was created with the QuikChange II XL following the provided protocol (Agilent-catalog no. 200521, **Data S9**). This mutated cDNA was then cloned into PGWB417 as described earlier. Mutagenesis was confirmed by Sanger sequencing.

#### *Rhizobium (Agrobacterium) rhizogenes* transformation

*Rhizobium rhizogenes* (Strain ATCC 15834) transformation followed the protocol previously described (*4*). Briefly, competent *R. rhizogenes* was transformed by electroporation with the desired binary vector, plated on nutrient agar (BD 247940) plates with the appropriate antibiotics (spectinomycin, 100 mg L−1), and incubated for 2-3 days at 28-30°C. *R. rhizogenes* colonies passing selection were inoculated from plates into 10 mL nutrient broth liquid medium (BD 90002-660) with the appropriate antibiotics (spectinomycin, 100 mg L−1) and were grown overnight at 30°C with shaking at 200 rpm. This culture was used to transform 40 to 50 fully expanded tomato cotyledons grown in sterile conditions for 8-10 days (just before the first true leaves emerge). Using a scalpel, 8-10 day old M82 cotyledons were cut and immediately immersed in the bacterial suspension at an optical density of 600 nm in Murashige and Skoog (MS, 1X) liquid medium for 20 minutes and then blotted on sterile Whatman filter paper and transferred (adaxial side down) onto <u>MS</u> agar plates (1X with vitamins, 3% sucrose, 1% agar) without antibiotic selection and incubated for 3 days at 25°C in dark. The cotyledons were then transferred to <u>MS</u> plates with Vitamins (MSP09-10LT), 1% agar and 3% sucrose with a broad spectrum antibiotic cefotaxime (200 mg L−1) and kanamycin (100 mg L−1) for selection of successfully transformed roots and returned to 25°C. At least three to five independent roots develop from each cotyledon. Antibiotic-resistant roots that emerged were further transferred to new selection. Fifteen independent roots were subcloned for each construct for further analysis.

#### Quantitative reverse transcription–PCR of overexpression lines

All quantitative <u>RT</u>-PCR primers were designed with Primer3Plus software (http://www.primer3plus.com/) (**Data S9**). Primers were designed to amplify a 100-150 bp region near the 3’ end of each target TF coding sequence. qRT-PCR was performed by setting up a 20 μL PCR reaction containing 5ul of cDNA (100ng/reaction) and 200 nM of each primer (PCRBIO Taq DNA Polymerase/Mix- catalog no. PB10.11-05 and EvaGreen dye; PCRBIO catalog number 89138-982). qRT–PCR was performed in a Bio-RAD CFX384-Real Time System with the following thermal cycling conditions: 5 min at 95°C, followed by 40 cycles of 20s at 95°C, 20s at 60°C, and 20s at 72°C. To ensure that PCR products were unique, a melting-curve analysis was performed after the amplification step. The experiment was carried out on a minimum of three independent lines and three technical replicates for each. To determine the fold change of the overexpression line relative to the wild type control (tomato transformed with *R. rhizogenes* with no plasmid), an absolute quantification method was conducted by generating a standard curve for each primer set. Values were normalized to the Ct value of an endogenous control gene (*Solyc07g025390*). The qPCR data for each gene is shown as a relative expression with respect to a control hairy root sample to which an expression value of 1 was assigned. Standard error of the mean (SEM) was then calculated from the normalized expression for each sample represented in the graphs (**Fig. S30**).

#### Histochemistry and confocal laser scanning microscopy for xylem phenotypes

Hairy root tissue was cleared for 4-5 days in ClearSee buffer (*85*). Detection of xylem vessel elements was conducted by incubation of cleared roots in Basic Fuchsin (0.04% w/v in ClearSee) for 24 hours followed by a 1-2 hour wash in the ClearSee buffer before imaging. Confocal Laser Scanning microscopy was carried out on a Zeiss LSM700 confocal with the 20X objective, Basic Fuchsin: 550-561 nm excitation and 570-650 nm detection. Root samples were mounted in ClearSee and scanned. Protoxylem vessel differentiation was first observed at 0.2-0.4 mm distance from the tip, while metaxylem vessels differentiate up to 1 cm from the root tip, in the maturation zone.

#### Statistical analyses for overexpression lines

Comparisons and significance of aberrant xylem phenotype frequencies (*SlbZIP11* and *SlKNAT1* - extra protoxylem or xylem breaks; *SlCORONA* - loss of bilateral symmetry; *SlPHB/PHV* protoxylem at metaxylem position) relative to the wild type control (tomato transformed with *R. rhizogenes* with no plasmid) were determined with a logistical regression method using Generalized Linear Model (GLM) in R Studio software (Version 1.2.5001). The output from the GLM model was then used to determine an odds ratio for each independent line. Analysis was done on 3 independent lines for each overexpression construct with a minimum of 12 biological replicates per line. The results of all statistical tests performed are reported in **Data S3**.

---

**Fig. S1.**
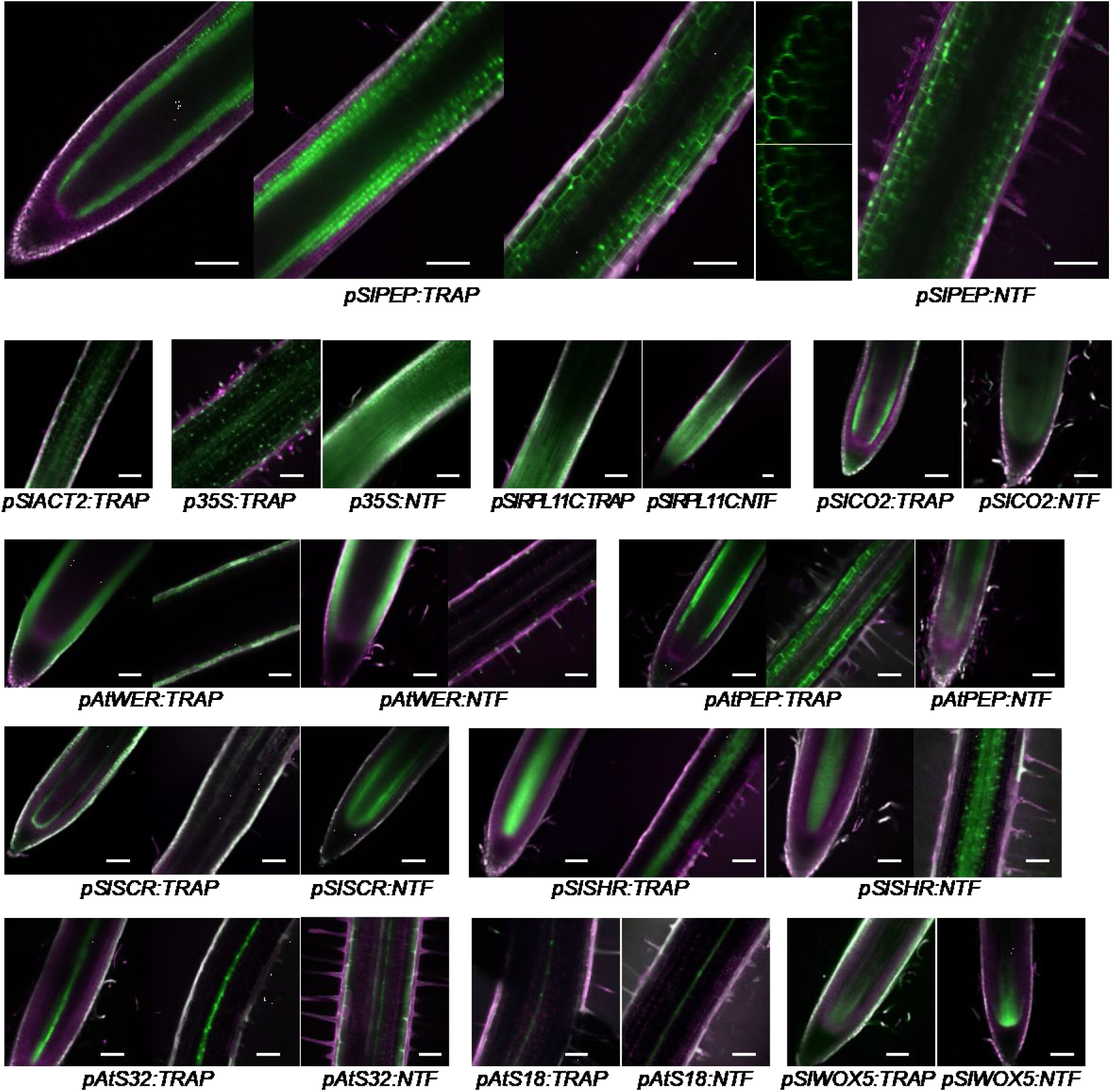
GFP expression in all the marker lines used for this study. Green represents the GFP signal, magenta the autofluorescence, and the overlay of the two appears white. Scale bars represent 100µm.

**Fig. S2.**
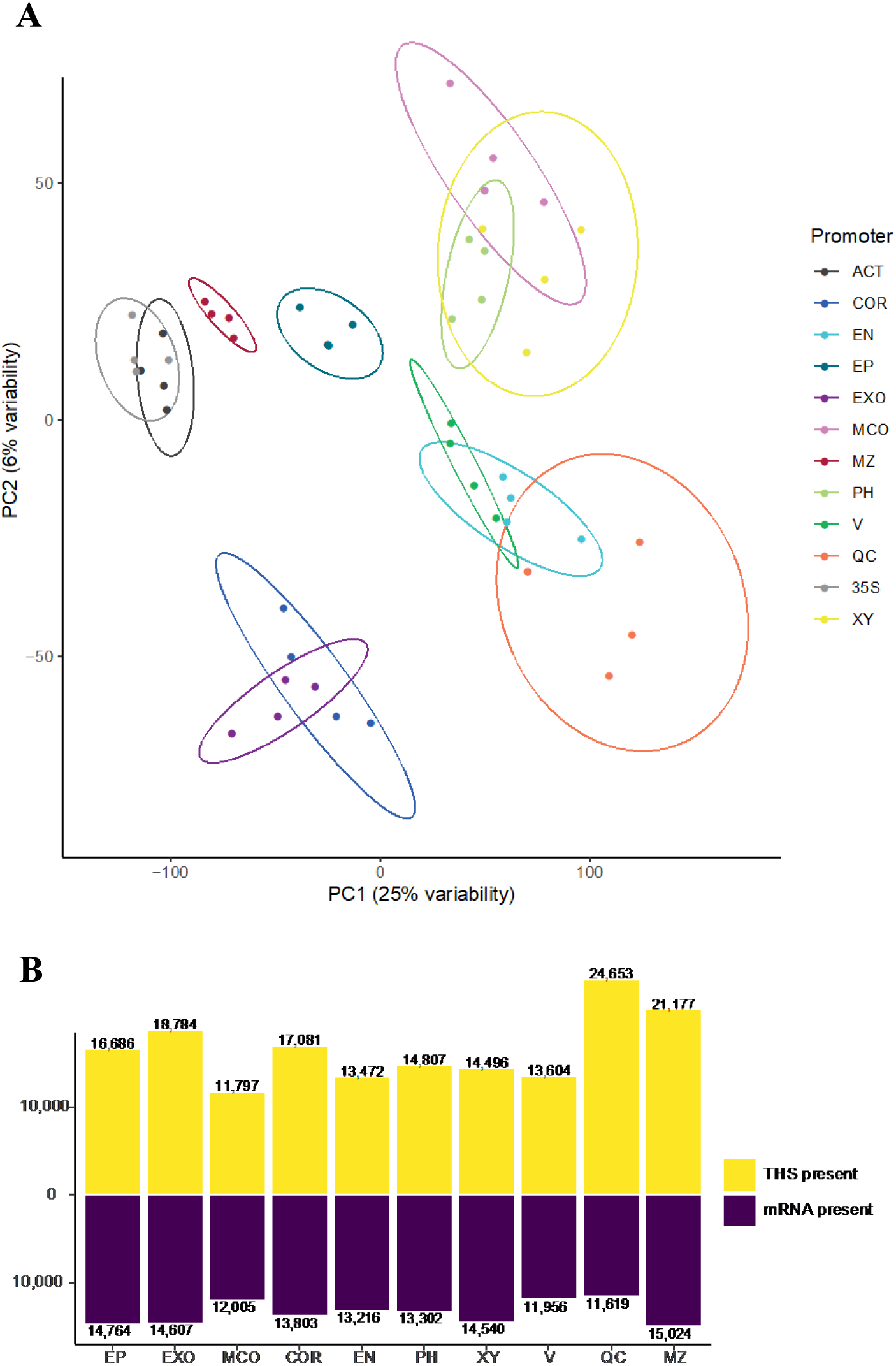
(**A**) **Principal component (PC) analysis of tomato marker-line derived translatomes.** Ribosome-associated transcript abundance after normalization to library size and batch effect correction. Each sample is indicated by a dot and colored by the marker-line. (**B**) Total number of transcripts and THSs detected in each cell type.

**Fig. S3.**
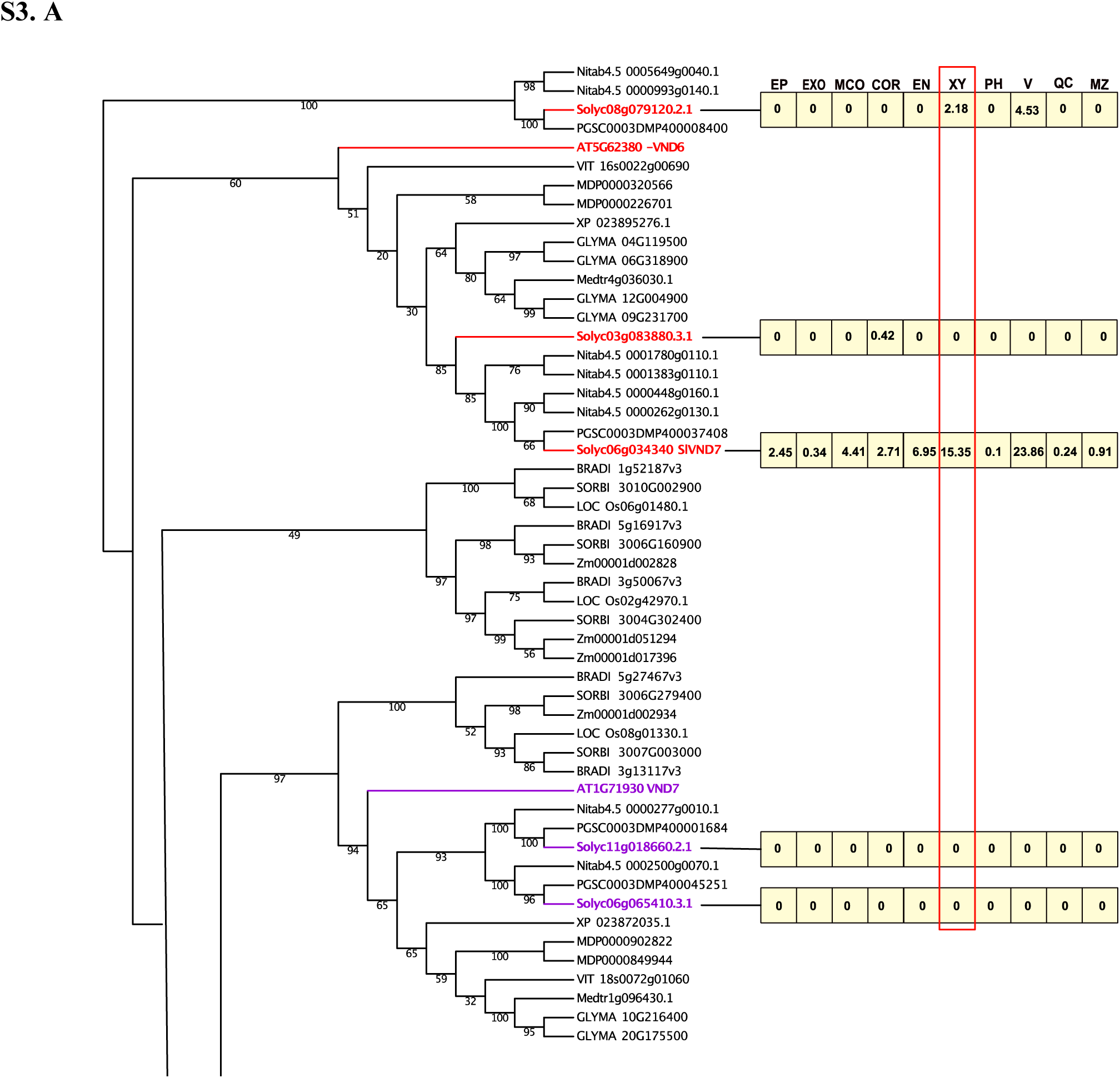
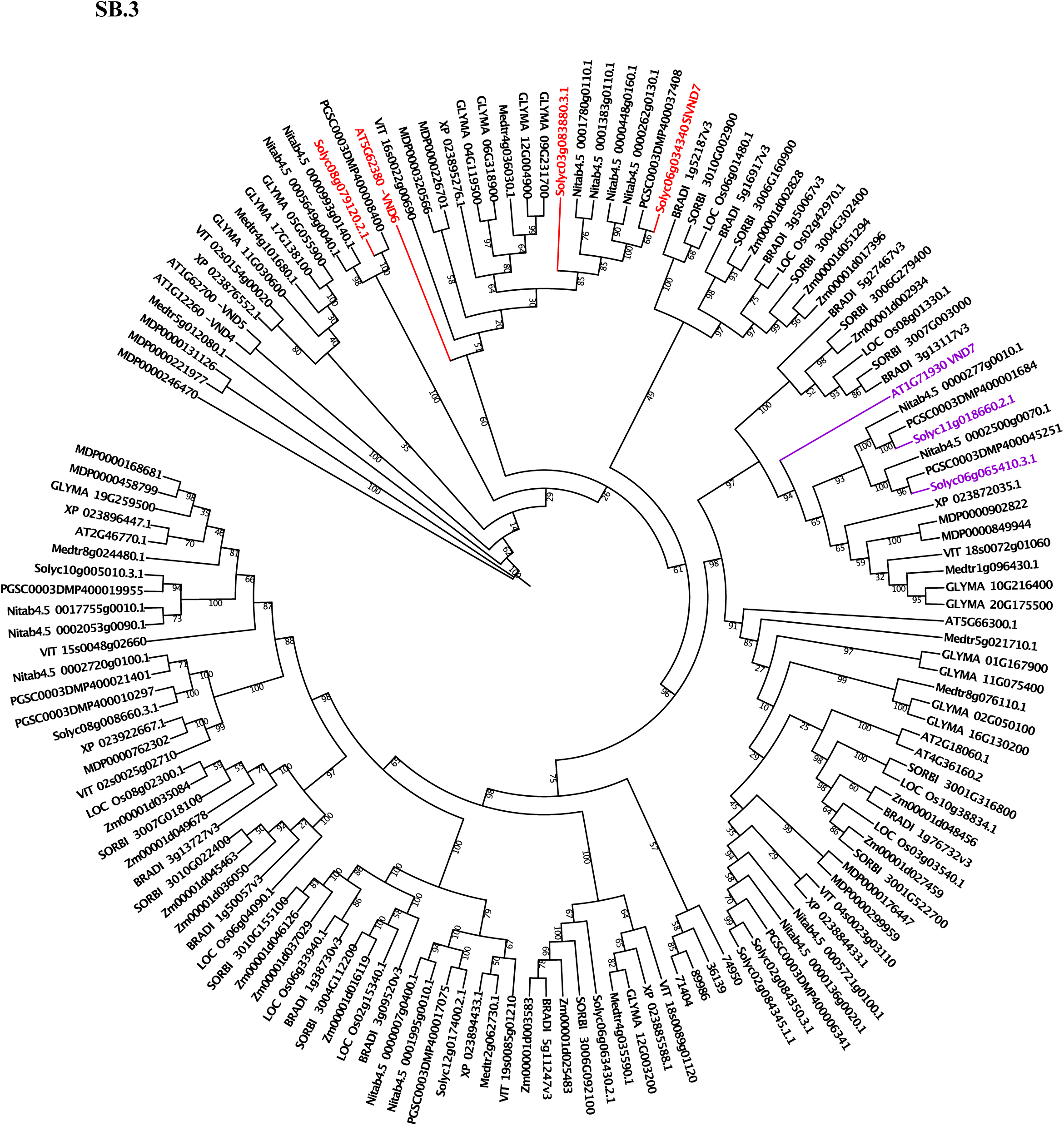
Phylogenetic analysis of full-length VND1-7 proteins in fifteen different species. (**A**) Phylogenetic tree showing VND6 and VND7 clades only. Putative orthologs of *AtVND6* and *AtVND7* in *Solanum lycopersicum* are highlighted in red and purple. Numbers in boxes represent median normalized TPM from our TRAP-RNA-seq data set in each cell type. (**B**) Full phylogenetic tree of *VND1-7*. AT, *Arabidopsis thaliana*; BRADI, *Brachypodium distachyon;* CHLRE, *Chlamydomonas reinhardtii; GLYMA, Glycine max;* MDP, *Malus domestica*; Medtr, *Medicago truncatula*; Nitab, *Nicotiana_tabacum*; XP, *Quercus suber*; LOC_Os, *Oryza sativa*; Solyc, *Solanum lycopersicum*;; Smo, *Selaginella moellendorffii*; PGSC, *Solanum tuberosum*; SORBI, *Sorghum bicolor*; VIT, *Vitis vinifera*; ZM, *Zea mays*. Phylogenetic trees were built using RAxML v8.2.12 (**Materials and Methods**).

**Fig. S4.**
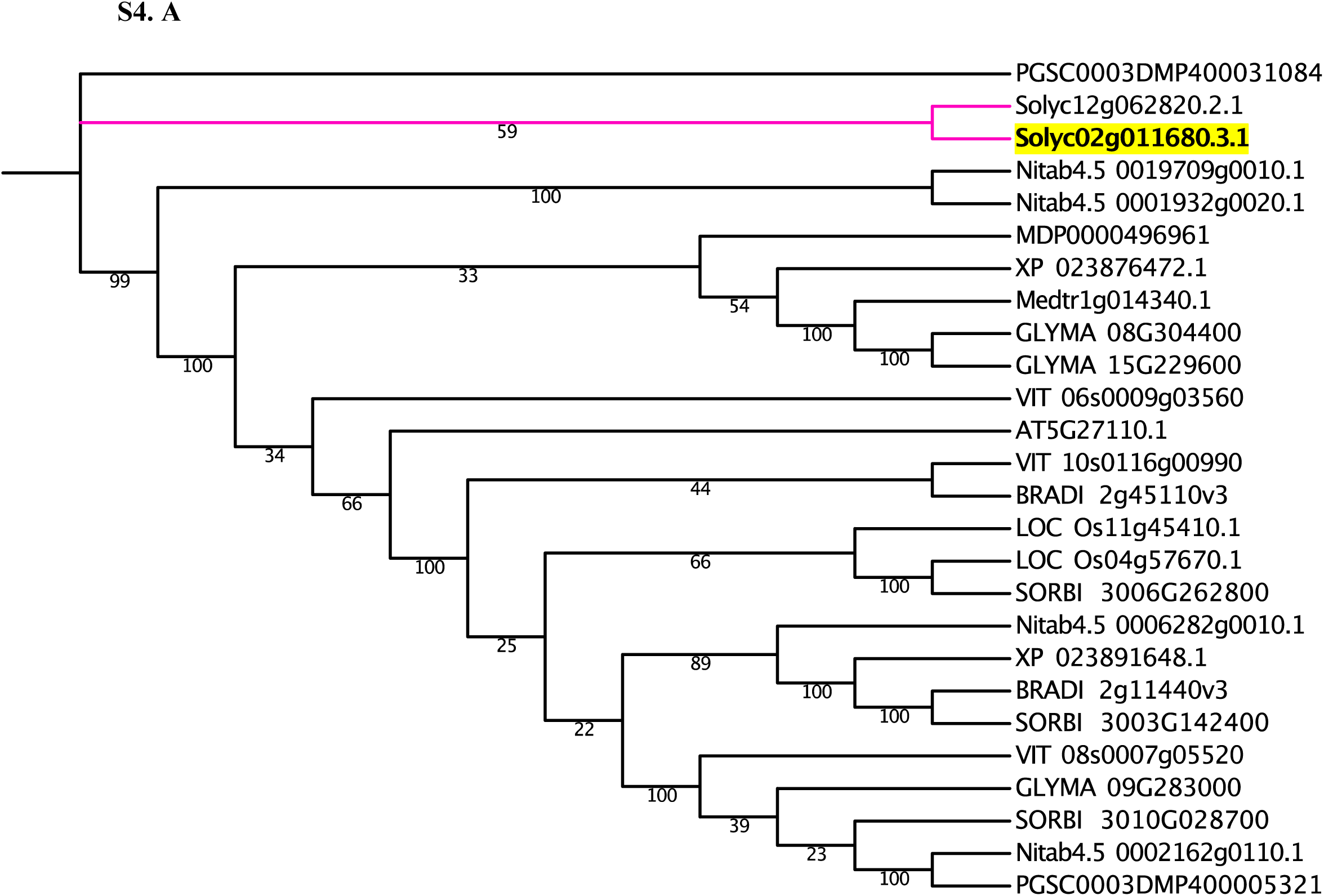
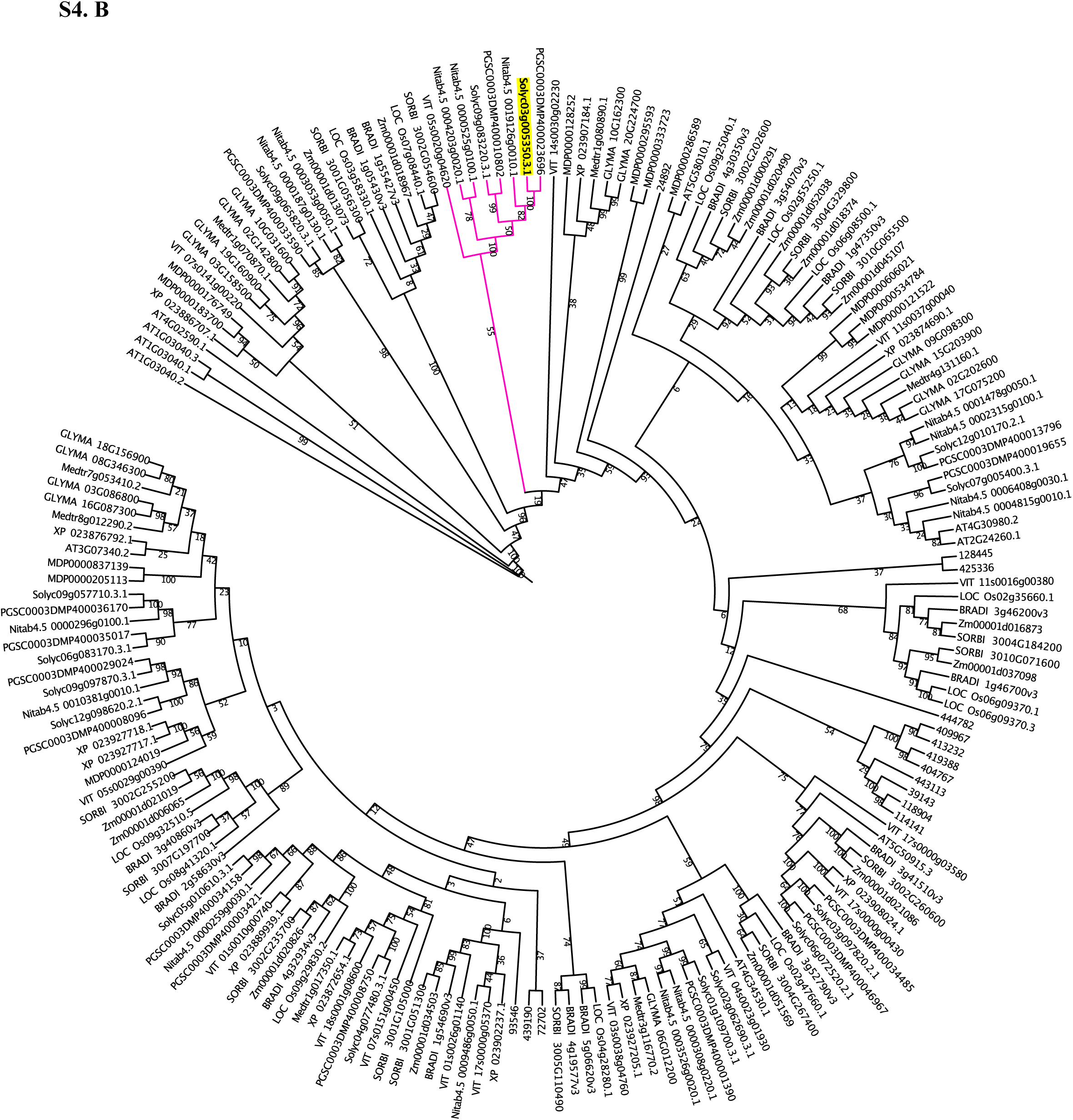
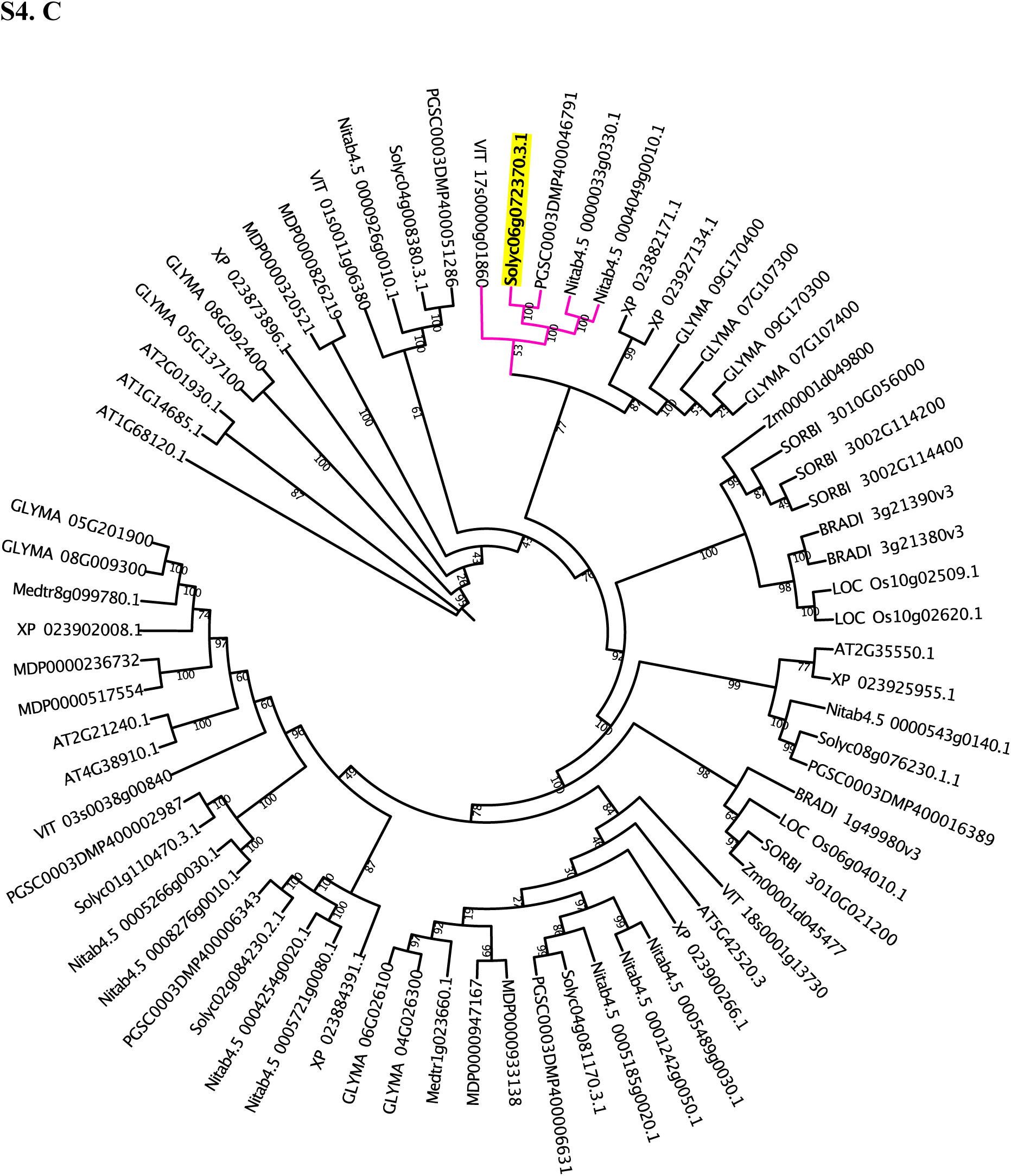
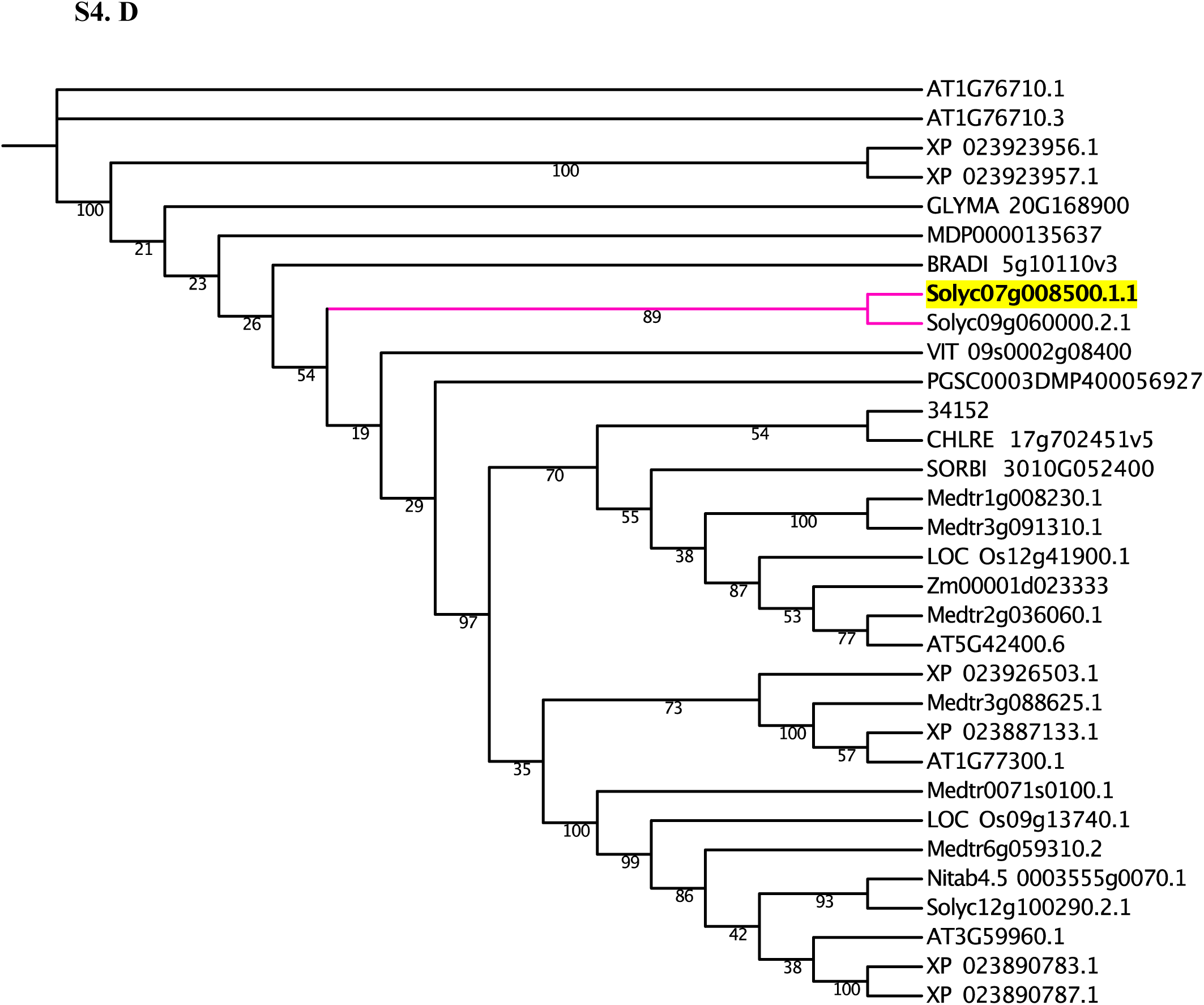
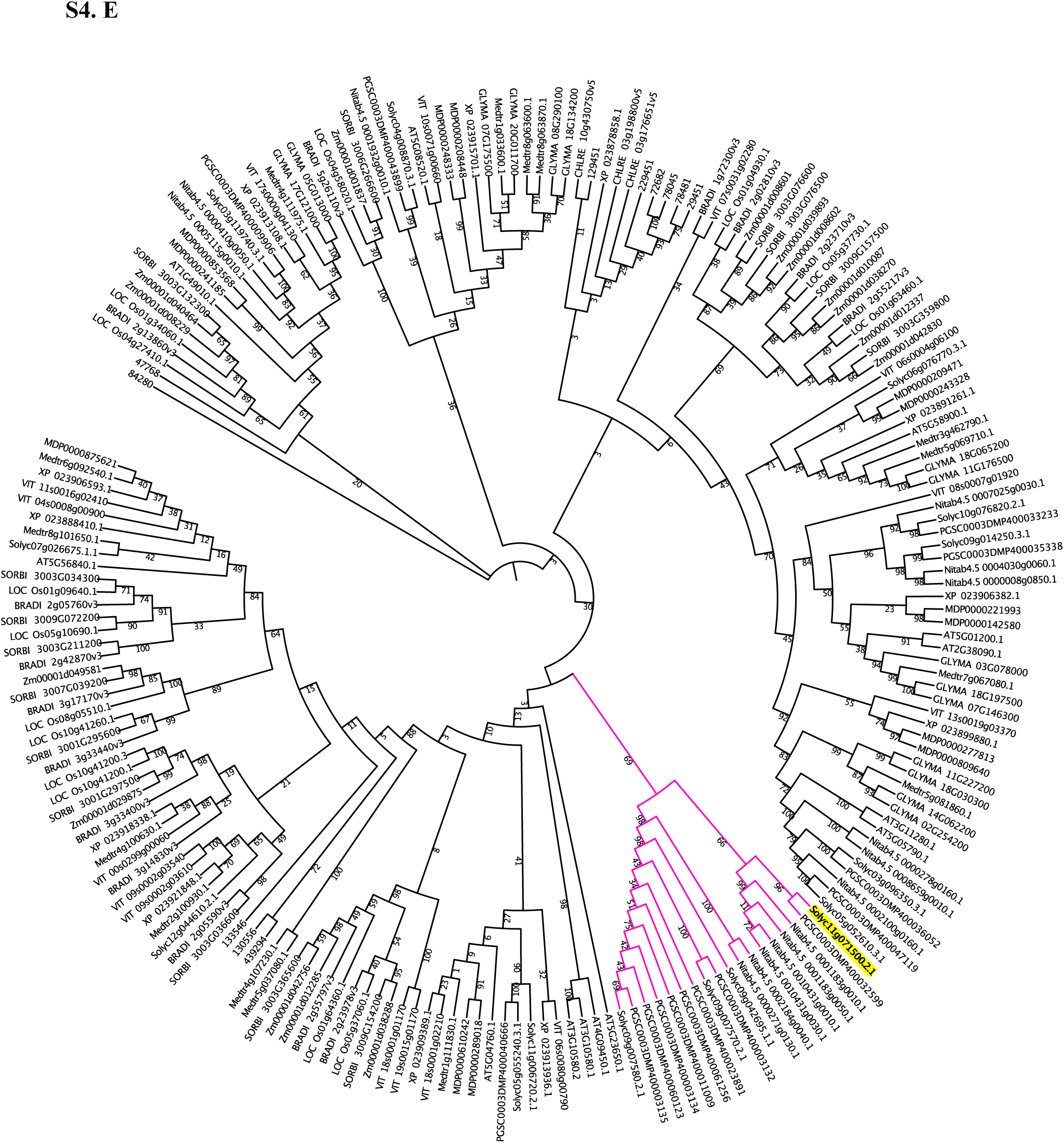
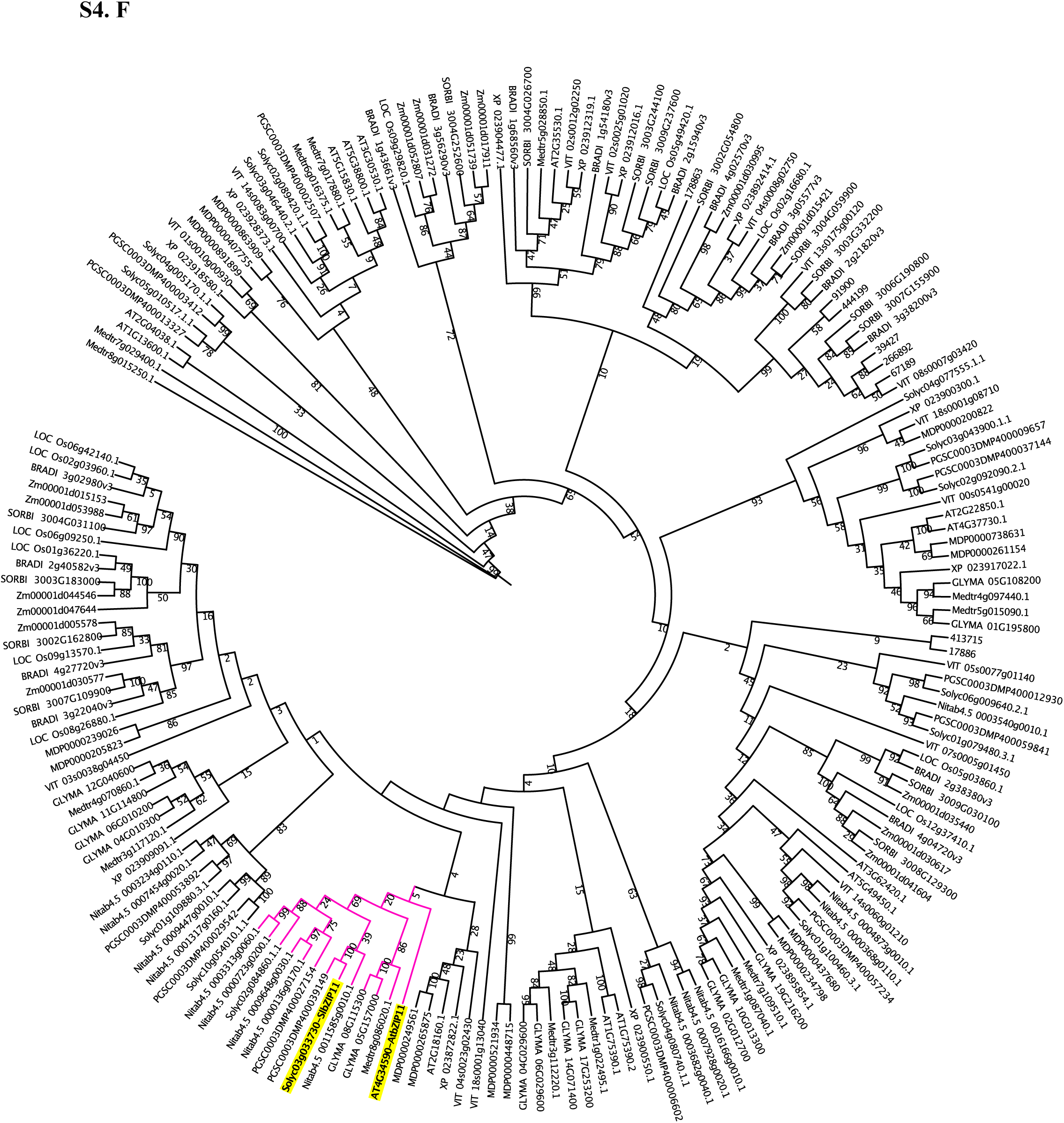
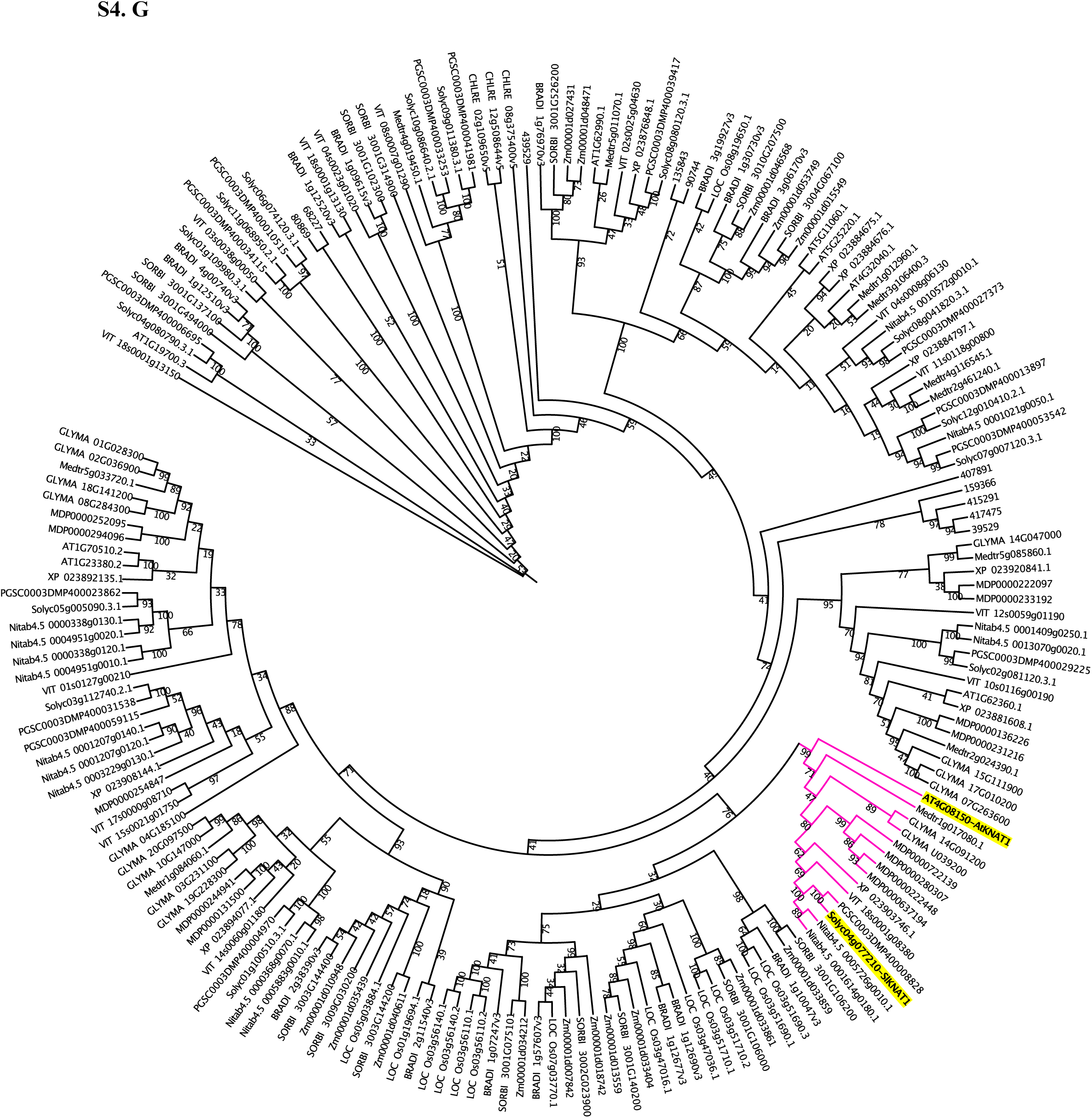
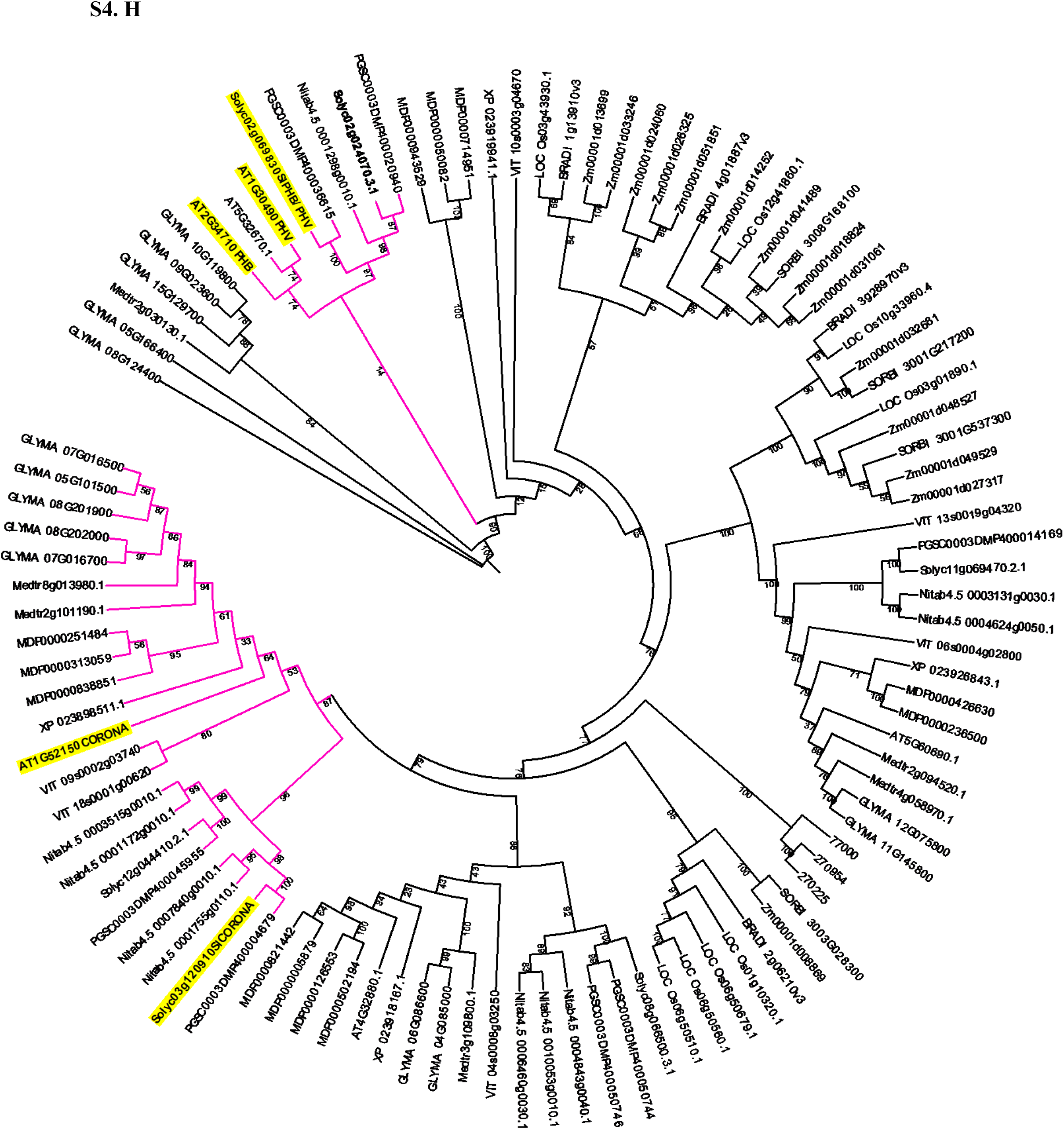
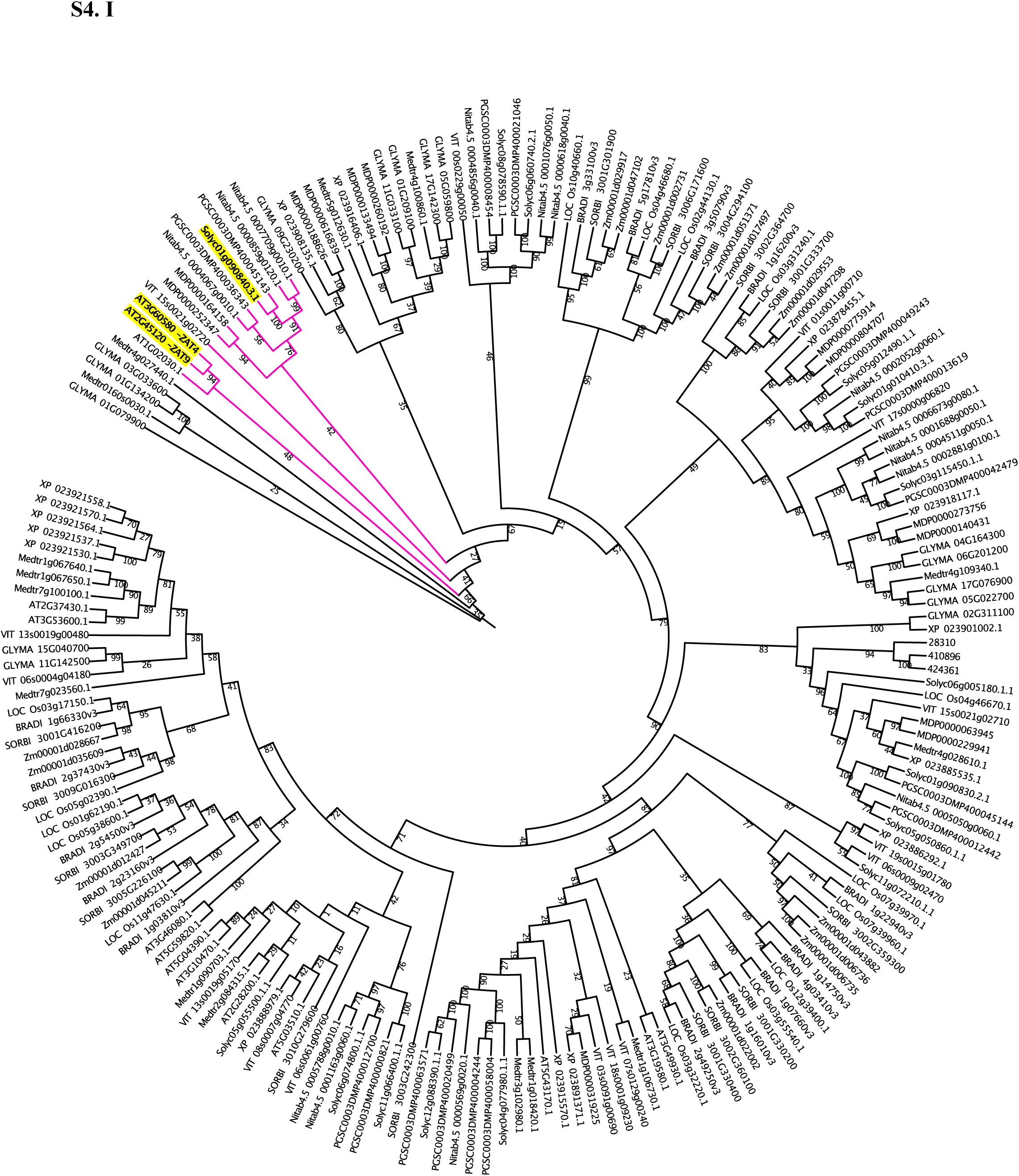
Gene family phylogenies. (**A-E**) Phylogenetic tree for 5 xylem enriched transcription factors that represent Solanaceae or Solanum expansion or tomato gene duplications. Clades/branches are highlighted in pink on the phylogenies and genes of interest highlighted in yellow. (**A**) *Solyc02g011680*/tomato duplication (**B**) *Solyc03g005350/*Solanaceae gene duplication (**C**)*Solyc06g072370/*Solanaceae-specific clade (**D**) *Solyc07g008500/*tomato duplication (**E**) *Solyc11g071500/*Solanum-expanded clade. (**F**) *SlbZIP11* (*Solyc03g033730),* (**G**) *SlKNAT1 (Solyc04g077210),* (**H**) HDZIPIII TF family, gene highlighted in green is another potential ortholog for *AtPHB/PHV* (**I**) *SlZAT4/9 (Solyc01g090840).* Phylogenetic trees were built using RAxML v8.2.12 (**Materials and Methods).** Legend: AT, *Arabidopsis thaliana*; BRADI, *Brachypodium distachyon;* CHLRE, *Chlamydomonas reinhardtii; GLYMA, Glycine max;* MDP, *Malus domestica*; Medtr, *Medicago truncatula*; Nitab, *Nicotiana_tabacum*; XP, *Quercus suber*; LOC_Os, *Oryza sativa*; Solyc, *Solanum lycopersicum*;; Smo, *Selaginella moellendorffii*; PGSC, *Solanum tuberosum*; SORBI, *Sorghum bicolor*; VIT, *Vitis vinifera*; ZM, *Zea mays*.

**Fig. S5.**
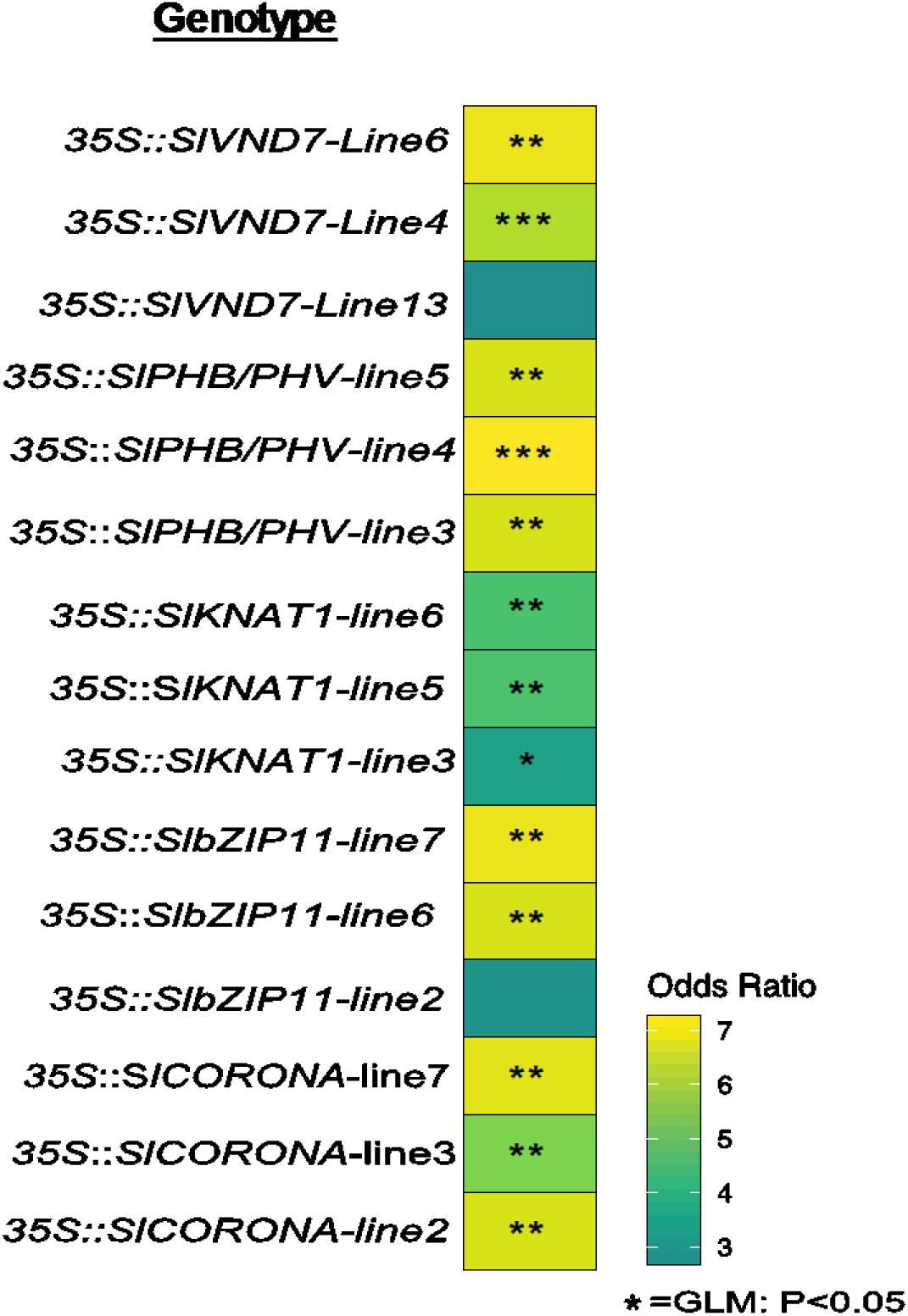
Quantification of abnormal xylem phenotypes in hairy root overexpression lines. Heatmap of log2 odds ratio of abnormal xylem phenotypes in 3 independent transgenic lines of *SlVND7, SlbZIP11*, *SlKNAT1, SlPHB/PHV* and *SlCORONA*. *n* = ∼15. See **Data S3** for all odds ratios and p-values.

**Fig. S6.**
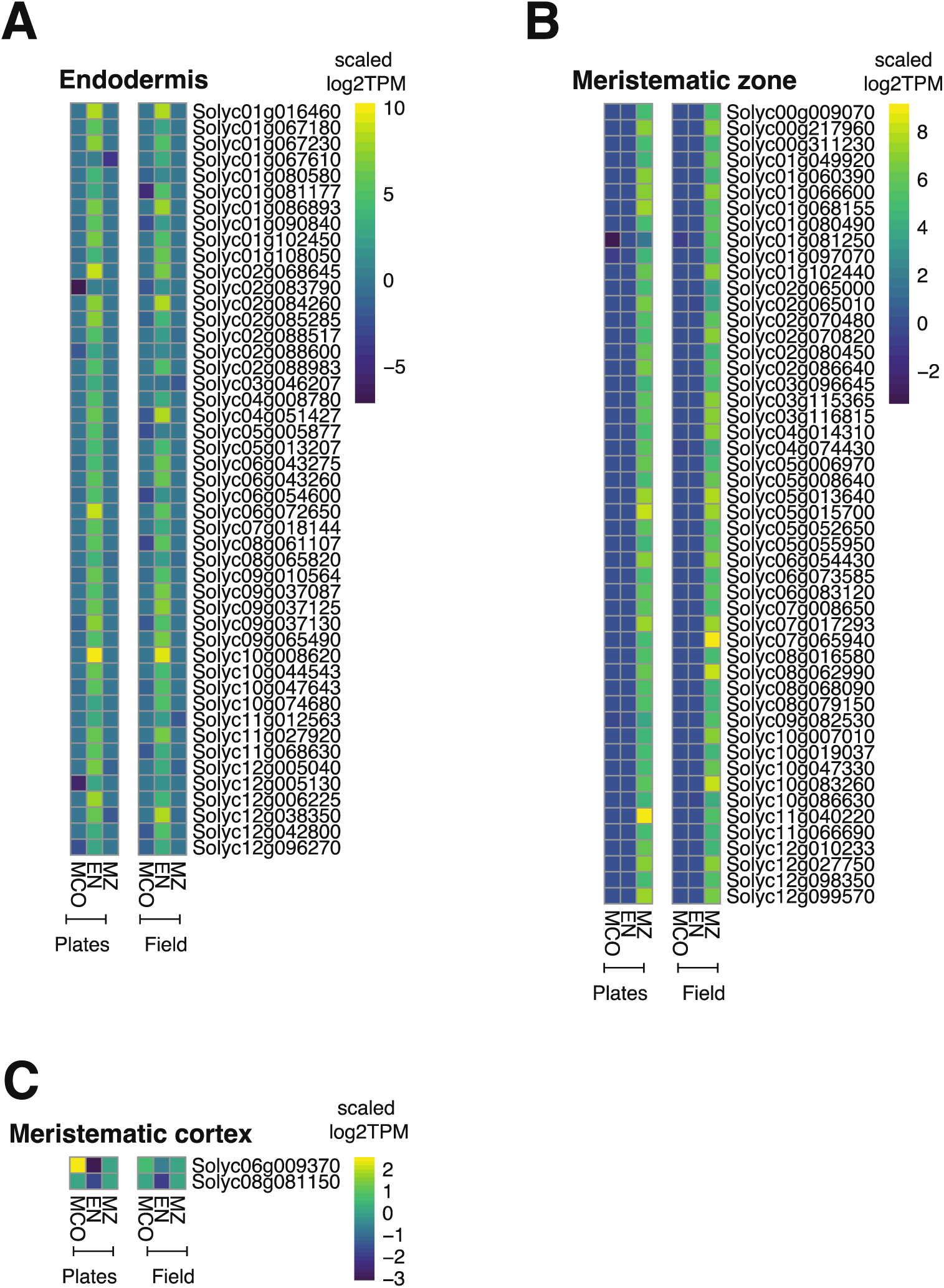
“Core” Cell Type-Enriched Genes. “Core” cell type-enriched gene ribosome-associated transcript abundance for the endodermis (**A**), meristematic zone (**B**) and cortex (**C**) in seedlings (plates) and the field.

**Fig. S7.**
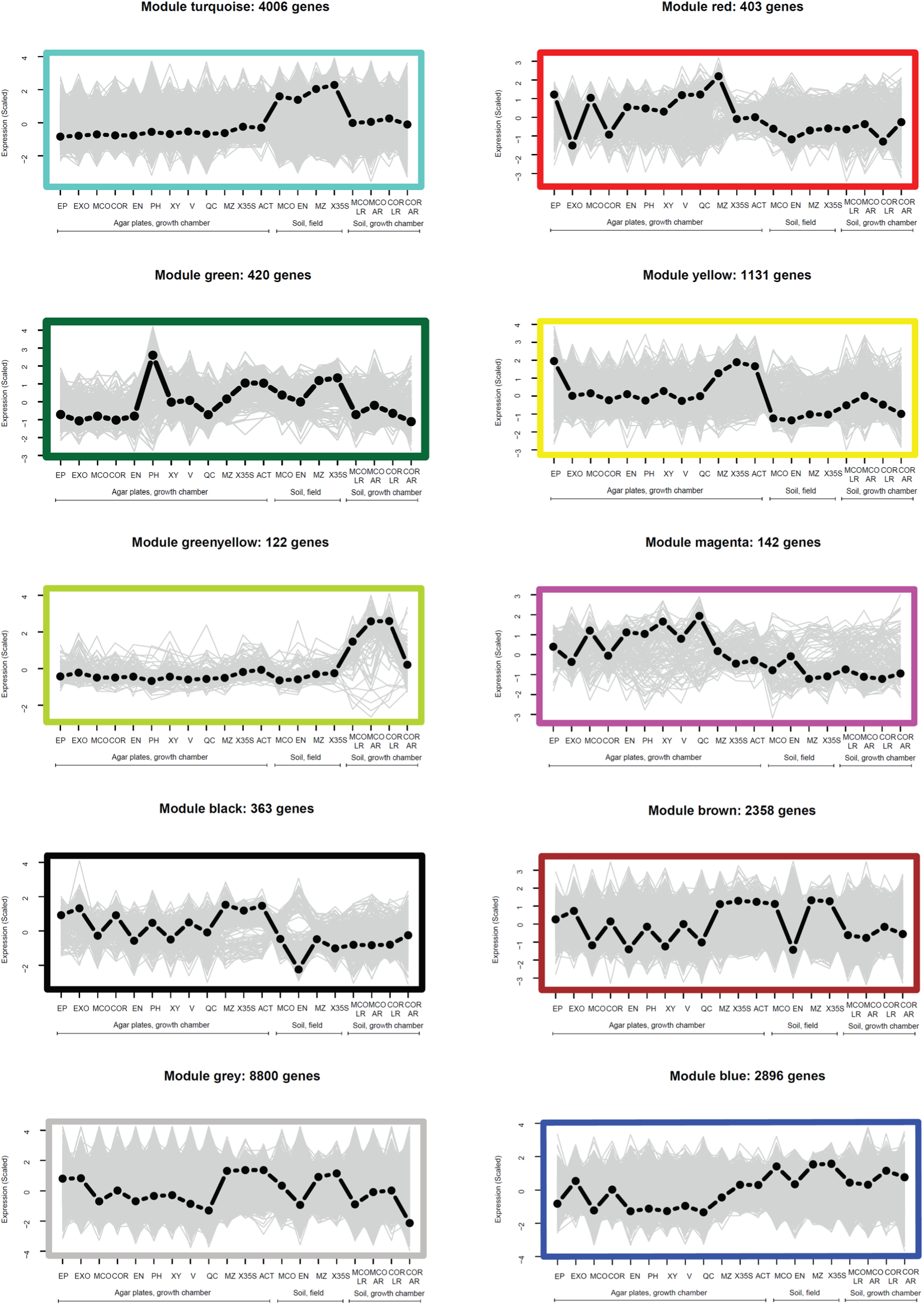
WGCNA Co-Expression Modules. Expression profiles of WGCNA co-expression modules not shown in **Fig. 3A-D**.

**Fig. S8.**
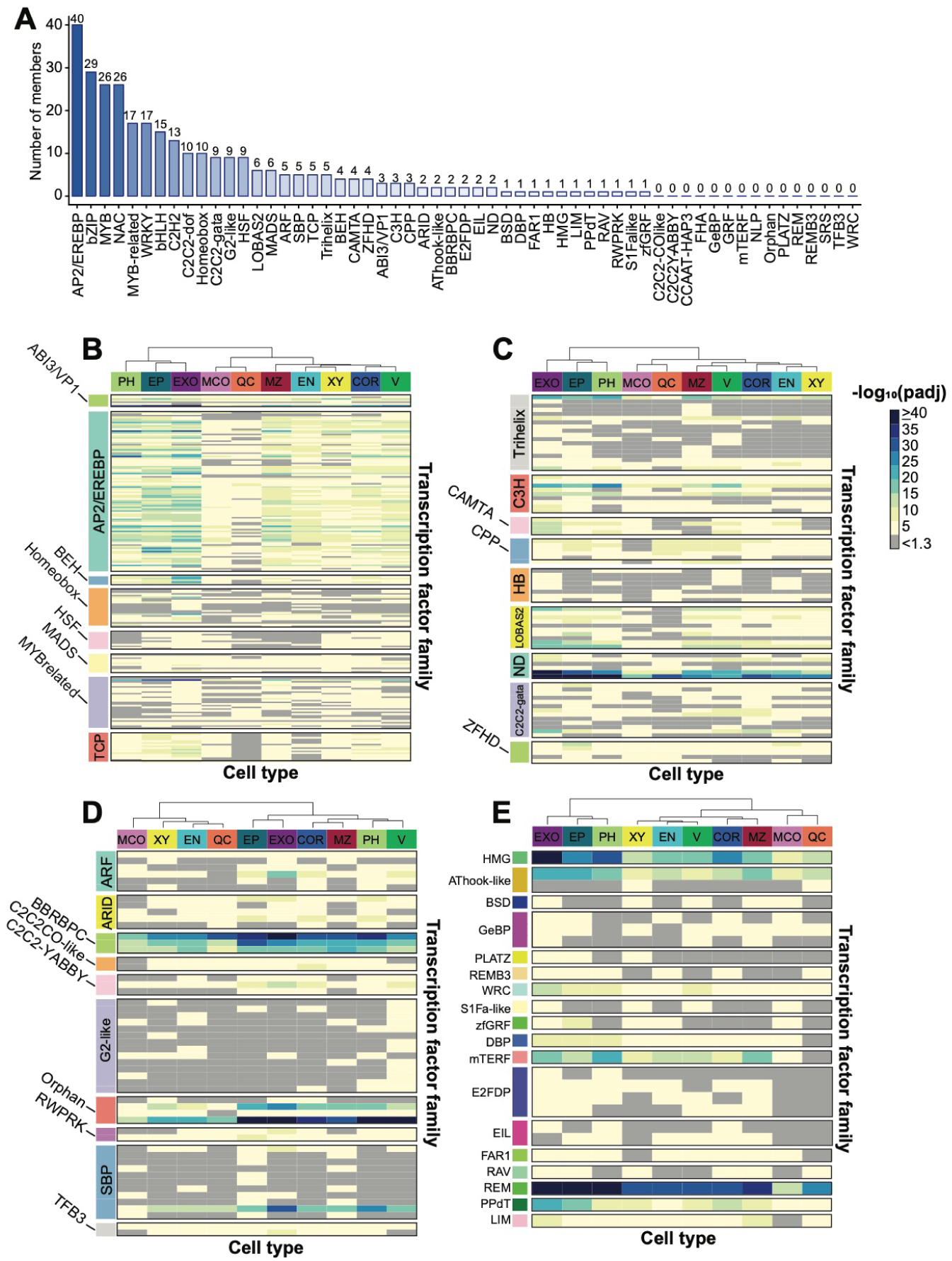
Transcription factor motifs enriched in 1-Kb promoters of cell type-enriched genes. (**A**) Histogram demonstrating the number of identified transcription factor motifs with expressologs in tomato. (**B-E**) All trees are hierarchically clustered to indicate similarity in enrichment across cell types. -log_10_ adjusted p-values are indicated according to the heatmap scale in the right part of the figure. (**B**) ABI3/VP1; BEH; Homeobox; Heat Shock Factor, MADS, and MYB-related transcription factor motif-enrichment. (**C**) Trihelix, C3H, CAMTA, CPP, Homeobox, LOB-AS2, C2C2-GATA and ZF-HD transcription factor motif enrichment. (**D**) ARF, ARID, BBRC/BPC; C2C2/CO-like; C2C2-YABBY, B2-like, Orphan, RWPRK, SBP and TFB3 transcription factor motif enrichment. (**E**) HMG, AThook-like, BSD, GeBP, PLATZ, REMB3, WRC, S1Fa-like, zfGRF, DBP, mTERF, E2FDP, EIL, FAR1, RAV, REM, PPdT, LIM transcription factor motif enrichment. COR=cortex; EN=endodermis; EP=epidermis; EXO=exodermis; MCO=meristematic cortex; MZ=meristematic zone; PH= phloem; V=vasculature; QC=quiescent center; XY= xylem.

**Fig. S9.**
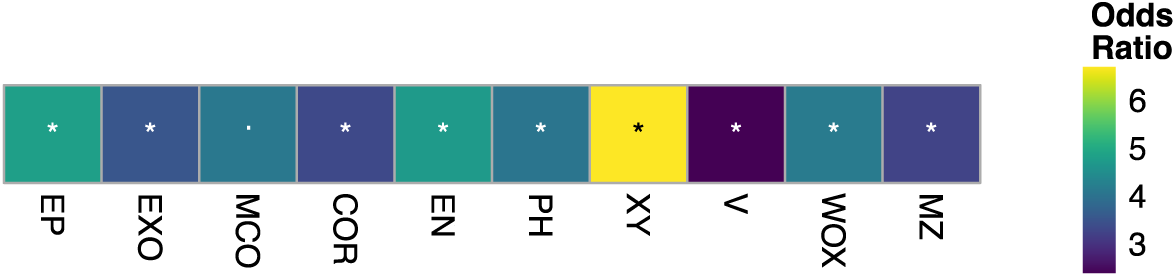
Enrichment of expressed genes near CTEARs. The heatmap summarizes the results from Fisher Exact Tests of expressed vs non-expressed genes near CTEARs (**Materials and Methods**). The color scale indicates the odds ratio. Symbols indicate level of significance (Asterisk denotes FDR < 0.05; dot denotes FDR = 0.05; Contingency matrices and results are in **Data S6**).

**Fig. S10.**
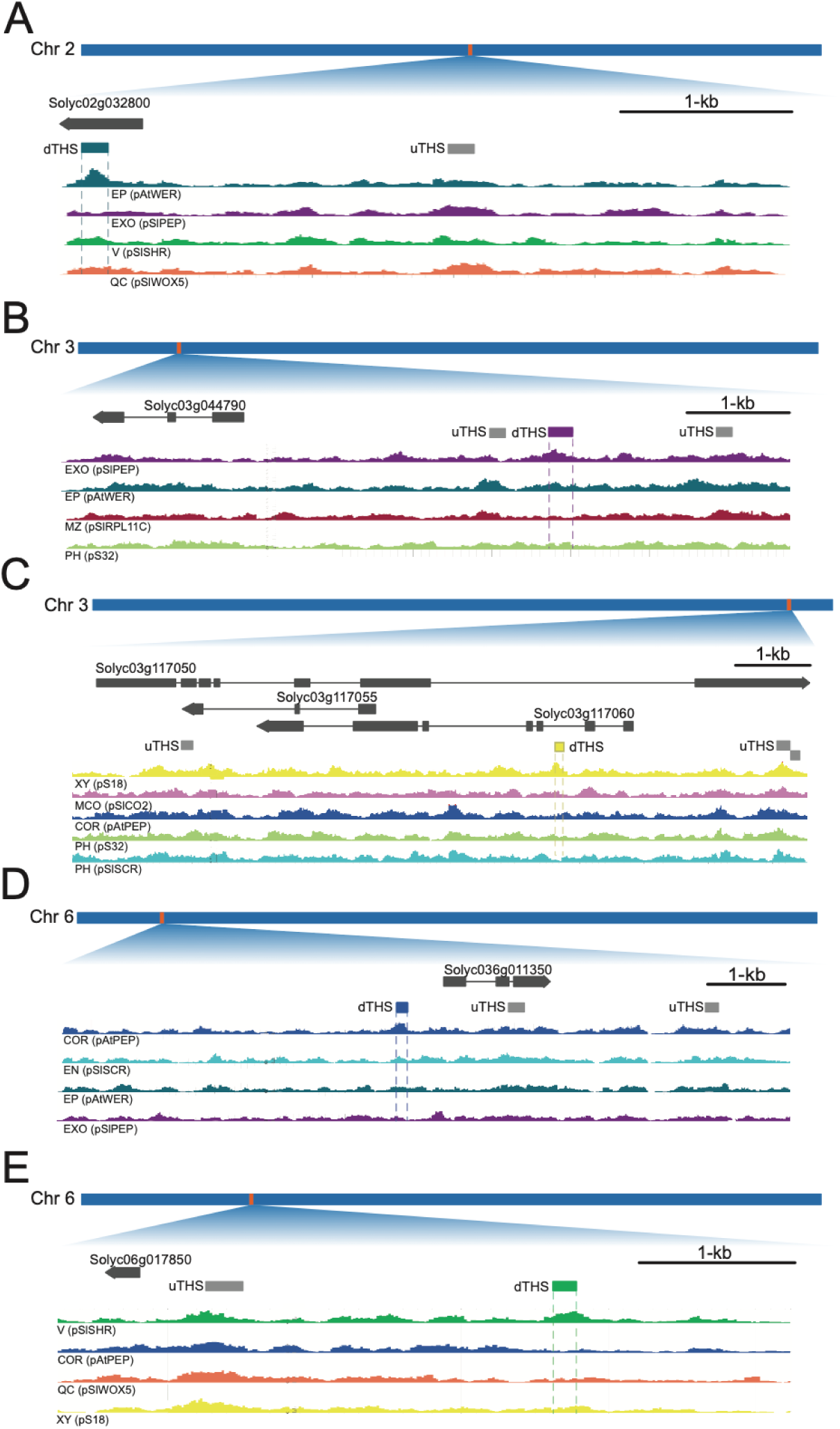
Representative browser shots for cell type-enriched accessible regions for the epidermis, exodermis, xylem, cortex and vasculature. (**A-E**) Y-axis=merged cut counts normalized with deeptools (Methods); x-axis = chromosome location; Gene models=top track; union THSs (uTHSs) are indicated with a grey rectangle while a cell type-enriched accessible regions (CTEARs) are indicated with a colored rectangles. COR=cortex; EN=endodermis; EP=epidermis; EXO=exodermis; MCO=meristematic cortex; MZ=meristematic zone; PH= phloem; V=vasculature; QC=quiescent center; XY= xylem. (**A**) CTEAR for epidermis. (**B**) CTEAR for exodermis. (**C**) CTEAR for xylem. (**D**) CTEAR for cortex. (**E**) CTEAR for vasculature.

**Fig. S11.**
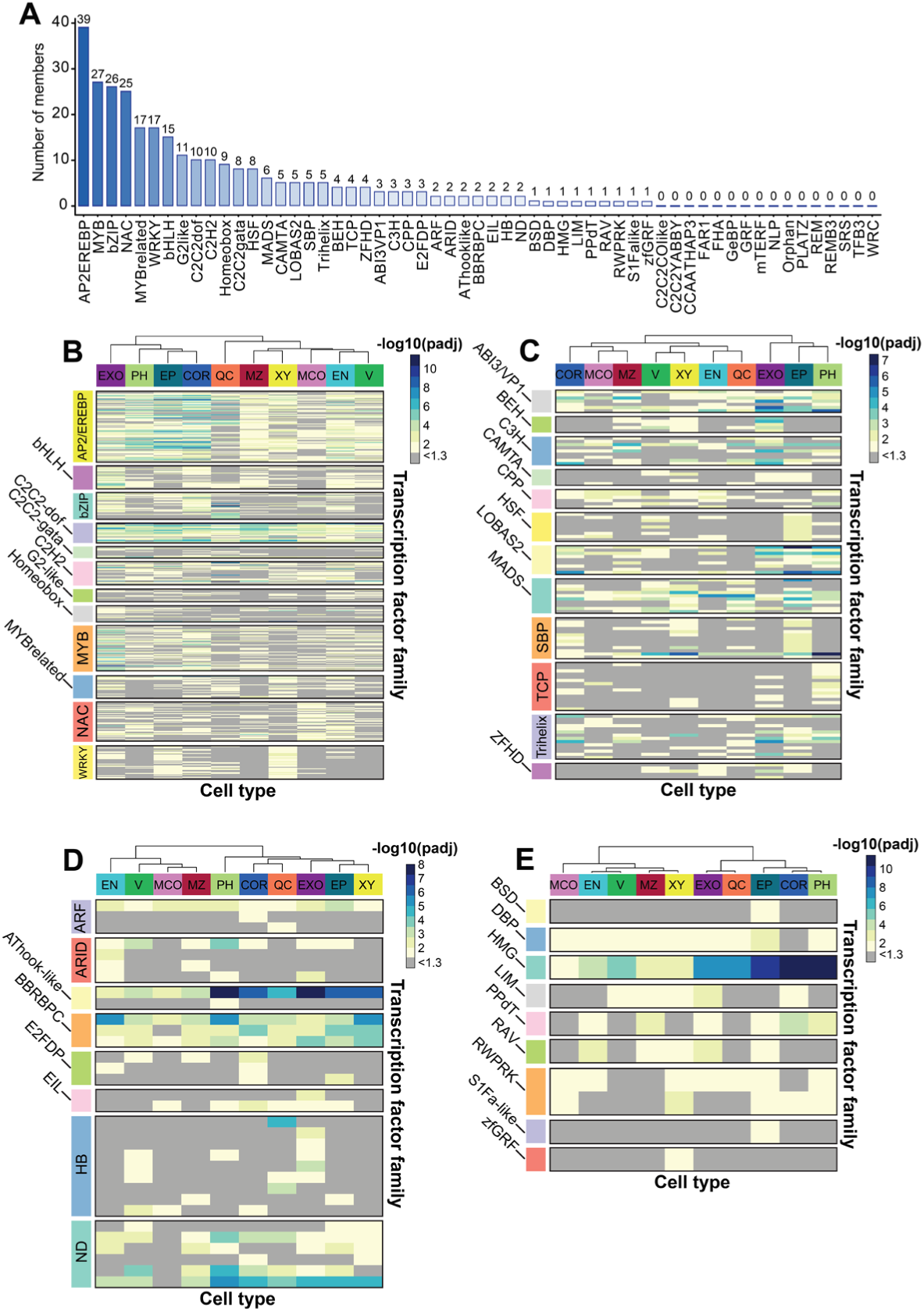
Transcription factor motifs enriched in accessible regions near cell type-enriched genes. Accessible regions 4-kb upstream of the transcription start site or 1-kb downstream of the transcription termination site of cell type-enriched genes were used to perform motif enrichment. (**A**) Histogram demonstrating the number of identified transcription factor motifs with expressologs in tomato. (**B-E**) All trees are hierarchically clustered to indicate similarity in enrichment across cell types. -log_10_ adjusted p-values are indicated according to the heatmap scale in the right part of the figure. (**B**) AP2/EREBP, bHLH, bZIP, C2C2-dof, C2C2-gata, C2H2, G2-like, MYB, MYB-related, NAC and WRKY transcription factor motif-enrichment. (**C**) ABI3/VP1, BEH, C3H, CAMTA, CPP, HSF, LOBAS2, MADS, SBP, TCP, Trihelix and ZFHD transcription factor motif enrichment. (**D**) ARF, ARID, AThook-like, BBRBPC, E2FDP, EIL, HB and ND transcription factor motif enrichment. (**E**) BSD, DBP, HMG, LIM, PPdT, RAV, RWPRK, SF1A-like and zfGRF transcription factor motif enrichment. COR=cortex; EN=endodermis; EP=epidermis; EXO=exodermis; MCO=meristematic cortex; MZ=meristematic zone; PH= phloem; V=vasculature; QC=quiescent center; XY= xylem.

**Fig. S12.**
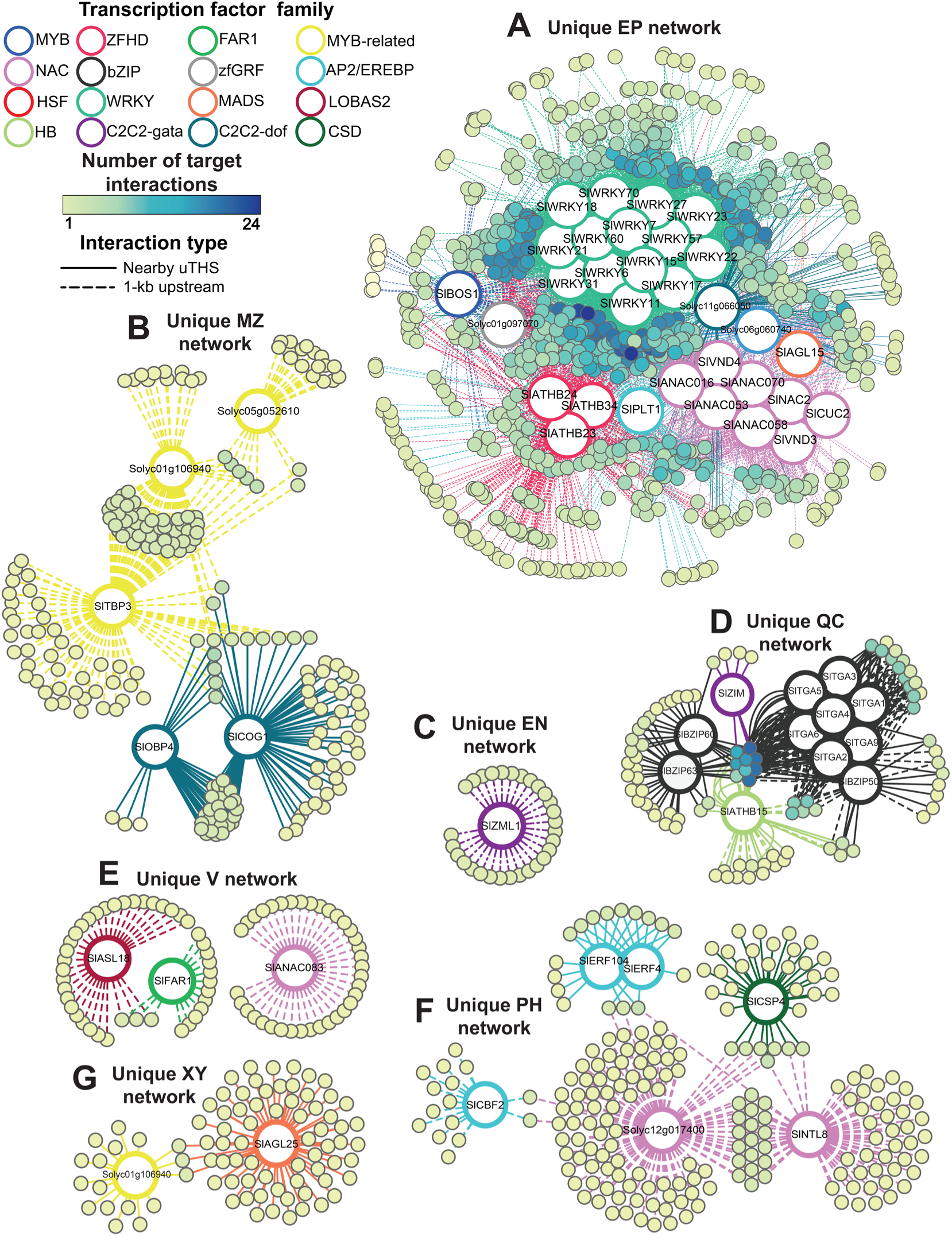
Inferred unique cell type transcriptional regulatory networks. (A-G) Solid edges. = motif-uTHS interaction; dashed edges = motif-1kb upstream regulatory region interaction; large circles = TF expressolog for cognate TF motif; colored edges indicate transcription factor family. Small circles = target genes which contain the motif in either the uTHS or 1kb upstream regulatory region; color scale indicates the number of target interactions. COR=cortex; EN=endodermis; EP=epidermis; MZ=meristematic zone; PH=phloem; V=vasculature; XY=xylem; QC=quiescent center.

**Fig. S13.**
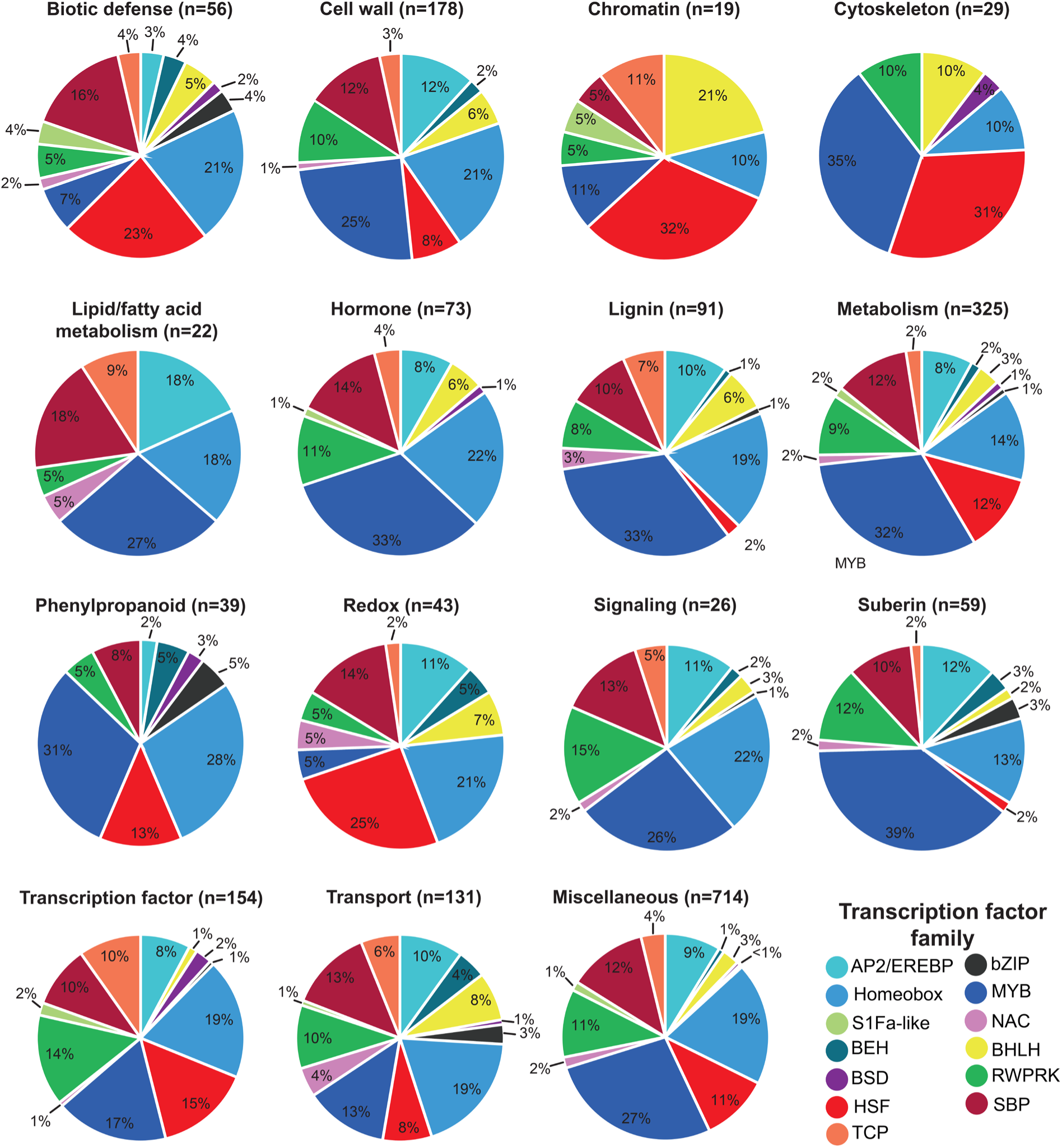
Exodermis regulatory modules based on ITAG3.2 gene annotation. Targets in the unique exodermis network were manually annotated for the following categories of interest: biotic defense, cell wall, chromatin, cytoskeleton, lipid/fatty acid metabolism, hormone, lignin, metabolism, phenylpropanoid, redox, signaling, suberin, transcription factor, transport, and miscellaneous. Color key for the pie charts denotes the family for the transcription factor motifs.

**Fig. S14.**
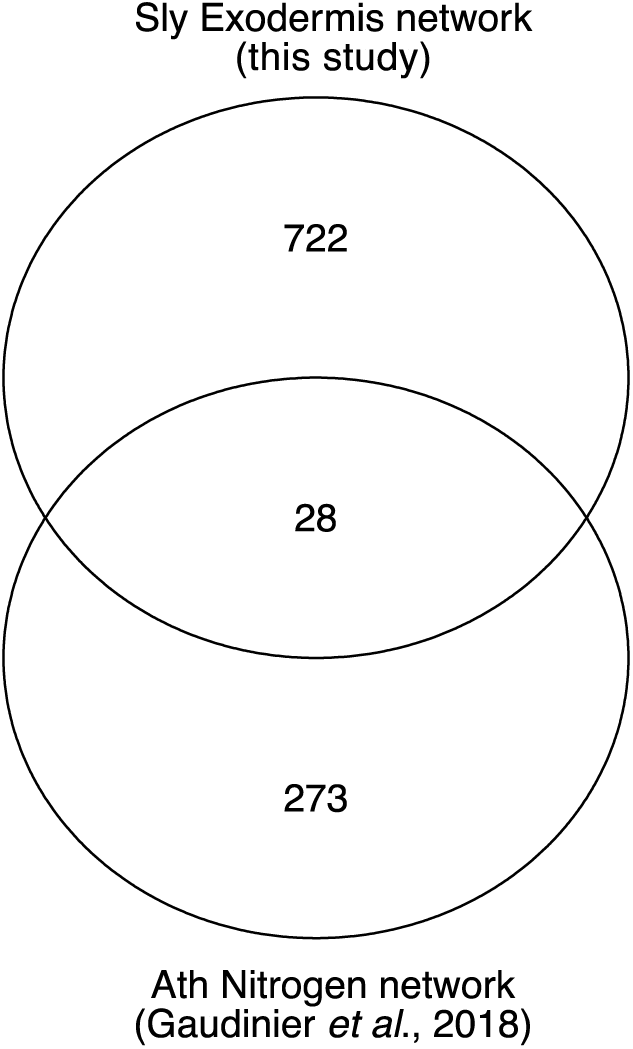
Overlap of Arabidopsis nitrogen metabolism network with tomato exodermis-specific network. Figure shows the number of tomato expressologs from an Arabidopsis nitrogen metabolism-related network (*27*) and genes in our exodermis-specific network.

**Fig. S15.**
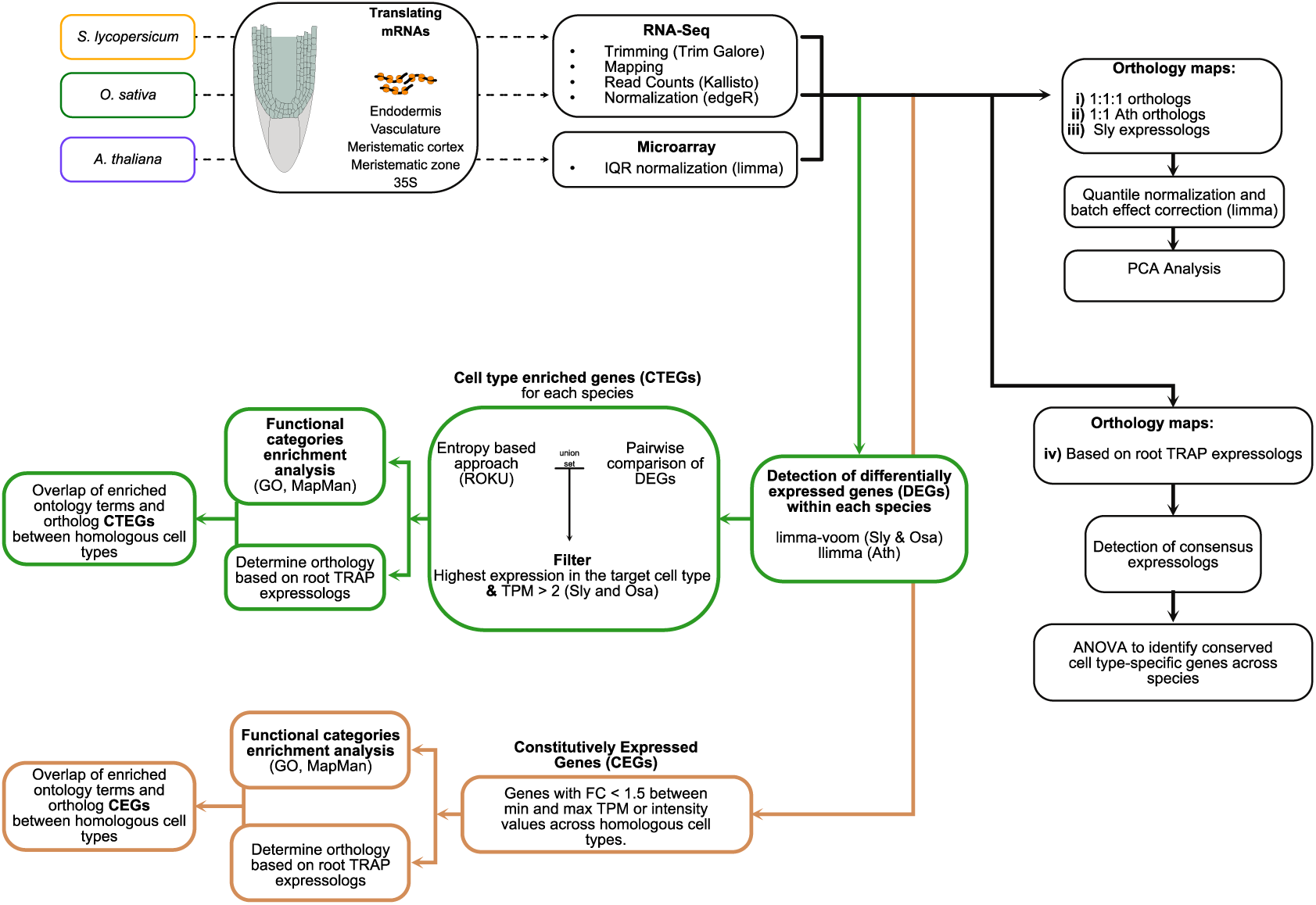
Flowchart for multispecies analysis. An overview of the pipeline and methods used for exploring the conservation of homologous cell type and tissues among tomato, Arabidopsis and rice. Ath=*Arabidopsis thaliana*; Osa=*Oryza sativa*; Sly=*Solanum lycopersicum*; IQR= interquartile range; TPM=transcript per million; GO=gene ontology; CTEG=cell type/tissue enriched genes.

**Fig. S16.**
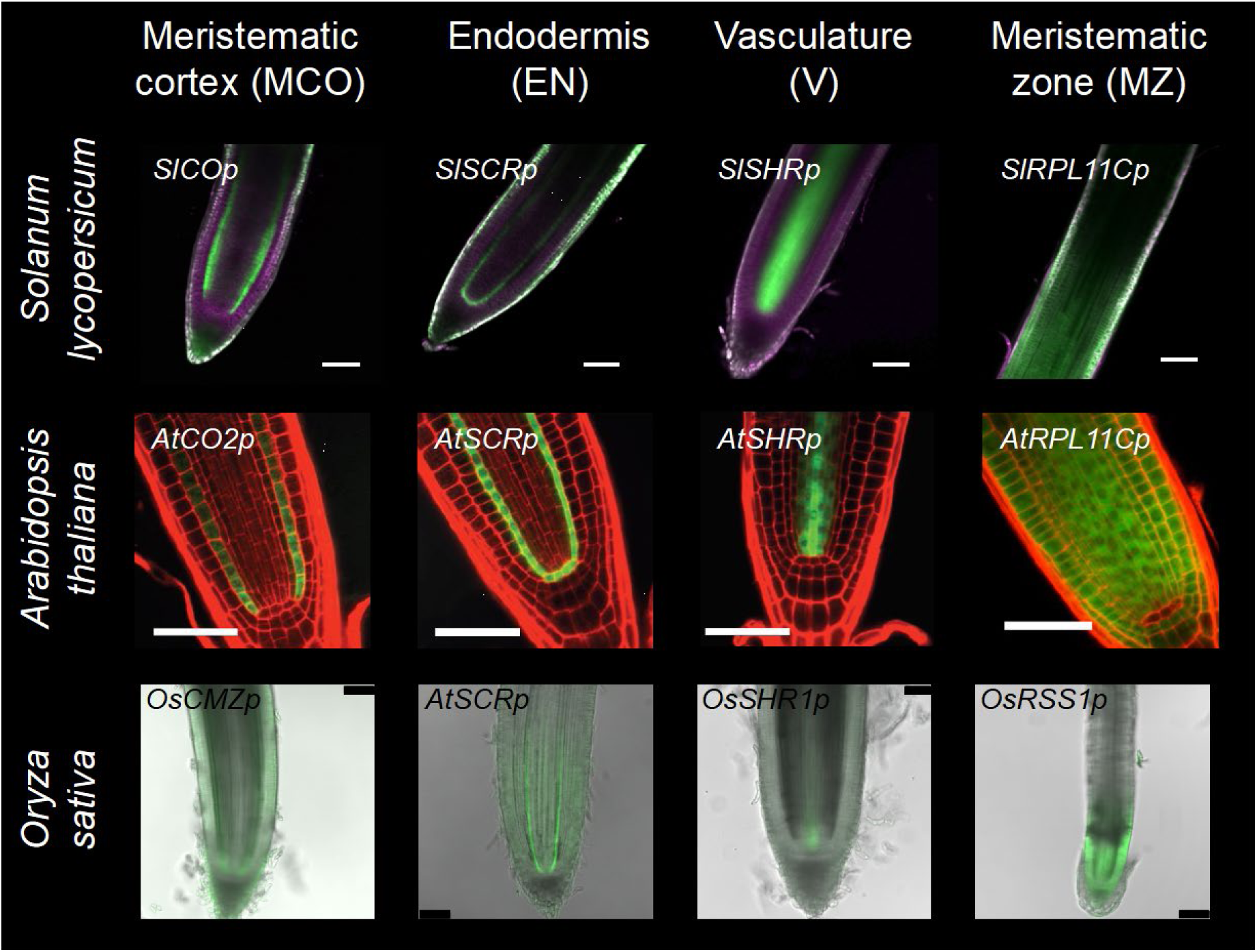
Promoters driving expression in potentially homologous cell types of tomato, rice and Arabidopsis. Expression patterns of GFP (green color) in the TRAP marker lines selected for the multi-species analysis. The promoters driving GFP expression are indicated for each line. Magenta color denotes autofluorescence for tomato and red color denotes propidium iodide staining for Arabidopsis. Arabidopsis data is adapted from (*5*). Scale bars represent 100µm for tomato, 50µm for Arabidopsis and 100µm for rice.

**Fig. S17.**
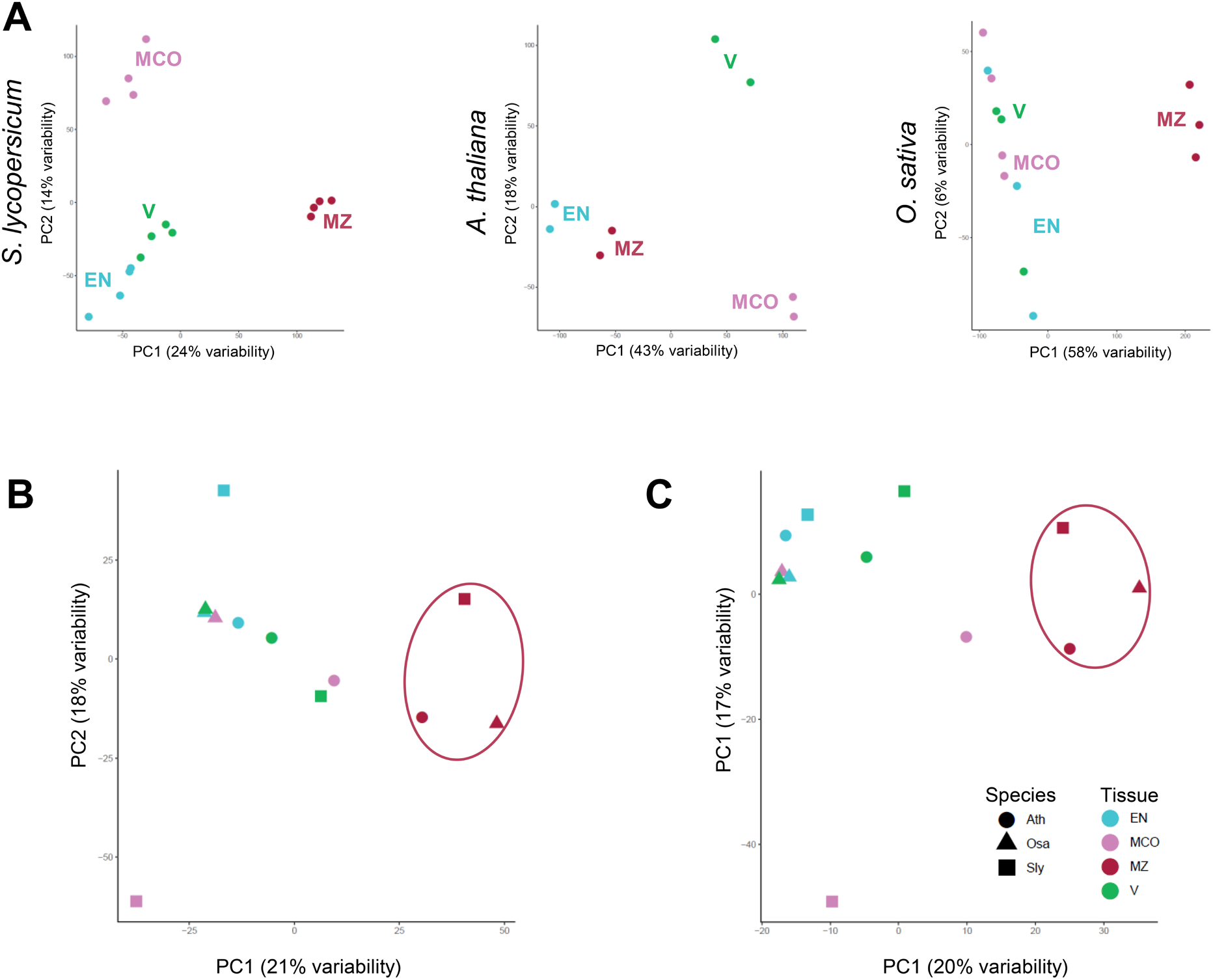
PCA Plots with Independently Derived Orthology Maps Demonstrate Similarity Across Species for Some Tissue Types. (**A**) Clustering of cell type and tissue expression profiles (batch-corrected log_2_ normalized CPM values) within tomato, Arabidopsis and rice using principal component analysis (PCA). (**B**) and (**C**) Clustering of cell type/tissue expression profiles between Arabidopsis (circle), rice (triangle) and tomato (square) using two independently derived orthology maps. (**B**) PCA plot of cell type/tissue expression of 3,505 1:1 orthologs, based on sequence homology to Arabidopsis, as determined by MapMan annotation files of tomato and rice. (**C**) PCA plot of cell type and tissue expression of 1,771 Arabidopsis and rice expressologs of tomato with an expression correlation coefficient > 0.6. Ath=*Arabidopsis thaliana*; Osa=*Oryza sativa*; Sly=*Solanum lycopersicum*; EN=endodermis; MCO=meristematic cortex; MZ=meristematic zone; V=vasculature.

**Fig. S18.**
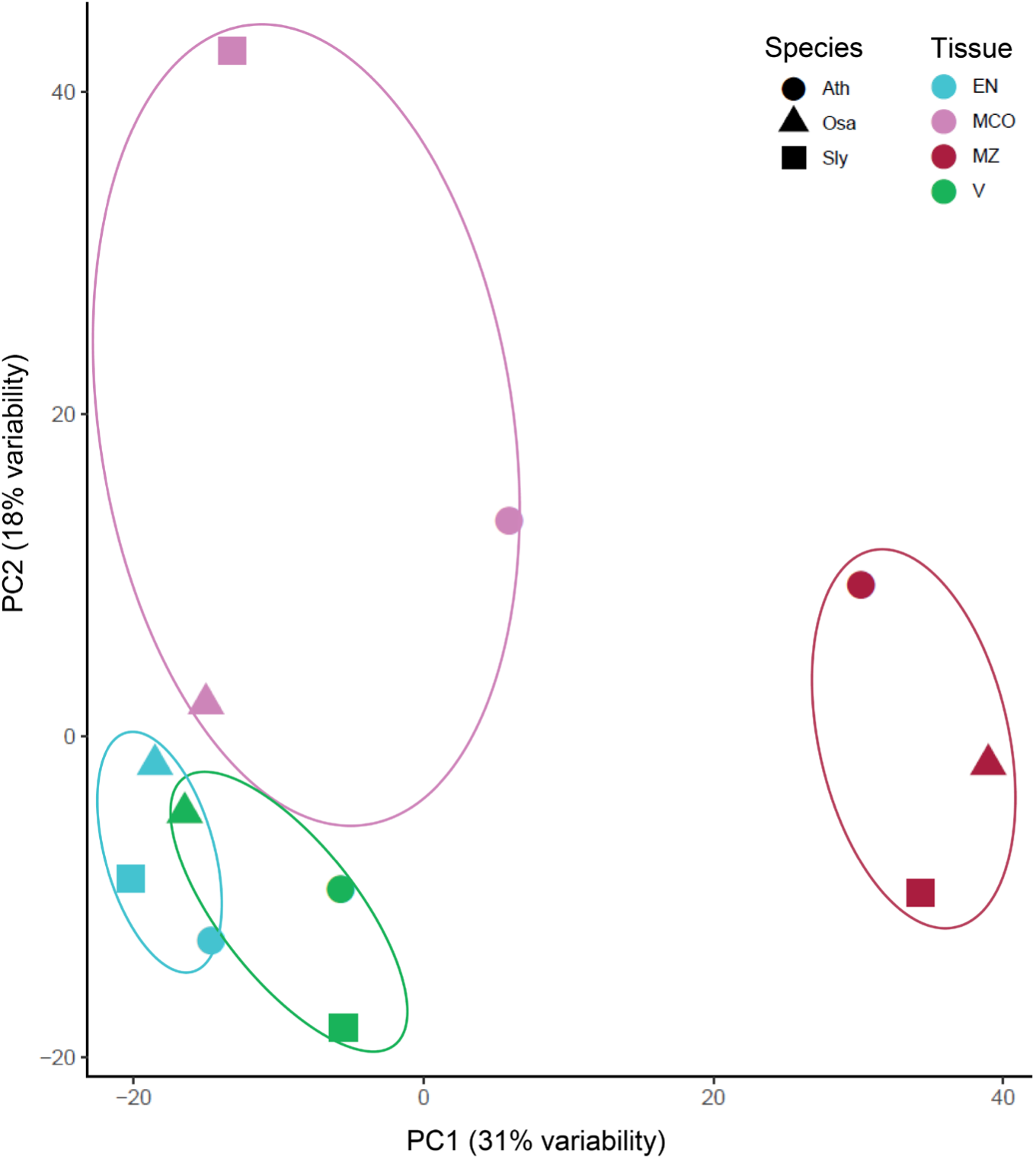
A Principal Component (PC) analysis of consensus expressolog expression. Clustering of cell type and tissue expression profiles of 1,585 consensus root TRAP expressologs between Arabidopsis (circle), rice (triangle) and tomato (square). Consensus expressologs have identical expressolog relationships independent of the reference species and positive expression correlations. Ath=*Arabidopsis thaliana*; Osa=*Oryza sativa*; Sly=*Solanum lycopersicum*; EN=endodermis; MCO=meristematic cortex; MZ=meristematic zone; V=vasculature.

**Fig. S19.**
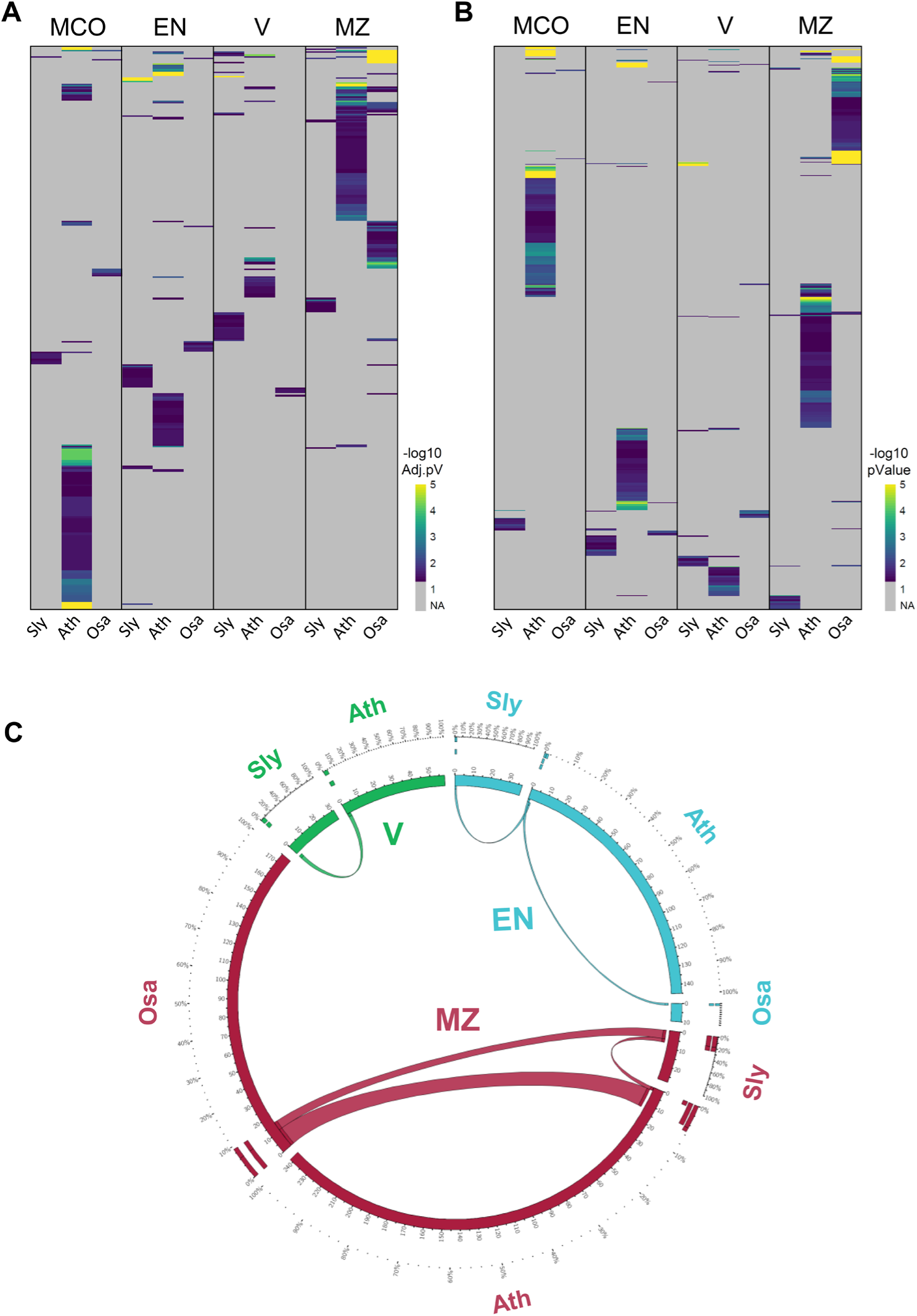
Ontology Term Enrichment within and between Species. (**A**) Overview of enriched MapMan and (**B**) Gene Ontology (GO) terms within and between species. (**C**) A circos plot indicating overlapping GO terms of homologous cell type/tissue between species. The width of the ribbon is proportional to the number of common terms. The numbers in the inner circle represent the number of terms within each group. The numbers in the outer circle represent the percentage of common terms. Ath=*Arabidopsis thaliana*; Osa=*Oryza sativa*; Sly=*Solanum lycopersicum;* EN=endodermis; MCO=meristematic cortex; MZ=meristematic zone; V=vasculature.

**Fig. S20.**
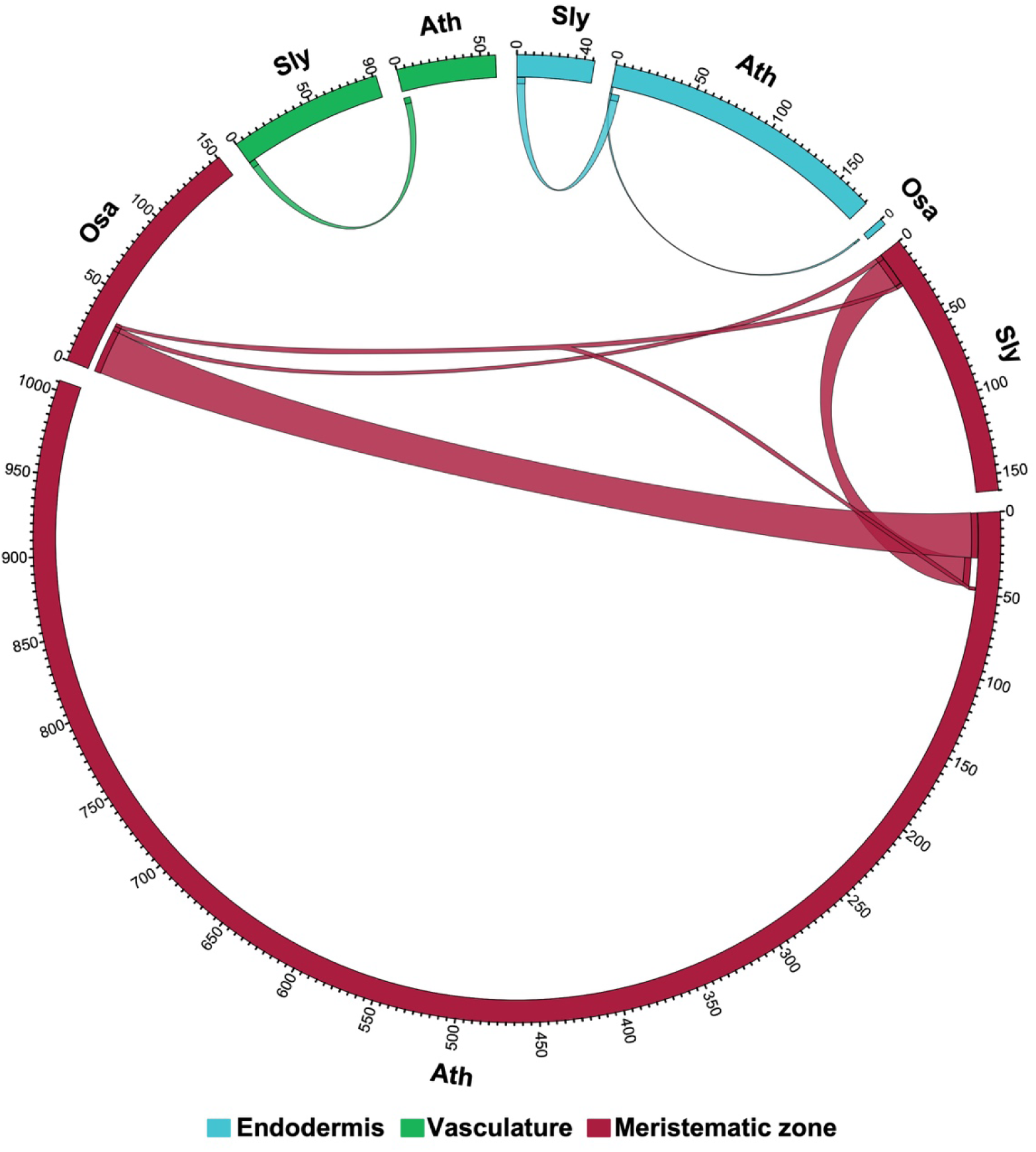
Circos Plot Indicating Overlapping Expressologs of Homologous CTEGs between Species. Ontology was determined based on 7,295 tomato and 6,424 rice root TRAP expressologs that have a reciprocal match and a positive expression correlation with Arabidopsis as a reference species. The width of the ribbon is proportional to the number of common expressologs. The numbers in the inner circle represent the number of expressologs within each group. The numbers in the outer circle represent the percentage of common expressologs. Ath=*Arabidopsis thaliana*; Osa=*Oryza sativa*: Sly=*Solanum lycopersicum;* EN=endodermis; MCO=meristematic cortex; MZ=meristematic zone; V=vasculature.

**Fig. S21.**
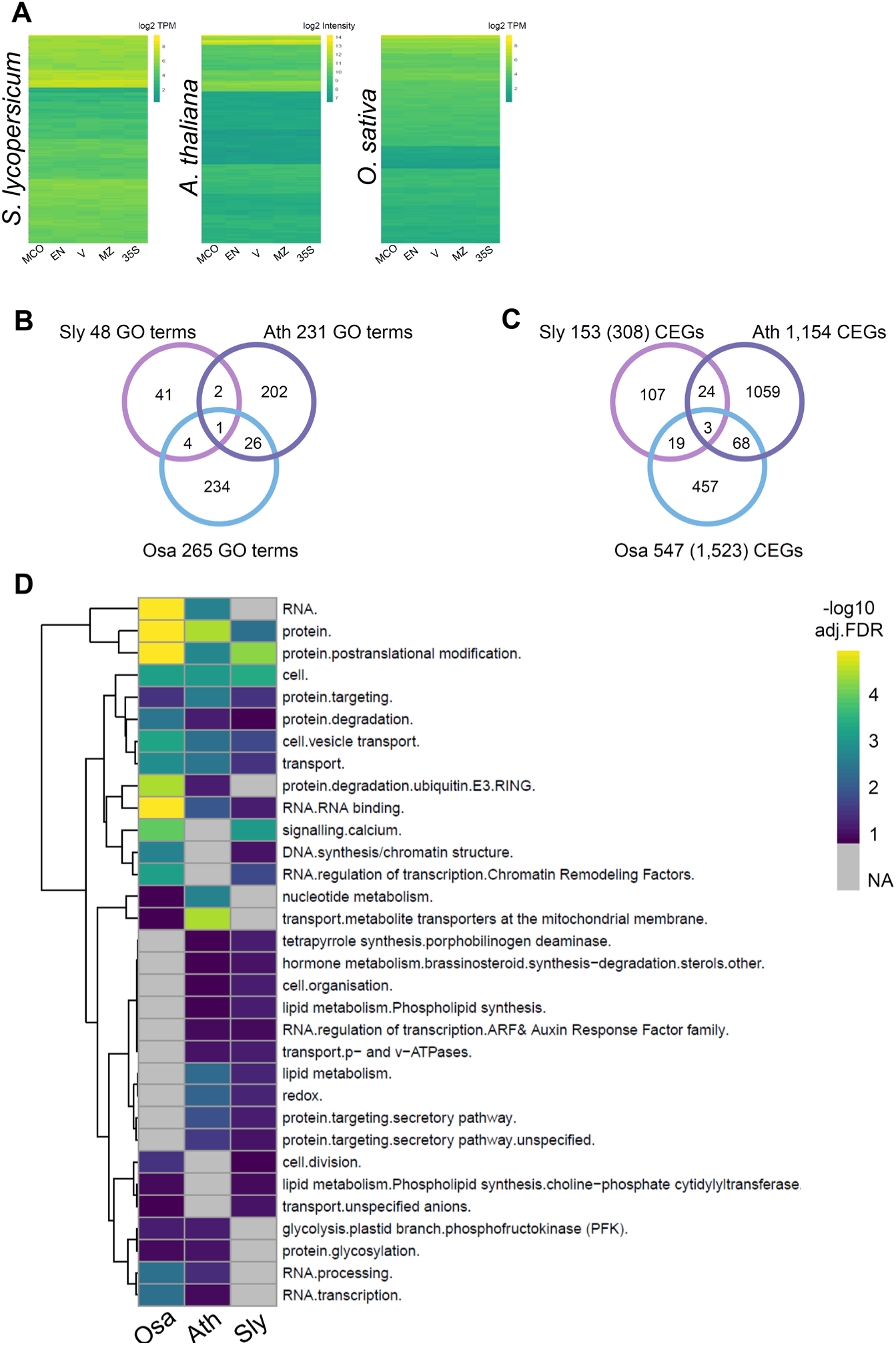
Overlap of Expressologs and Ontology Terms of Constitutively Expressed Genes (CEGs) Between Species. (**A**) Expression patterns of CEGs within each species. (**B**) Venn diagram of common and unique Arabidopsis root TRAP expressologs. Orthology was determined based on 7,295 tomato and 6,424 rice root TRAP expressologs that have a reciprocal match and a positive expression correlation with Arabidopsis. Numbers in parenthesis indicate the original number of CEGs detected within tomato and rice. (**C**) Overlapping MapMan terms between the CEGs. Ath=*Arabidopsis thaliana*; Osa=*Oryza sativa*: Sly=*Solanum lycopersicum*.

**Fig. S22.**
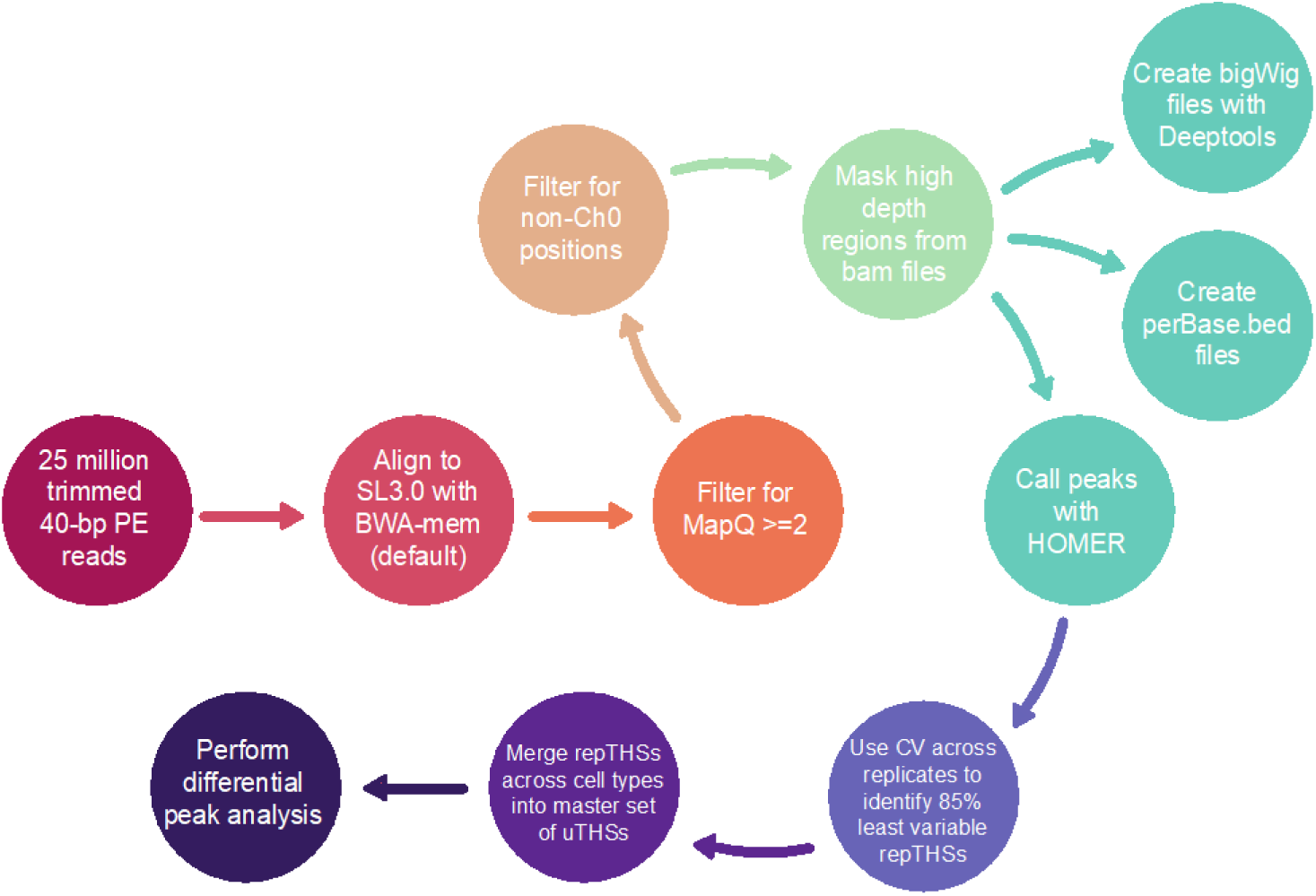
Flowchart for identification and annotation of ATAC-seq data. Data analysis overview for methods used for identification of replicate transposase hypersensitive sites.

**Fig. S23.**
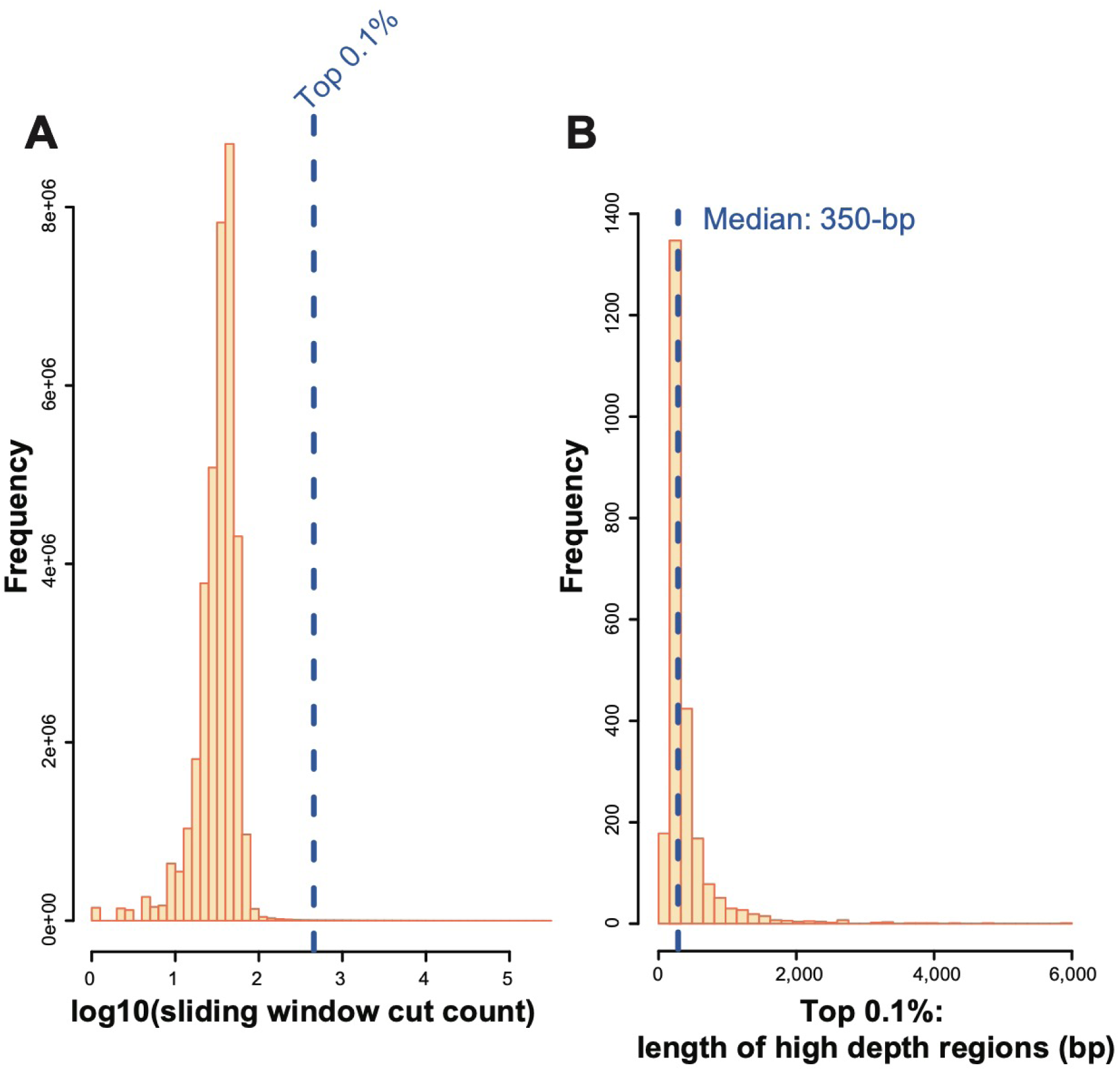
Identification of High Depth Sequencing Regions. (**A**) Cut counts from genomic DNA-based ATAC-seq were tallied across 150-bp sliding windows (step size 20-bp). X-axis, log_10_ number of reads in window. Y-axis: frequency. The blue dashed line represents the top 0.1% most accessible windows. (**B**) X-axis: length of top 0.1% high depth sequencing regions. Y-axis: frequency. This graph demonstrates the distribution of sizes for these high sequencing depth regions. Blue dashed line represents median high depth sequencing region size.

**Fig. S24.**
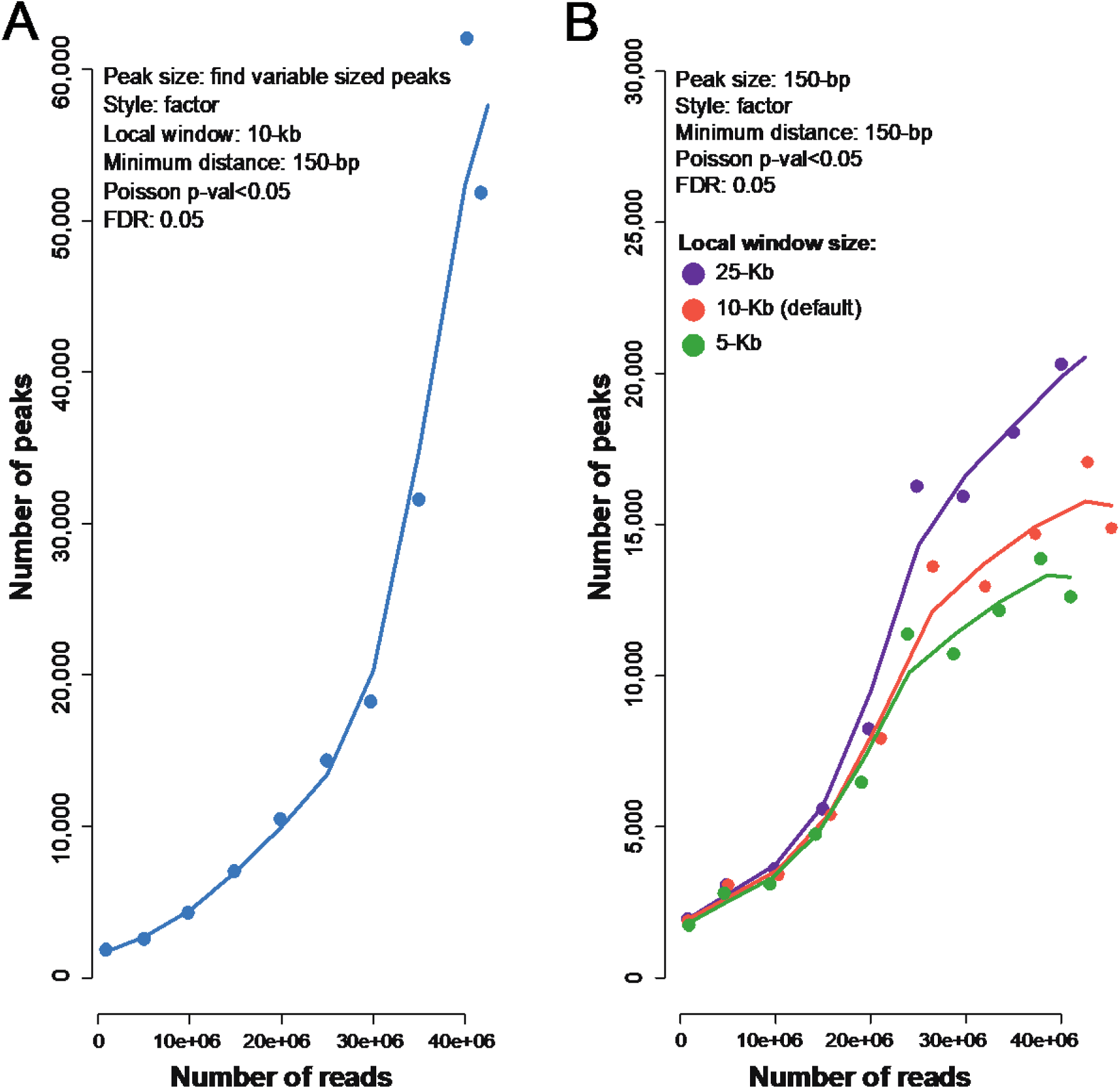
Choice of window size parameter for ATAC peak calling. (**A**) Peaks were called with increasing numbers of sampled reads from the sample WOX_O08. Here, the HOMER findPeaks parameters “-style factor”, “-minDist 150”,“-region” and “-regionRes 1”. X-axis: number of reads used to call peaks. Y-axis: number of peaks discovered. (**B**) Peaks were identified with HOMER using three different window sizes as well as the parameters “-size 150”, “-minDist 150” “-region” and “-regionRes 1”. X-axis: number of reads used to call peaks. Y-axis: number of peaks discovered.

**Fig. S25.**
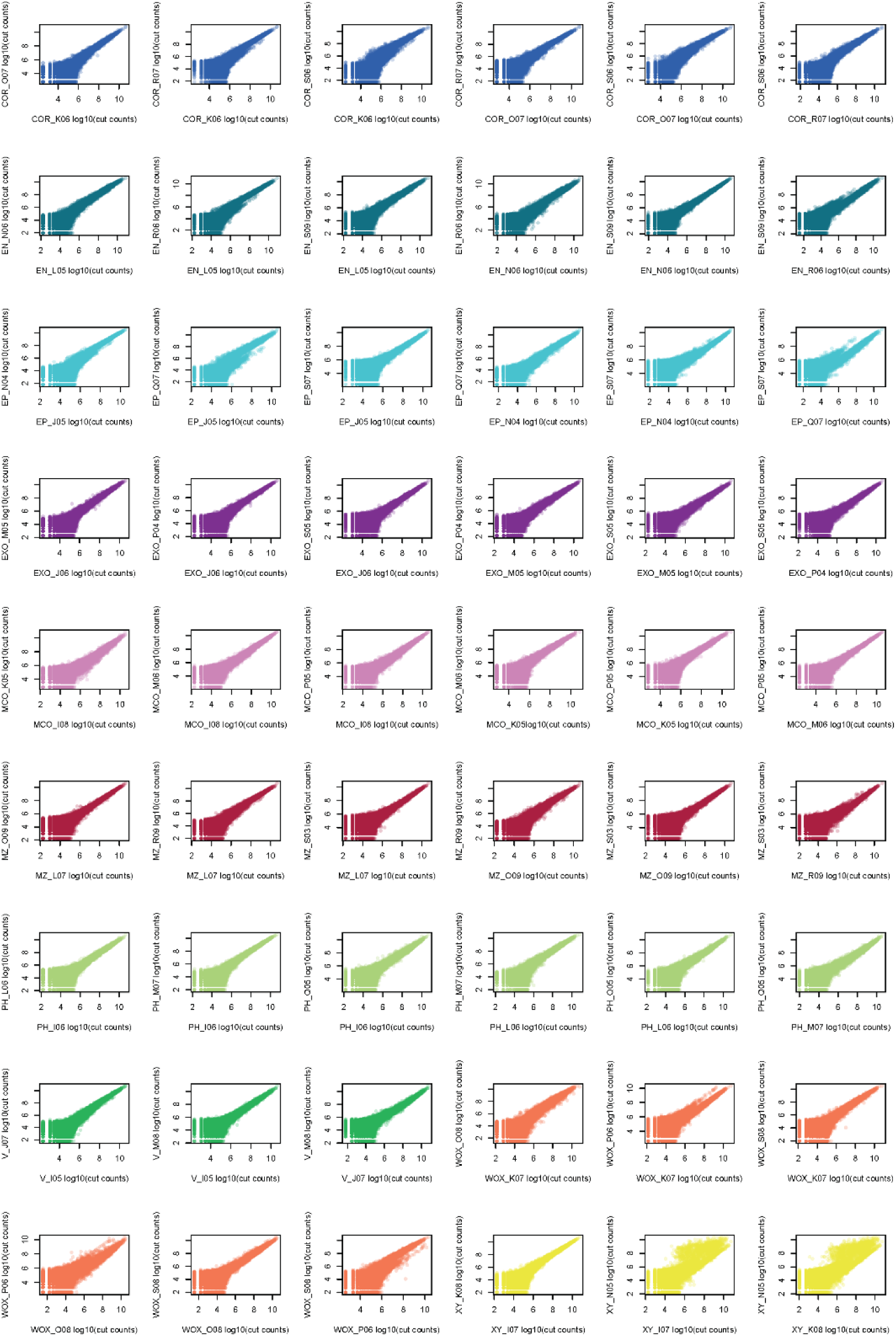
Pairwise comparison of ATAC cut-counts between cell types. Scatter plots for each pairwise comparison of replicates used within a cell type group. THSs discovered within a cell type group were merged and then used to tally the number of cut counts at that THS in each replicate. X-axis: log_10_ cut counts within a given replicate. Y-axis: log_10_ cut counts within a separate replicate. Cell type groups refer to promoters used INTACT lines (**Data S1**): COR (*pAtPEP*), EN (*pSlSCR*); EP (*pAtWER*); EXO (*pSlPEP*); MCO (*pSlCO2*); MZ (*pSlRPL11C*); PH (*pS32*); V (*pSlSHR*); WOX (*pSlWOX5*); XY (*pS18*).

**Fig. S26.**
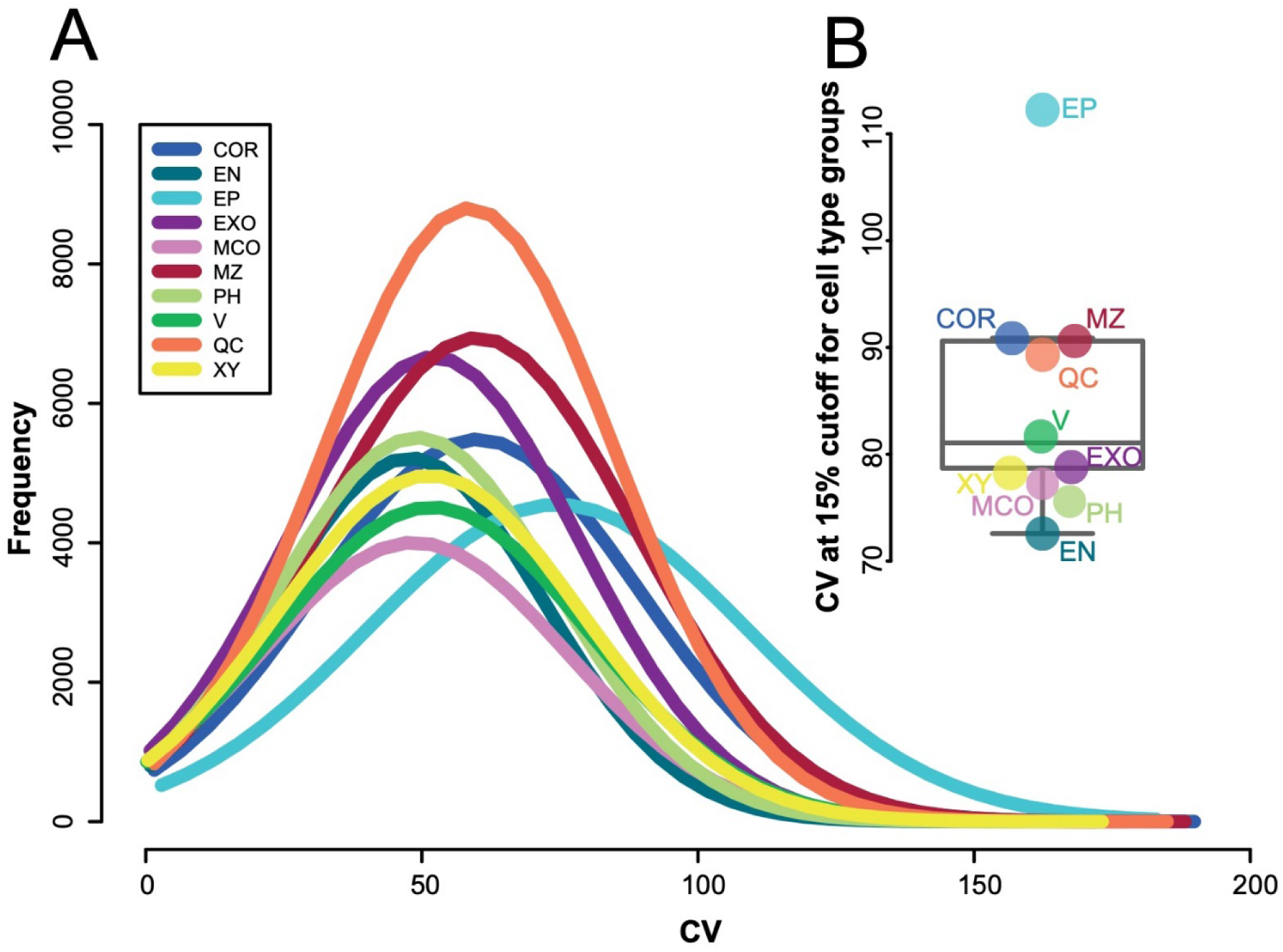
Threshold for identification of 15% most variable THSs for removal. THSs discovered within a cell type group were merged and then used to tally the number of cut counts at that THS in each replicate. The coefficient of variation (CV) for cut counts was calculated at each replicate THS across all replicates. (**A**) X-axis: CV across the replicate THSs for each cell type (see color legend) prior to filtering. (**B**) Boxplot of CV cutoff values for the top 15% most variable replicate THSs for each cell type.

**Fig. S27.**
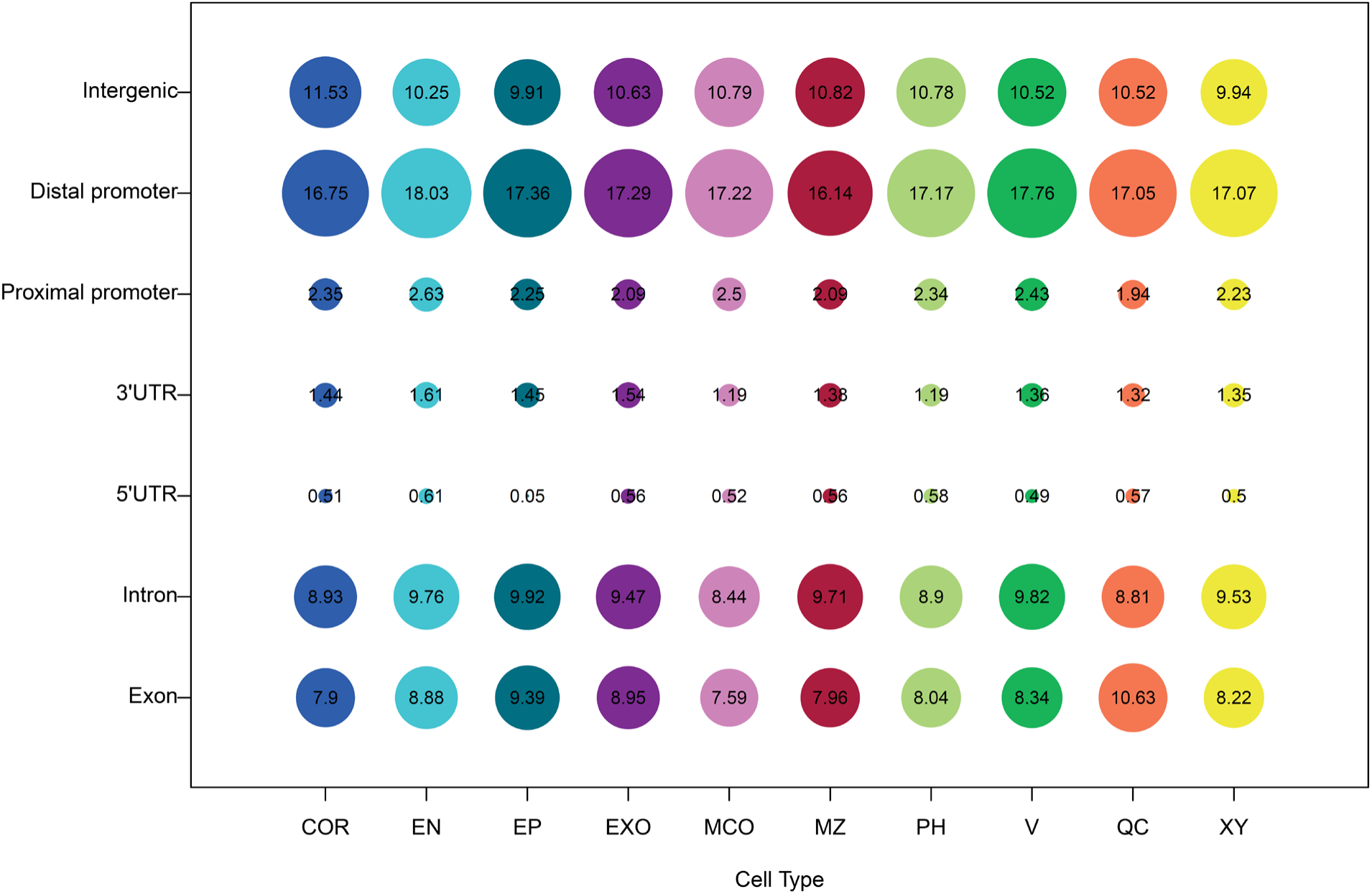
Distribution of annotated genomic regions per cell type. Replicate THSs (repTHSs) were discovered as described in **Materials and Methods.** X-axis: cell type repTHS group. Y-axis: genomic distribution of repTHSs. Proximal promoter = 500 bp upstream of transcription start site. Distal promoter = Between 500 bp and 4kb upstream of the transcription start site. Size of circles is proportional to the percent of repTHSs that are 50% or more overlapping the class of genomic features denoted on the Y-axis. Cell type groups refer to promoters used INTACT lines (**Data S1**): COR=cortex; EN=endodermis; EP=epidermis; EXO=exodermis; MCO=meristematic cortex; MZ=meristematic zone; PH= phloem; V=vasculature; QC=quiescent center; XY= xylem.

**Fig. S28.**
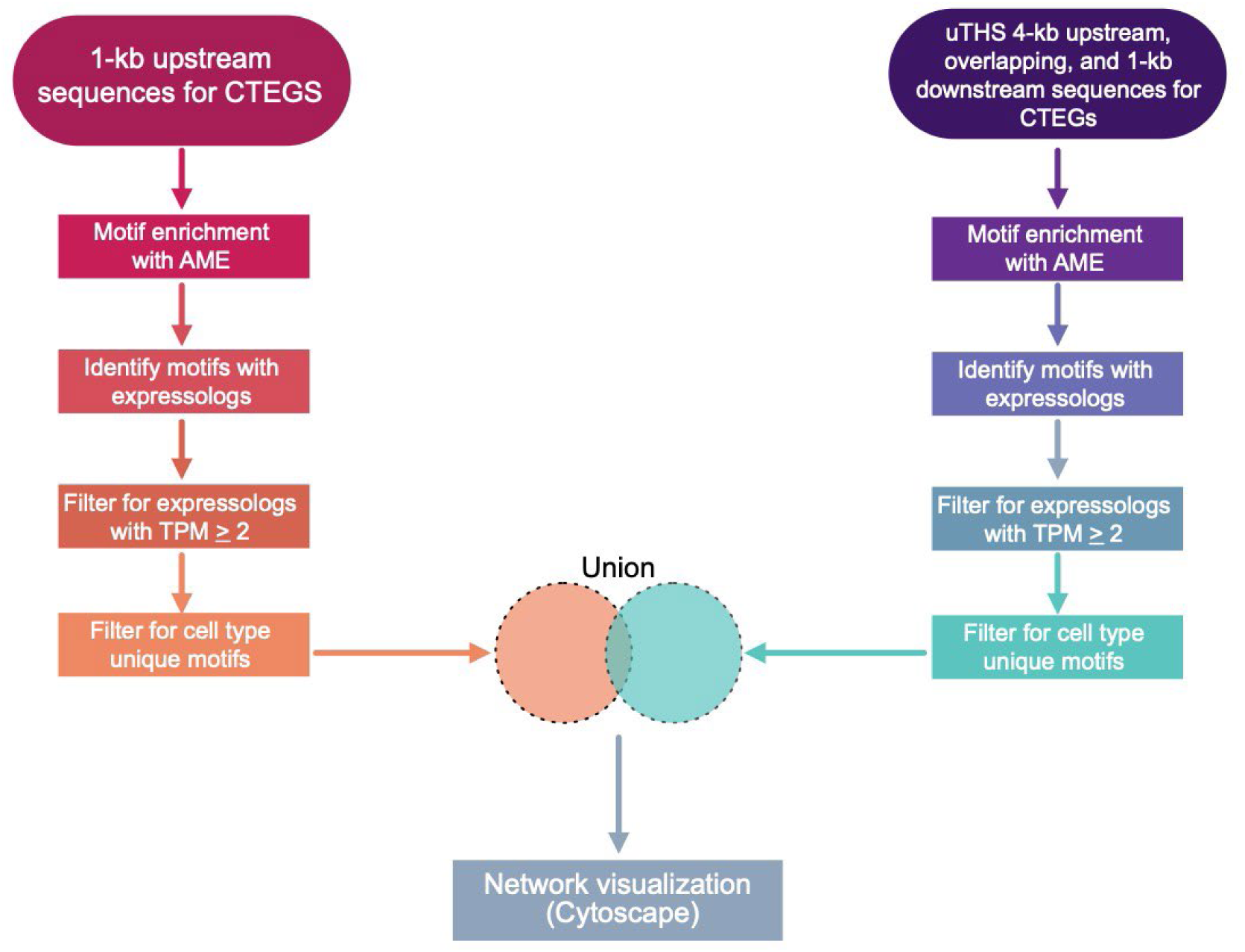
Unique cell type network construction pipeline. Data analysis overview for methods used to construct inferred cell type-unique regulatory networks.

**Fig. S29.**
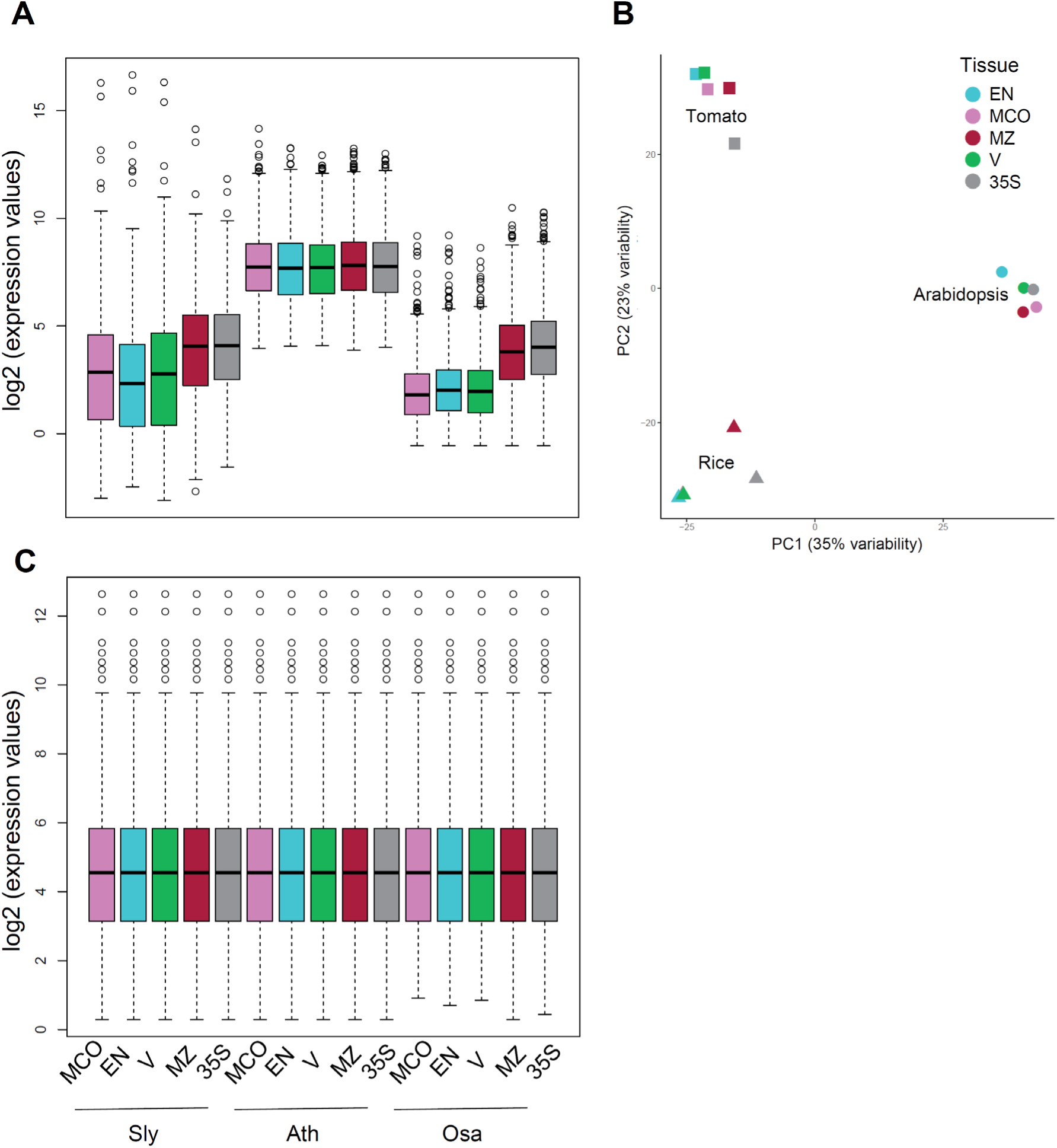
Data demonstrating different dynamic range of translatome data (microarray vs. sequencing) and sample clustering pre-batch effect correction. Boxplots of log_2_ expression values of the homologous cell types and tissues (**A**) before and (**B**) after quantile normalization. Expression data for Arabidopsis are normalized log_2_ intensities and filtered log_2_ counts per million for rice and tomato. Tomato data was also corrected for the differences observed due to differences in sequencing dates. (**C**) Clustering of cell type and tissue expression profiles of 2,642 1:1:1 orthologs between Arabidopsis (circle), rice (triangle) and tomato (square) using principal component (PC) analysis without batch effect correction. At=*Arabidopsis thaliana*; Os=*Oryza sativa*: Sl=*Solanum lycopersicum;* EN=endodermis; MCO=meristematic cortex; MZ=meristematic zone; V=vasculature.

**Fig. S30.**
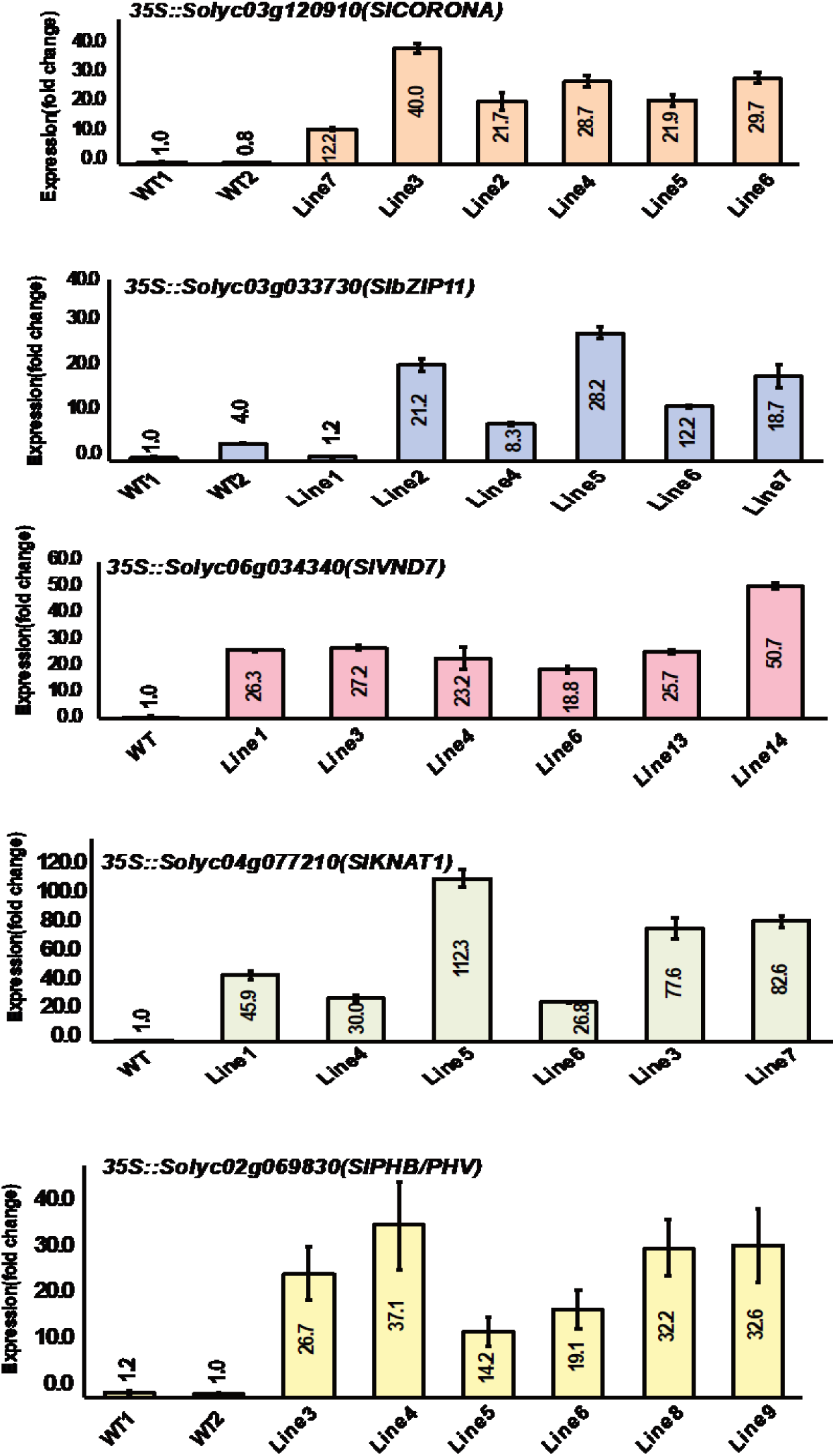
qPCR data for overexpression lines. qRT-PCR measurements for wild type and six independent xylem TF overexpression lines in hairy roots.

**Data S1** - Tomato atlas TRAP-seq promoters, contrasts for differential expression and cell type-enriched genes.

**Data S2** - Gene Ontology and Mapman term enrichment in tomato cell type-enriched genes.

**Data S3** - p-values and odds scores associated with aberrant xylem phenotypes.

**Data S4** - Promoters used in tomato field and pot TRAP-seq data; WGCNA modules and ontology term enrichment.

**Data S5** - Core tomato cell type-enriched genes and ontology term enrichment WGCNA modules and ontology term enrichment.

**Data S6** - ATAC-seq summary data; uTHS regions and dTHS regions are reported in standard BED format. uTHS and dTHS bed files have an additional fourth field denoting region ID. dTHS bed file has an additional fifth field denoting the cell type where the dTHS was found to be enriched.

**Data S7** - Cytoscape network files.

**Data S8** - Multispecies analysis data. Cell type-enriched genes and orthology maps; conserved cell type-enriched genes; within-species cell type/tissue enriched-genes and their overlaps; ontology term enrichment within and between species; constitutively expressed genes and ontology term enrichment across species.

**Data S9** - Fisher’s exact test for overlap of Arabidopsis nitrogen network genes in the tomato exodermis.

**Data S10** - Primers and promoter/reporter sequences.

